# Multiple overlapping SNARE complexes drive endosome maturation in Drosophila nephrocytes

**DOI:** 10.64898/2026.02.03.703479

**Authors:** Dávid Hargitai, Márton Molnár, András Rubics, Iván Bodor, Dóra Baukál, Anikó Nagy, Virág Balogh, Zsófia Simon-Vecsei, Gábor Juhász, Péter Lőrincz

## Abstract

Endosomal maturation determines whether internalized cargo is recycled or degraded, yet the molecular logic governing early endosomal fusion remains poorly defined. This process is often depicted as a linear Rab5-to-Rab7 transition mediated by a single, ordered SNARE pathway, but extensive redundancy in mammalian systems has obscured pathway architecture. Here, using Drosophila nephrocytes as a genetically tractable in vivo model with minimal SNARE redundancy, we show that early endosome maturation is driven by multiple parallel, non-interchangeable SNARE-dependent pathways.

We first resolve a long-standing discrepancy in Syntaxin 7 family orthology, demonstrating that the Drosophila protein previously termed Syx7/Avl is functionally analogous to mammalian STX12 rather than STX7, while late endosomal and lysosomal fusion is mediated by a distinct Syntaxin 7 homolog (Syx13). Based on this reclassification, we define a Syx12L-Snap29-Ykt6 complex that drives canonical homotypic early endosomal fusion. In addition, we identify two related SNARE assemblies - Syx7L-Snap29-Ykt6 and Syx7L-Snap29-Vamp7-that promote later stages of endosomal and lysosomal fusion with distinct Rab GTPase requirements.

These partially compensatory complexes remain active when the canonical pathway is disrupted, producing divergent morphological outcomes, including the formation of aberrant endolysosomal swirls. We establish Snap29 as a central Qbc-SNARE integrating all endosomal fusion routes and uncover a dual role for Ykt6 in promoting maturation while also participating in endosomal recycling.

Together, our findings revise the prevailing model of endosome maturation, revealing a network of parallel, regulated fusion pathways that confer robustness and plasticity to the endolysosomal system.

## INTRODUCTION

Eukaryotic cells internalize transmembrane receptors and extracellular material through endocytosis. Following uptake, cargo is either recycled back to the plasma membrane or delivered to lysosomes for degradation. Early (sorting) endosomes constitute the central decision-making hub of this pathway. These dynamic tubulo-vesicular compartments mature through coordinated membrane remodelling and fusion events that are tightly regulated by phosphatidylinositol-phosphates and small GTPases, including Rab proteins. In their GTP-bound state, Rab proteins recruit effector molecules that control vesicle targeting, tethering, and fusion. Membrane fusion itself is executed by SNARE (Soluble N-ethylmaleimide–sensitive factor Attachment protein REceptor) proteins, whose assembly is promoted by the SNARE chaperone SM-proteins and tethering factors that bridge opposing membranes and stabilize trans-SNARE complexes (*1–5*).

In addition to large coiled-coil tethering proteins such as Early Endosomal Antigen 1 (EEA1) and Rabenosyn-5, endosome maturation relies primarily on two related multisubunit tethering complexes. The HOPS complex promotes fusion of lysosomes with Rab7- and/or Rab2-positive late endosomes, autophagosomes, secretory granules, and Golgi-derived carriers (*6–18*). Its counterpart, CORVET (Class C core vacuole/endosome tethering), mediates tethering and fusion of Rab5-positive early endosomal membranes (*9, 19–27*). HOPS and CORVET share a conserved class C core composed of Vps11, Vps16, Vps18, and Vps33, but differ in their Rab-interacting subunits: CORVET contains the Rab5-binding proteins Vps8 and Vps3 (or Vps39-2/ Tgfbrap1/Trap1 in mammalians), whereas HOPS incorporates the Rab7 adaptor Vps41 and the Rab2-binding subunit Vps39 in yeast (*9, 21, 22, 27–29*). In Drosophila, CORVET is further simplified into a smaller variant termed miniCORVET, composed of Vps8 and only three class C subunits (Dor/Vps18, Car/Vps33A, and Vps16A), rendering Vps11 specific to HOPS in flies (*30, 31*).

SNARE complexes generally consist of three or four SNARE proteins, each contributing one or two SNARE motifs. Within the resulting four-helix bundle, the motifs fall into four classes - Qa, Qb, Qc, and R (*32*). Functional SNARE complexes obey the QabcR rule, incorporating one motif from each class, and therefore usually consist of four different proteins (*32–35*). An important exception is provided by Qbc-SNAREs, which contain two SNARE motifs; in these cases, a single Qa- and a single R-SNARE are sufficient to form a functional complex (*33*). By the late 2000s, a canonical set of SNARE complexes had been proposed to mediate the major fusion steps of endosomal maturation in yeast and mammalian cells. In mammals, the STX12 (Qa)–VTI1A (Qb)–STX6 (Qc)–VAMP4 (R) complex was shown to drive homotypic early endosome fusion (*34*). As endosomes mature, late endosomes form and undergo homotypic fusion or heterotypic fusion with early endosomes via the STX7 (Qa)–VTI1b (Qb)–STX8 (Qc)–VAMP8 (R) complex (*35–37*). Subsequent fusion of late endosomes with lysosomes is mediated by the closely related STX7 (Qa)–VTI1b (Qb)–STX8 (Qc)–VAMP7 (R) complex, which also supports homotypic lysosome fusion (*38, 39*).

More recent work, however, has substantially revised this linear view. As summarized by Dingjan and colleagues, multiple SNARE complexes can often catalyze the same fusion step, with their relative contributions varying across cell types and physiological contexts (*40*). Proteins such as VTI1a and VTI1b, previously considered essential and functionally distinct, can be partially redundant or even dispensable in specific settings (*41–44*). Similar context-dependent redundancy has been proposed for several other SNAREs (*40, 45–48*), underscoring the plasticity and complexity of endosomal fusion in mammalian cells.

In contrast to mammals, SNARE redundancy is markedly lower in Drosophila, which encodes roughly half as many SNARE proteins while retaining clear homologs for most mammalian counterparts (*49–51*). In flies, Syx7 (Qa) (*52, 53*), Snap29 (Qbc) (*54*), Syb (R) (*55*), and nSyb (R) (*55*) have been implicated in early endocytic steps, whereas Syx13 (Qa) (*14, 56, 57*), Syx16 (Qa) (*58*), Syx17 (Qa) (*59, 60*), Syx18 (Qa) (*61*), Sec20 (Qb) (*61*), Snap29 (Qbc) (*54, 59, 60*), Ykt6 (R) (*62*), and Vamp7 (R) (*14, 59, 60*) have been linked to lysosomal fusions and autophagy. Additional roles in recycling and retrograde transport have been described for Syx1 (Qa) (*63*), Syx4 (Qa) (*63*), Syx16 (Qa) (*64*), Ykt6 (R) (*65*), and Syb (R) (*55, 64, 66*). Importantly, we noticed a notable and still unexplained difference in the literature between mammals and Drosophila: Syntaxin7 is consistently described as a late endosomal/lysosomal SNARE in mammalian cells (*38, 67*), yet in flies it has been reported to function primarily at early endosomes (*52, 53*).

Despite the identification of numerous SNAREs associated with endosome maturation, only a limited number of functional SNARE complexes have been rigorously defined. Many studies have focused on individual SNAREs or small subsets of the family, often without biochemical validation of complex formation, leaving key mechanistic questions unresolved. Increasing evidence from multiple model systems indicates that several SNARE assemblies can act in parallel during endosomal maturation, complicating efforts to define linear pathways and obscuring how vesicle fate is determined within this highly interconnected system.

Here, we take advantage of the reduced SNARE redundancy of Drosophila to perform a comprehensive, system-level analysis of SNARE function in the endolysosomal pathway. Using Drosophila nephrocytes as a well-established model of endocytosis (*11, 30, 31*), we identify multiple SNARE complexes that act in parallel during the early stages of endosome maturation. Our results resolve long-standing ambiguities regarding the functional assignment of Syx7-family Qa-SNAREs in flies, leading us to propose a revised nomenclature that reflects both evolutionary history and functional specialization. Based on genetic, biochemical, and ultrastructural analyses, we present an updated model of early endosomal fusion that incorporates pathway branching, differential Rab5/Rab7 dependence, and regulated partial redundancy among SNARE complexes. Together, our findings establish how overlapping SNARE machineries coordinate vesicle fate decisions within the endolysosomal system.

## RESULTS

### Syx7 (Qa), Snap29 (Qbc) and Ykt6 (R) are required for early endosomal fusion in nephrocytes

To systematically assess the contribution of SNARE proteins to endosomal maturation, we performed an RNAi-based screen by targeting 23 of the 25 Drosophila SNARE genes one by one using 77 independent, commercially available RNAi constructs. We compared all phenotypes to a control (luciferase RNAi); Vti1b was excluded due to the lack of available strains, and Syx17 was omitted because we previously showed that its depletion blocks endosome maturation indirectly via a secondary “tethering lock” effect (*68*). As a primary readout, we quantified the size of Rab7–positive endosomes, which sensitively reflects defects along the endocytic pathway (Fig. 1A shows statistics for representative RNAi constructs, and representative micrographs are shown in Figs. S1 and S2). Impaired early endosomal fusion typically results in reduced late endosome size (Fig. 1A-E) (*30*), whereas defects in lysosomal fusion, degradation, or recycling lead to enlarged late endosomes (Fig. 1A) (*11, 30, 31, 69–72*). SNAREs were considered hits when at least two independent RNAi lines produced comparable phenotypes with the same effect on Rab7+ endosome size. Based on increased late endosome size, we identified Syx1 (Fig. S1B), Syx13 (Fig. S1F), Syx16 (Fig. S1G), Syx18 (Fig. S1H), Membrin (Fig. S1J), Sec20 (Fig. S1K), Syb (Fig. S2E) and Vamp7 (Fig. S2G) as SNAREs acting in later stages of endosome maturation and/or recycling in Drosophila. Strikingly, silencing of only three SNAREs - Syx7 (Qa) (Fig. 1C, S1E), Snap29 (Qbc) (Fig. 1D, S2C), and Ykt6 (R) (Fig. 1E, S2H) - consistently resulted in reduced average Rab7-positive endosome size, indicative of defects at earlier stages Notably, the Qa–Qbc–R identities of Syx7, Snap29 and Ykt6 makes them a plausible candidate set for a single functional SNARE complex. While a single RNAi construct against Use1 (Qc) (Fig. S1P) also caused a mild endosome size reduction, the lack of independent supporting evidence led us to exclude it from further analysis.

**Figure 1.**
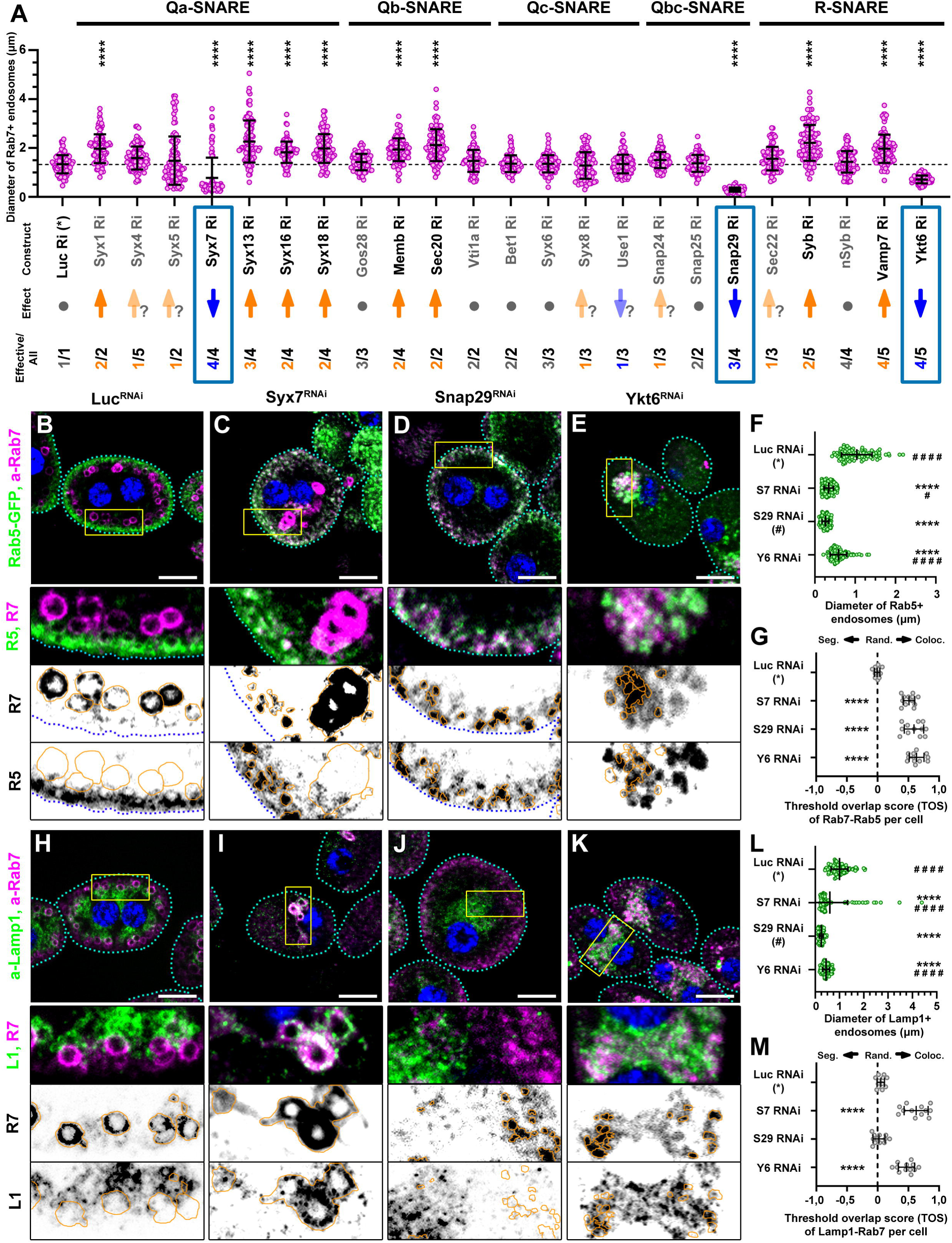
RNAi screen identifies Syx7, Snap29, and Ykt6 as key executors of early endosomal fusion. A) Quantification of Rab7^+^ late endosome size in nephrocytes expressing selected SNARE RNAi constructs. The dashed line indicates the mean endosome size in control (Luciferase RNAi) cells. ****, p < 0.0001. Grey dots indicate no significant change; orange arrows indicate increased size; blue arrows indicate decreased size. Pale arrows denote phenotypes observed in only one independent RNAi line. Fractions below arrows indicate the number of RNAi lines showing a phenotype relative to the total number tested. Representative images from the full screen and complete quantifications are shown in Figures S1 and S2. B–E) Rab5–GFP–positive early endosomes and Rab7^+^ late endosomes are reduced in size in Syx7 RNAi (C), Snap29 RNAi (D), and Ykt6 RNAi (E) nephrocytes compared to control (B). Blue channel shows nuclei (DAPI). Cell outlines are indicated by dashed cyan lines. Scale bars: 10 μm. Rab7^+^ structures are outlined by continuous orange lines in the black-and-white insets. Large Rab7^+^ structures with abnormally thick membranes are apparent in Syx7 RNAi cells (C). F) Quantification of Rab5^+^ endosome size shown in B–E. N = 100 endosomes from 10 cells. Asterisks (*) indicate comparisons to control; green hashes (#) indicate comparisons to Snap29 RNAi. ****, #### p < 0.0001; # p < 0.05. Snap29 depletion results in the smallest early endosomes. G) Quantification of Rab5–Rab7 colocalization in B–E using the Threshold Overlap Score (TOS) reveals increased overlap upon depletion of all three SNAREs. ****, p < 0.0001. H–K) Rab7^+^ late endosomes and Lamp1^+^ lysosomes are smaller in Syx7 RNAi (I), Snap29 RNAi (J), and Ykt6 RNAi (K) nephrocytes compared to control (H). Blue channel shows nuclei (DAPI). Cell outlines are indicated by dashed cyan lines. Scale bars: 10 μm. Rab7^+^ structures are outlined by continuous orange lines in the black-and-white insets. The large Rab7^+^ structures observed in Syx7 RNAi cells are Lamp1-positive (I). L) Quantification of Lamp1^+^ lysosome size shown in H–K. N = 100 lysosomes from 10 cells. Asterisks (*) indicate comparisons to control; green hashes (#) indicate comparisons to Snap29 RNAi. ****, #### p < 0.0001. Snap29 depletion results in the smallest lysosomes. M) Quantification of Rab7–Lamp1 colocalization using TOS reveals increased overlap in Syx7- and Ykt6-depleted cells. ****, p < 0.0001.

To characterize Syx7, Snap29 and Ykt6 RNAi phenotypes in more detail, we analyzed early endosomes and lysosomes using Rab5-GFP (Fig. 1B-G) and anti-Lamp1 (Fig. 1H-M) staining, respectively, in combination with anti-Rab7 labeling. In all three knockdowns, Rab5-positive early endosomes were significantly smaller (Fig. 1F), supporting a role for Syx7, Snap29, and Ykt6 in early endosomal fusions. Snap29 depletion caused the most pronounced reduction in early endosome size. Moreover, colocalization between Rab5-GFP and anti-Rab7, quantified using the Threshold Overlap Score (TOS) method (Fig. 1G), was increased in all three conditions, indicating impaired endosomal maturation. According to the literature [0,1; 1] TOS suggests colocalization,]-0,1;0,1[ means close to random overlap, while [-1; - 0,1] TOS argues for segregation of the signals (*73*). Late endosomes were also affected in all three knockdowns, although with distinct phenotypic signatures. Snap29 silencing resulted in uniformly small Rab7-positive compartments (Fig. 1D), mirroring the early endosome phenotype. In contrast, Syx7 knockdown (Fig. 1C) reduced average late endosome size but additionally produced a subset of unusually large Rab5-GFP negative, Rab7-positive compartments with surprisingly thick Rab7 signal—a phenotype not observed in any other single SNARE depletion. Ykt6 silencing (Fig. 1E) generated smaller-than-control Rab7-positive endosomes with markedly increased Rab5-GFP colocalization.

Lysosomes were similarly affected: anti-Lamp1–positive compartments were smaller in all three knockdowns than in the control (Fig. 1H). In Syx7-depleted cells (Fig. 1I), Lamp1 strongly colocalized with Rab7 on the above-mentioned large aberrant endolysosomal structures. In contrast, Snap29 (Fig. 1J) silencing did not increase Lamp1–Rab7 colocalization relative to controls, whereas Ykt6 depletion (Fig. 1K) resulted in a population of Rab7/Lamp1 double-positive endolysosomes.

### Silencing of Snap29, Syx7, or Ykt6 impairs endocytic uptake

Our morphological analyses suggested that Snap29, Syx7, and Ykt6 function at early stages of endosome maturation. To determine whether these defects also compromise endocytic activity, we performed tracer uptake assays in nephrocytes. As an initial screen, we monitored short-term (5 min) uptake of FITC–Avidin following RNAi-mediated silencing of individual SNARE genes using constructs selected based on the results of our a-Rab7 screen (Fig. 2A). While depletion of Syx1 (Fig. 2A, S3B), Syx5 (Fig. 2A, S3D), or Use1 (Fig. 2A, S3P) caused a moderate reduction in tracer internalization, silencing of Syx7 (Fig. 2A, S3E), Snap29 (Fig. 2A, S3S), or Ykt6 (Fig. 2A, S3X) completely abolished detectable FITC–Avidin uptake (Fig. 2A). This result indicated a severe disruption of endocytosis, in line with our endosome size and colocalization analyses. Garland nephrocytes internalize material through deep plasma membrane invaginations termed lacunae, which are lined by a slit diaphragm that is structurally and functionally analogous to that of mammalian podocytes. This filtration barrier is essential for nephrocyte detoxification and hemolymph sieving, and it has a size exclusion limit close to that of FITC–Avidin (66 kDa). Previous studies have shown that impaired trafficking of slit diaphragm components can lead to “clogging” at the lacunar entrance, reducing the effective exclusion limit and thereby blocking uptake of larger tracers (Fig. 2B) (*74*).

**Figure 2.**
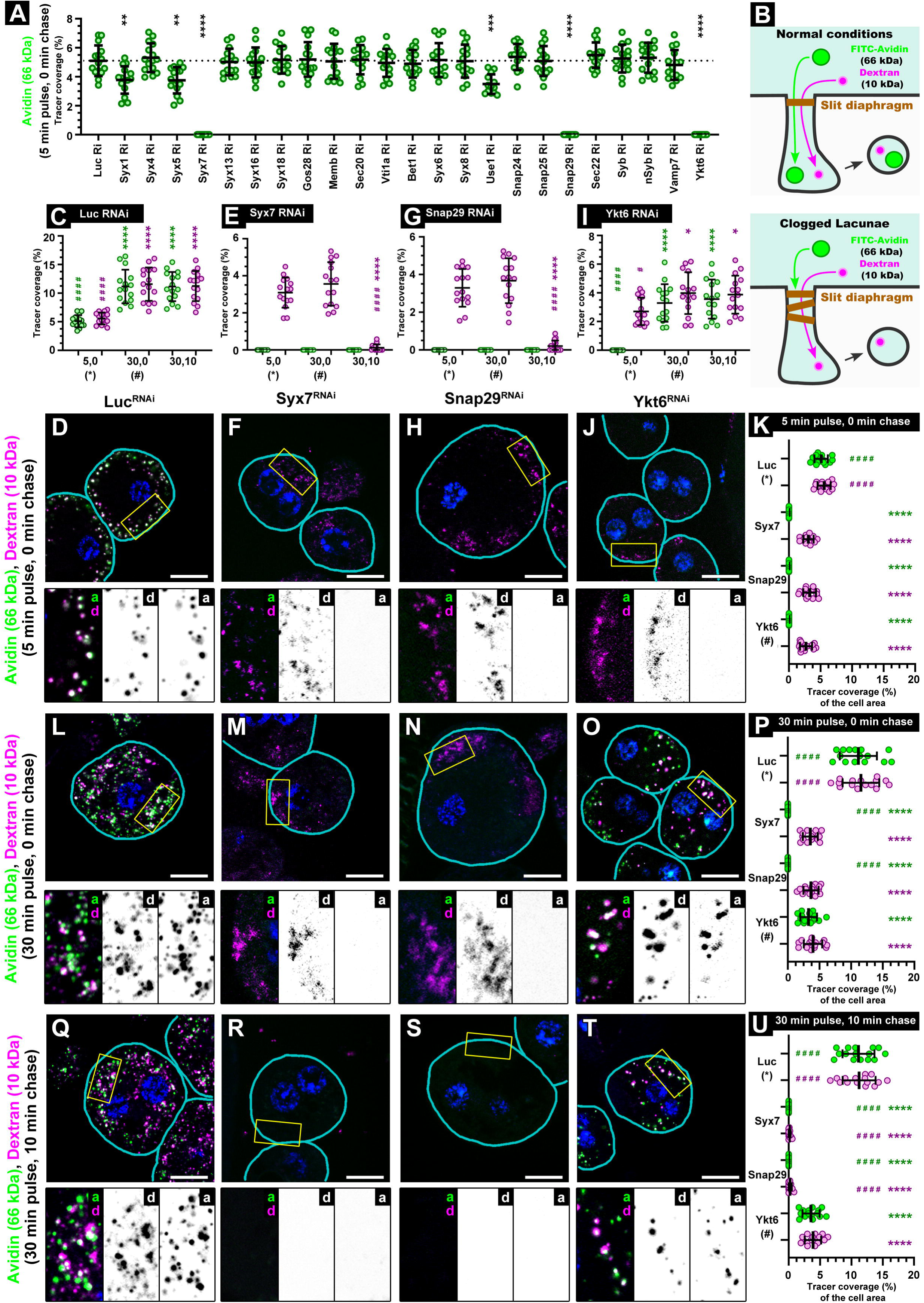
Silencing of Syx7, Snap29, or Ykt6 impairs endocytic uptake. A) Quantification of Avidin uptake across representative SNARE RNAi conditions. Dashed line indicates mean tracer coverage in control cells. **** *p* < 0.0001; *** *p* < 0.001; ** *p* < 0.01. Representative images are shown in Figure S3A–X. B) Schematic illustrating lacunar clogging as a potential mechanism underlying impaired uptake in the hits identified by the Avidin uptake screen. C-U) Nephrocytes were pulsed with fluorescent Avidin (66 kDa) and Dextran (10 kDa) for 5 min (D, F, H, J, K) or 30 min (L-U), then fixed immediately (D, F, H, J, K, L-P) or chased for 10 min before fixation (Q–U). Blue channel shows nuclei (DAPI); cell outlines are indicated by cyan lines. Scale bars: 10 μm. In control cells (D, L), both tracers rapidly accumulated in large peripheral endosomes and subsequently in deeper compartments, consistent with lysosomal delivery. In Syx7 (F, M) and Snap29 (H, N) RNAi cells, Avidin uptake was nearly absent, and Dextran signal was strongly reduced and rapidly cleared during chase (R, S), suggesting impaired uptake and enhanced clearance, potentially via recycling. In Ykt6 RNAi cells (J), Avidin uptake was also minimal and Dextran uptake reduced, but internalized cargo became detectable after prolonged pulse (O) and persisted during chase (T), indicating slower uptake and impaired clearance. C, E, G, I) Quantification of internalized tracer signal coverage (% of cell area) in different RNAi backgrounds. C) Control RNAi (D, L,), E) Syx7 RNAi (F, M, R), G) Snap29 RNAi (H, N, S), I) Ykt6 RNAi (J, O, T). Asterisks (*) indicate comparisons to 5 min pulse, no chase; green hashes (#) indicate comparisons to 30 min pulse, no chase. ****, #### *p* < 0.0001, * p < 0.05. K, P, U) Quantification of internalized tracer signal coverage (% of cell area) during different experimental settings, corresponding to D-J, L-O, and Q-T, respectively. Asterisks (*) indicate comparisons to control; green hashes (#) indicate comparisons to Ykt6 RNAi. ****, #### *p* < 0.0001.

To test whether such clogging contributes to the observed phenotype, we compared uptake of FITC–Avidin and a much smaller fluorescent dextran (10 kDa) following 5- and 30-minute incubations. In control nephrocytes, both tracers were efficiently internalized, with intracellular signal increasing over time (Fig. 2C, D, L). In contrast, Syx7- (Fig. 2E, F, K) or Snap29-depleted cells (Fig. 2G, H, K) failed to internalize FITC–Avidin, while dextran uptake was reduced relative to controls (Fig. 2F, H, K) and did not increase with prolonged incubation (Fig. 2M, N, P). These findings are consistent with clogging of the slit diaphragm, although the time-independent dextran levels suggested an additional defect. The stable dextran signal raised the possibility that internalized material might be rapidly removed from the cell via recycling. We therefore tested whether impaired endosome maturation promotes accelerated recycling to the plasma membrane. Nephrocytes were incubated with both tracers for 30 minutes, followed by a 10-minute chase in tracer-free medium (Fig. 2Q-U). In control cells, tracer levels remained unchanged (Fig. 2Q, U). In contrast, Syx7- (Fig. 2R, U) or Snap29-depleted cells (Fig. 2S, U) were almost completely devoid of dextran after the chase, supporting the notion of rapid recycling in these conditions.

Ykt6 depletion produced a distinct phenotype. After 5 minutes (Fig. 2I, J, K), uptake resembled that observed upon Syx7 or Snap29 silencing (Fig. 2F, H). However, after 30 minutes, FITC–Avidin was clearly internalized and dextran levels increased compared to the short incubation (Fig. 2I, O, P). This indicates that loss of Ykt6 does not disrupt slit diaphragm sieving but instead reduces overall endocytic efficiency. Consistently, tracer retention following the 10 min chase (Fig. 2I, T, U) was comparable to that after continuous incubation (Fig. 2O), indicating that Ykt6 silencing does not trigger rapid recycling. To further corroborate these findings, we performed a silver nitrate uptake assay by rearing larvae on silver nitrate–containing food, in which silver accumulates and precipitates in the lysosomes of nephrocytes. In all three SNARE knockdowns, but most profoundly in Syx7 and Snap29, silver accumulation was reduced relative to controls (Fig. S3Y) and mirrored the altered lysosomal patterns observed by anti-Lamp1 staining (Fig. 1H-K).

Together, these uptake assays demonstrate that Syx7, Snap29, and Ykt6 are required for efficient early endocytic function, although the loss of each SNARE affects the pathway in a distinct manner. To determine whether these proteins act together within the same molecular machinery, we next examined their localization and physical interactions.

### Snap29, Syx7, and Ykt6 localize to Rab5-positive endosomes and form a SNARE complex

To determine the subcellular localization of the three candidate SNAREs, we analyzed garland nephrocytes expressing Rab5–GFP (Fig. 3A-C) or stained for Rab7 (Fig. 3D-F). Syx7 (Fig. 3A, D, H) and Snap29 (Fig. 3B, E, H, I) were detected using specific antibodies, while Ykt6 localization was assessed using a Ykt6–HA transgene (Fig. 3C, F, H, I). All three SNAREs showed strong colocalization with Rab5-positive early endosomes (Fig. 3A-C) and minimal overlap with Rab7-positive compartments (Figure 3D-F), as quantified by Threshold Overlap Score (TOS) (Fig. 3G). Notably, in control cells the TOS values between Rab5–GFP and Rab7 were close to zero similar to the SNARE and Rab7 pairs (Fig. 1G), suggesting that Syx7, Snap29, and Ykt6 still be transiently present on Rab5/Rab7 double-positive endosomes and are removed when Rab7 conversion occurs.

**Figure 3.**
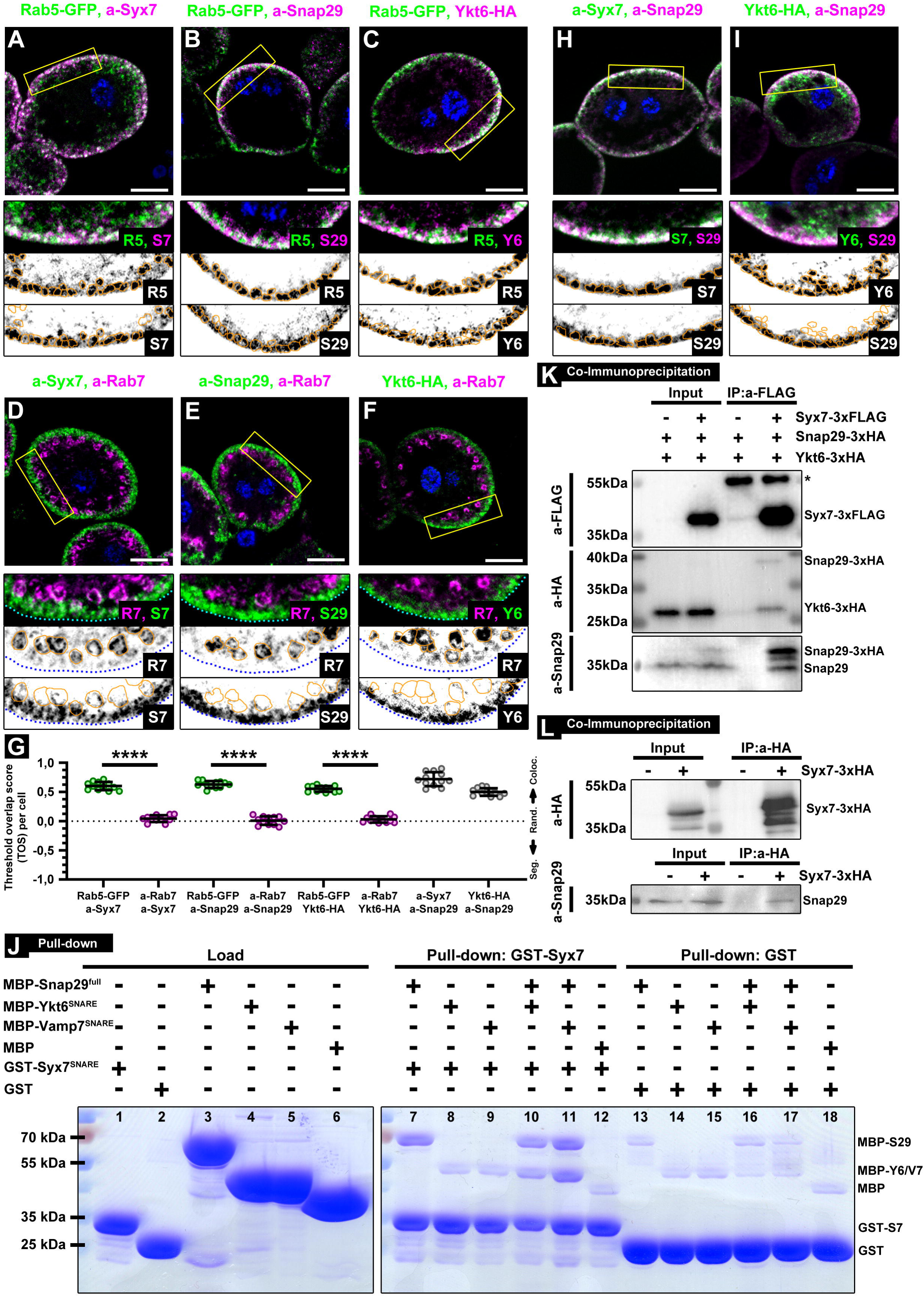
Syx7, Snap29, and Ykt6 localize to early endosomes and form a SNARE complex. A–I) Colocalization analyses of Syx7, Snap29, and Ykt6 with endosomal markers and with each other in nephrocytes. Blue channel shows nuclei (DAPI); cell outlines are indicated by dashed cyan lines. Scale bars: 10 μm. A–C) Endogenous Syx7 (A), endogenous Snap29 (B), and Ykt6–HA (C) localize predominantly to Rab5–GFP^+^ early endosomes. Rab5-GFP^+^ structures are outlined by continuous orange lines in the black-and-white insets. D–F) Minimal overlap is observed between Syx7 (F), Snap29 (G), or Ykt6–HA (H) and Rab7^+^ late endosomes. Rab7^+^ structures are outlined by continuous orange lines in the black-and-white insets. H, I) Endogenous Snap29 colocalizes with endogenous Syx7 (D) and Ykt6–HA (E). Syx7^+^ (H) and Ykt6^+^ (I) structures are outlined by continuous orange lines in the black-and-white insets. G) Quantification of colocalization using Threshold Overlap Score (TOS). SNAREs show significantly higher colocalization with Rab5^+^ endosomes than with Rab7^+^ compartments. TOS also confirms colocalization between Snap29 and Syx7 or Ykt6. **** p < 0.0001. J) In vitro reconstitution of SNARE interactions. Lanes 1–6: purified recombinant proteins. Lane 7: Syx7 SNARE domain pulls down full-length MBP–Snap29. Lanes 8–9: Syx7 SNARE domain does not bind the SNARE domains of Ykt6 or Vamp7 alone. Lanes 10–11: Syx7 SNARE domain binds Snap29 together with the SNARE domains of Ykt6 or Vamp7 when all three are present. Lane 12: Syx7 SNARE domain shows negligible binding to MBP alone. Lanes 13–18: GST control shows no relevant binding. (K) Co-immunoprecipitation from Drosophila S2R^+^ cells shows that Syx7–3xFLAG precipitates Snap29–3xHA, Ykt6–3xHA, and endogenous Snap29. L) Syx7–3xHA precipitates endogenous Snap29 in cultured Drosophila S2R+ cells.

We next asked whether these SNAREs reside on the same vesicles. Double immunostaining for Syx7 and Snap29 (Fig. 3H), as well as for Snap29 and Ykt6–HA (Fig. 3I), revealed substantial colocalization in both cases (Fig. 3G). The slightly lower Threshold Overlap Score (TOS) observed for Snap29–Ykt6 (Fig. 3G) is consistent with the formation of a Qa/Qbc precomplex prior to R-SNARE engagement and further supports that Ykt6 may also exert SNAP29-independent functions on endosomal membranes. The strong functional similarity at early endosomes and their overlapping localization suggested that Syx7, Snap29, and Ykt6 act together as a SNARE complex. To test this, we performed biochemical interaction assays. In GST pull-down experiments (Fig. 3J), bacterially expressed GST–Syx7 SNARE motif efficiently bound full-length MBP–Snap29. In contrast, MBP–Ykt6 bound to GST–Syx7 only in the presence of MBP–Snap29, indicating that Syx7 engages R-SNAREs via the Qbc-SNARE Snap29. Because previous work has implicated Vamp7 as an alternative endosomal R-SNARE in Snap29-containing complexes (*14, 59, 75*), we tested whether the Syx7–Snap29 precomplex could also engage Vamp7. Indeed, the closely related MBP–Vamp7 SNARE motif showed the same Snap29-dependent binding behavior as Ykt6, whereas control GST and MBP proteins showed negligible binding.

These experiments demonstrate three key points: (i) Syx7, Snap29, and Ykt6 can assemble into a SNARE complex without intrinsic biochemical constraints at the SNARE-motif level; (ii) the Qa-SNARE Syx7 engages R-SNAREs only through the Qbc-SNARE Snap29; and (iii) the Syx7–Snap29 Q-SNARE precomplex can interact with more than one type of R-SNARE, including both Ykt6 and Vamp7.

Because in vitro pull-down assays do not capture the full regulatory context of SNARE assembly (*76*), we next tested complex formation in cultured fly cells. Using Drosophila S2R+ cells expressing tagged proteins, co-immunoprecipitation experiments (Fig. 3K) showed that Syx7–FLAG efficiently precipitated both Ykt6–HA and Snap29–HA. Importantly, Syx7–FLAG also co-precipitated endogenous Snap29. This interaction was further confirmed by co-immunoprecipitation from lysates expressing only Syx7–HA (Fig. 3L), which likewise precipitated endogenous Snap29.

Together, our localization and biochemical data demonstrate that Syx7, Snap29, and Ykt6 form a functional SNARE complex acting at Rab5 positive early endosomes.

### Drosophila Syx7/Avl (CG5081) is functionally analogous to mammalian STX12, whereas Syx13 (CG11278) corresponds to STX7

Comparison of our findings with the mammalian literature revealed an apparent inconsistency: in mammalian cells, STX7 functions at late endosomes and lysosomes, whereas in Drosophila Syx7 localizes to early endosomes and is required for early endosomal fusion (*52, 77*). Given the generally conserved roles of SNARE proteins, we sought to resolve this discrepancy. *In silico* analyses of SNARE evolution indicate that the endosomal Syntaxin7 family comprised two paralogs (Syx7′a and Syx7′b) in the last common metazoan ancestor (LCMA; Fig. 4A) (*49*). *Drosophila* retained both genes: Avl/Syx7 (derived from Syx7′a) and Syx13/Syx20 (derived from Syx7′b). In vertebrates, Syx7′a underwent duplication, giving rise to STX7 and STX12/STX13, while Syx7′b (named TSNARE1 or STX20 in human) was disrupted by insertion of a Harbinger transposon (*78*), resulting in transformation of its SNARE function into a regulatory one (*79*).

**Figure 4.**
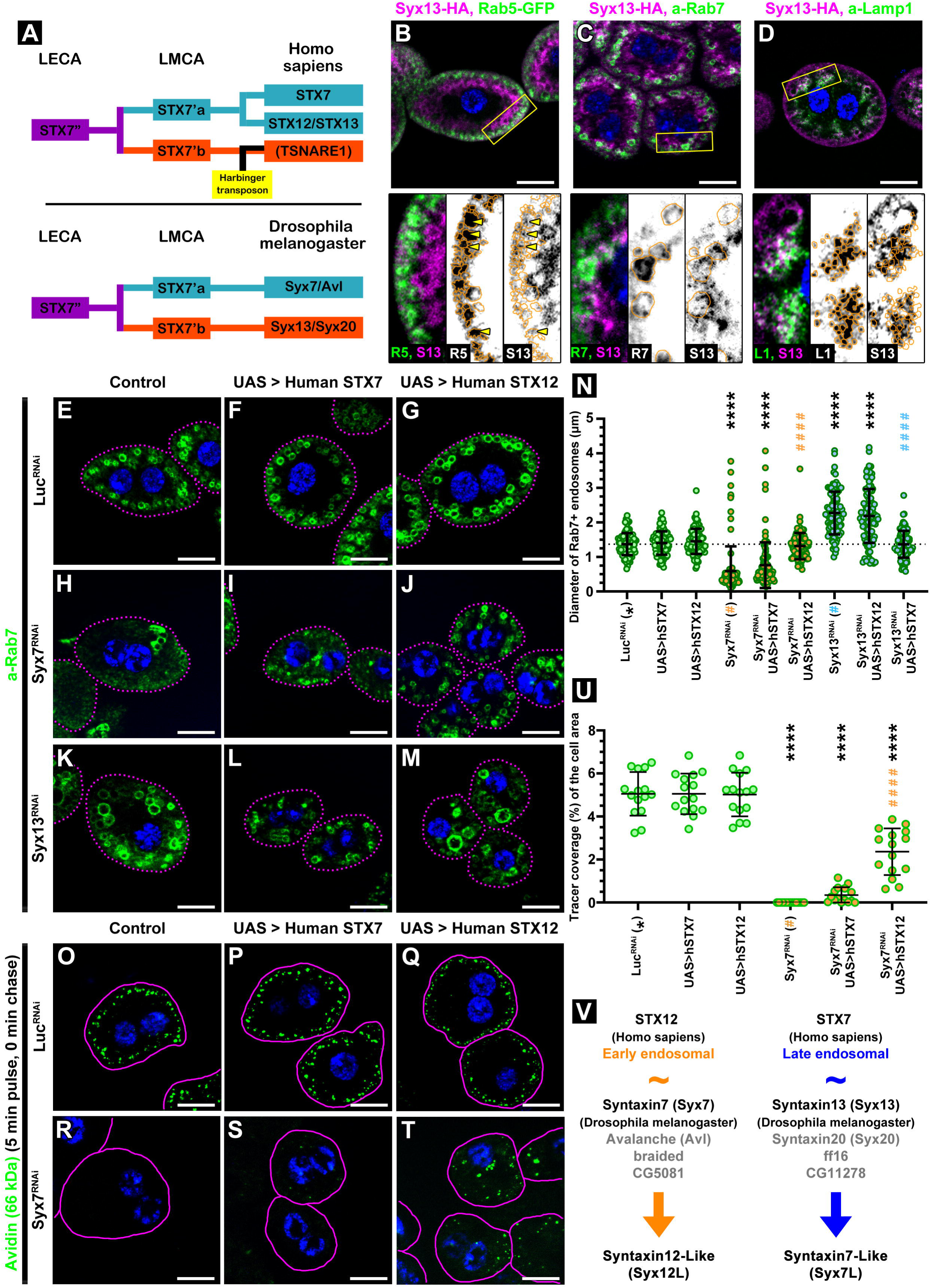
Functional reassignment of Syntaxin7 family members in Drosophila. A) Schematic depicting the evolutionary relationships of Syntaxin7 family members in humans and Drosophila. LECA: Last Eukaryotic Common Ancestor, LMCA: Last Metazoan Common Ancestor C–D) Syx13-HA localizes to Rab7^+^ late endosomes (C) and Lamp1^+^ lysosomes (D) in nephrocytes. Only minimal but detectable colocalization is observed with Rab5^+^ early endosomes (B, yellow arrowheads). Blue: nuclei (DAPI). Rab5-GFP^+^ (B), Rab7^+^ (C) and Lamp1^+^ (D) structures are outlined by continuous orange lines in the black-and-white insets. Scale bars: 10 μm. E–U) Rescue experiments using human STX7 and STX12 transgenes in Drosophila nephrocytes. E–M, O–T) Blue: nuclei (DAPI). Cell outlines are indicated by magenta lines. Scale bars: 10 μm. E–G) Rab7^+^ endosomes appear normal in control (E), hSTX7-expressing (F), and hSTX12-expressing (G) cells. H-J) Rab7^+^ endosomes are significantly smaller in Syx7 RNAi cells (H); this phenotype is not rescued by hSTX7 (I) but is partially rescued by hSTX12 (J). K–M) Rab7^+^ endosomes are enlarged in Syx13 RNAi cells (K); this phenotype is not rescued by hSTX12 (M) but is partially rescued by hSTX7 (L). N) Quantification of Rab7^+^ endosome size shown in E–M. N = 100 endosomes from 10 cells. Asterisks (*) indicate comparisons to control; orange and cyan hashes (#) indicate comparisons to Syx7 and Syx13 RNAi cells, respectively. ****, #### p < 0.0001. O–T) Avidin uptake and transport to endosomes proceed normally in control (O), hSTX7 expressing (P), and hSTX12 expressing (Q) cells. In Syx7 RNAi cells (R), avidin uptake is nearly abolished; this defect is not rescued by hSTX7 (S) but is partially rescued by hSTX12 (T). Blue: nuclei (DAPI). Cell outlines are indicated by dashed cyan lines. Scale bars: 10 μm. U) Quantification of avidin uptake shown in O–T. Asterisks (*) indicate comparisons to control; orange hashes (#) indicate comparisons to Syx7 RNAi. ****, #### p < 0.0001. V) Schematic of the updated nomenclature of Drosophila Syntaxin 7 family members: Syx7 is renamed Syx12L and Syx13 is renamed Syx7L.

Because of the similarity in nomenclature, one might infer that Drosophila Syx7 corresponds to human STX7; similarly, at least one study inferred that Drosophila Syx13 corresponds to STX12/13 (*57*). However, phylogenetic analysis indicates a different relationship: Drosophila Syx13 represents the ancestral Syx7′b lineage whose SNARE function was disrupted in mammals (*78, 79*), whereas Drosophila Syx7 corresponds to the Syx7′a lineage that duplicated in vertebrates (*49, 50*). In support of this assignment, Syx7 localizes predominantly to Rab5-positive early endosomes in Drosophila, whereas Syx13–HA is enriched on Rab7-positive late endosomes and Lamp1-positive lysosomes, with only minimal overlap with Rab5, indicating a primarily late endosomal function (Fig. 4B–D). To directly test functional equivalence, we expressed human STX7–HA or STX12–HA in Syx7- or Syx13-depleted nephrocytes (Fig. 4E-N). Expression of either human SNARE in control cells had no significant effect on late endosome size (Fig. 4E-G). In Syx7-silenced cells, human STX12–HA (Fig. 4J) - but not STX7–HA (Fig. 4I) - rescued the endosomal phenotype (Fig. 4H, N). Conversely, in Syx13-silenced cells (Fig. 4K), STX7–HA (Fig. 4L) - but not STX12–HA (Fig. 4M) - restored normal late endosome size (Fig. 4N). Functional equivalence was further supported by uptake assays: the near-complete loss of FITC–Avidin internalization caused by Syx7 silencing (Fig. 4O, R, U) was rescued by STX12–HA (Fig. 4Q, T, U) but not by STX7–HA (Fig. 4P, S, U).

Together, our experimental data, evolutionary analysis, and the existing literature indicate that functional orthology between Drosophila and mammalian Syntaxin7 family members has shifted through paralogous compensation. In Drosophila, Syx7 retains early endosomal function, while Syx13 acts at late endosomes; in mammals, these roles are partitioned between STX12 and STX7, respectively, most likely because tSNARE1 became a regulatory SNARE (*78, 79*). Since the current nomenclature of Drosophila Syx7 and Syx13 reflects neither their evolutionary history nor their functional equivalence to mammalian proteins, we propose a revised naming scheme (Fig. 4V). We suggest renaming Drosophila Syx7 as Syntaxin12-like (Syx12L) and Syx13 as Syntaxin7-like (Syx7L), where “-like” denotes functional analogy while allowing for species-specific divergence. Hereafter, we adopt this revised nomenclature throughout the manuscript.

### Disruption of early endosome fusion enhances recycling in a Ykt6- and Exocyst-dependent manner

Fluorescence microscopy revealed clear but previously unexplained phenotypic differences among knockdowns of the three SNAREs. To resolve these differences at early stages of endosome maturation, we examined the nephrocyte periphery by electron microscopy using tannic acid impregnation to visualize lacunae and clathrin-coated pits (Fig. 5A-D). Control nephrocytes (Fig. 5A) displayed well-developed lacunae with abundant clathrin-coated pits and large endosomes. In contrast, Syx12L silencing (Fig. 5B) caused a marked deepening of lacunae accompanied by expanded luminal spaces and a pronounced reduction in endosome size. Snap29 depletion (Fig. 5C) also increased lacunar depth, and endosomes were small. Notably, Snap29-deficient lacunae frequently contained intracellular cargo, consistent with our previous finding that Snap29 loss promotes autophagosome secretion in nephrocytes (Fig. S3Z) (*68*). In sharp contrast, Ykt6 silencing (Fig. 5D) led to a drastic shallowing of lacunae, while clathrin-coated pits and medium-sized endosomes were preserved, which fits with the result of our uptake experiments.

**Figure 5.**
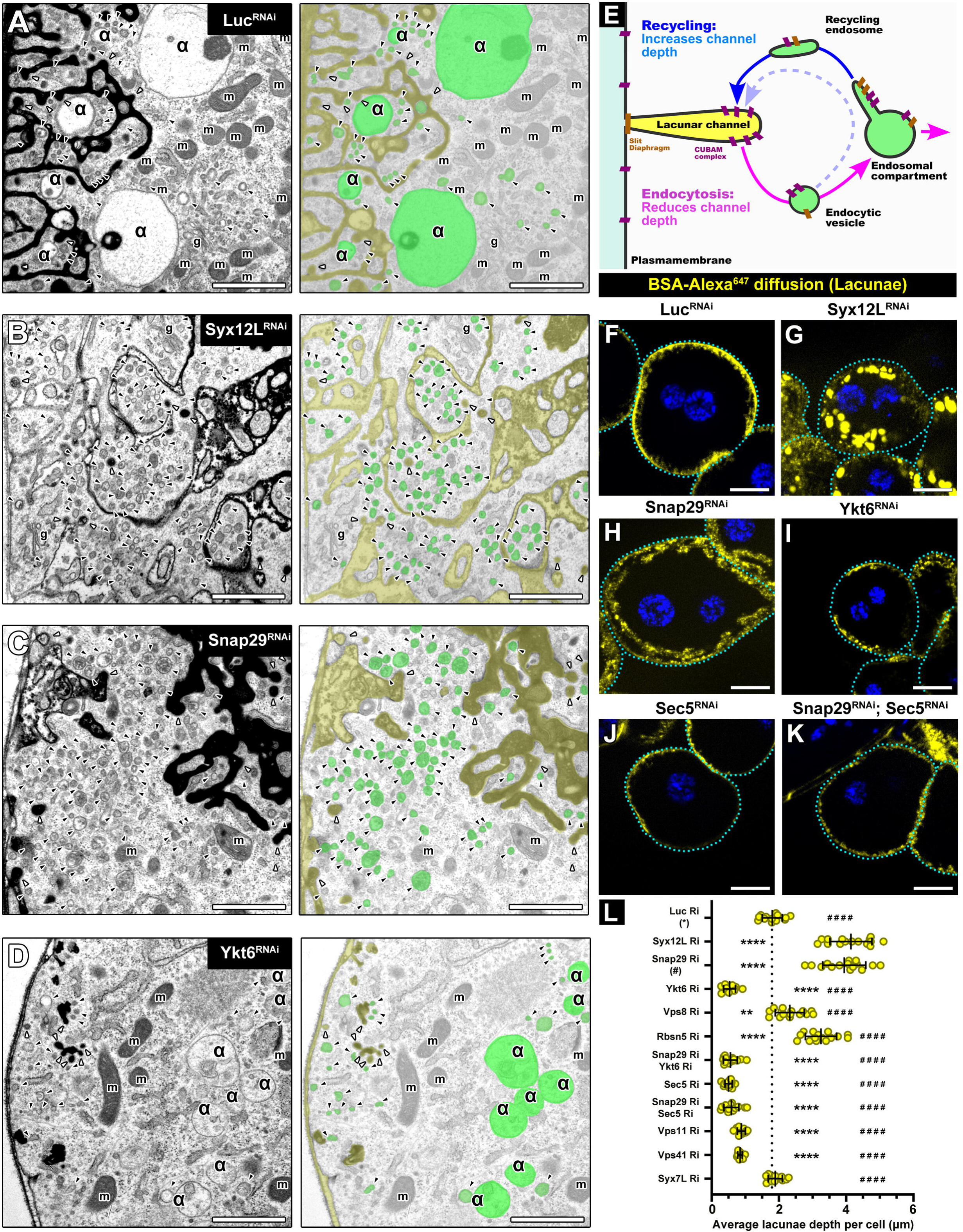
Lacunar depth is increased in Syx12L and Snap29 RNAi cells but is decreased in Ykt6 depleted cells. A-D) Transmission electron micrographs of the peripheral regions of tannic acid (TA)–impregnated nephrocytes. TA fills the lacunar system and appears as dark electron-dense material in the lacunar lumen (colored orange in false-color images). Scale bars: 1 μm. m, mitochondria; α, late endosomes (colored green). White arrowheads indicate TA-filled clathrin-coated pits; black arrowheads indicate clathrin-coated and uncoated early endocytic vesicles and small early endosomes. A) Lacunar depth is normal in control cells, with large late endosomes present. B–C) Lacunar depth is markedly increased in Syx12L (B) and Snap29 (C) RNAi cells. These cells lack normally sized endosomes and instead contain numerous small endosomes. Note that lacunae are dilated in Syx12L RNAi cells (B), whereas the lacunar lumen contains secreted autophagosomes in Snap29 RNAi cells (C; higher magnification in Fig. S3Y). D) Ykt6 RNAi cells lack deep lacunae, and endosomes are smaller than in controls but larger than in Syx12L (B) or Snap29 (C) RNAi cells. E) Schematic illustrating the relationship between lacunar depth and endosomal recycling versus degradation. CUBAM: cubilin-amnionless complex F–K) Channel diffusion assay using fluorescent BSA shows increased lacunar channel depth in Syx12L (G) and Snap29 (H) RNAi cells compared with controls (F). In contrast, lacunar channels are shallower in Ykt6 RNAi cells (I), similar to Sec5 single (J) and Sec5/Snap29 double RNAi (K) cells. Note the intense internal BSA signal in Syx12L cells (G), corresponding to lacunar dilations seen in (B). Blue channel shows nuclei (DAPI); cell outlines are indicated by dashed cyan lines. Scale bars: 10 μm. L) Quantification of lacunar channel depth from (F–K and Fig. S4 A1–A6). The dotted line indicates the mean lacunar depth in control cells. Asterisks (*) indicate comparisons to control; hashes (#) indicate comparisons to Snap29 RNAi. ****, #### p < 0.0001; ** *p* < 0.01.

The ultrastructural differences in lacunar depth pointed to altered recycling dynamics. In nephrocytes, endocytosis occurs primarily at the lacunar membrane, and membrane homeostasis depends on the balance between endocytosis-driven membrane removal and recycling-mediated membrane return (Fig. 5E). Accordingly, enhanced recycling would be expected to deepen lacunae, whereas impaired recycling would decrease their depth. To quantify recycling activity and strengthen our EM data, we used a fluorescent channel diffusion assay to label the lacunar lumen (Fig. 5F-L, S4A). Consistent with the EM data, Syx12L (Fig. 5G) or Snap29 (Fig. 5H) silencing significantly increased average channel depth, albeit with distinct morphologies. In contrast, Ykt6 depletion (Fig. 5I) reduced channel depth, with channels absent from large portions of the cell surface.

Because early endosomal fusion is impaired upon Syx12L or Snap29 depletion, yet based on our uptake experiments endocytosis remains rapid, we hypothesized that fusion-defective endocytic vesicles are rerouted into the recycling pathway. Supporting this idea, silencing of the early endosomal tethering factors Rbsn5 (Fig. 5L, S4A1) or Vps8 (Fig. 5L, S4A2) similarly increased channel depth, whereas depletion of Vps11 (Fig. 5L, S4A3) or Vps41 (Fig. 5L, S4A4) subunits of late endosomal HOPS tethering complex resulted in shallower channels. In turn, silencing of late endosomal Syx7L (Fig. 5L, S4A5) resulted in channel depth similar to control. Thus, enhanced recycling is specifically triggered by disruption of early - but not late -endosomal fusions. Ykt6 depletion (Fig. 5I, L), however, produced the opposite phenotype, suggesting an additional role in recycling. Consistent with this, Ykt6 silencing was epistatic to Snap29 depletion (Fig. 5L, S4A6) with respect to channel depth, mirroring the behavior of the Sec5 subunit of the recycling tether Exocyst (*80*) (Fig. 5J, K, L). This suggests that Ykt6 plays an important role in membrane recycling.

To further assess the balance of endocytosis and recycling we studied a physiologically critical cargo class of nephrocytes, the slit diaphragm components. In both Drosophila garland nephrocytes and mammalian podocytes, the filtration function depends on the correct assembly and plasma membrane localization of the slit diaphragm (*81, 82*). In control nephrocytes, heat fixation followed by immunostaining for slit diaphragm proteins Pyd (ZO-1) and Sns (Nephrin) revealed a characteristic fingerprint-like, homogeneous cortical pattern (Fig. S4B), while medial sections showed predominant localization at the plasma membrane (Fig. S4L). At the ultrastructural level, slit diaphragms appeared as electron-dense structures at the opening of lacunae (Fig. S4C, M).

Syx12L depletion disrupted this organization (Fig. S4D, J, K), resulting in reduced cortical colocalization of Pyd and Sns, decreased slit diaphragm density, and extensive surface regions lacking organized sieving membranes, confirmed by electron microscopy (Fig. S4E). In medial sections (Fig. S4N, T), slit diaphragm proteins were detected deeper within the cell, corresponding to multiple ectopic slit diaphragms observed by electron microscopy (Fig. S4O). Snap29 depletion (Fig. S4F, G, J, K) did not reduce overall cortical slit diaphragm density or component colocalization, but led to the formation of deeper, ectopic slit diaphragms (Fig. S4P, Q, T), likely reflecting aberrant recycling and altered lacunar morphology. These ectopic structures observed both in Syx12L and Snap29 silencing provide a plausible explanation for the ‘clogged’ lacunae and the consequent block in FITC–Avidin uptake (Fig. 2)

In contrast, Ykt6 depletion (Fig. S4H-K) caused a pronounced reduction in cortical slit diaphragm density accompanied with the loss of the ‘fingerprint’ pattern and the marked decrease of the colocalization between Pyd and Sns, indicating defective slit diaphragm assembly and trafficking. Although most slit diaphragm proteins remained associated with the plasma membrane (Fig. S4R, T), electron microscopy revealed only sparse slit diaphragms at the openings of abnormally shallow lacunae (Fig. S4S).

Consistent with these findings, the recycling cargo Cubilin (*83*) showed reduced peripheral localization in all three SNARE knockdowns (Fig. S4U-X). In Syx12L-depleted cells (Fig. S4V), Cubilin accumulated in a pattern similar to the expanded lacunae, whereas in Snap29-depleted cells (Fig. S4W) it retained a lacuna-associated distribution. In contrast, Cubilin predominantly localized to the intracellular region, most probably to endosomes upon Ykt6 depletion (Fig. S4X), consistent with impaired recycling and explaining the reduced endocytic uptake observed under these conditions (Fig. 2).

### Early endosome fusion bypass leads to aberrant endolysosomal ‘Swirl’ formation

To assess later stages of endolysosomal maturation, we analyzed deeper regions of tannic acid–impregnated nephrocytes (Fig. 6A-D) and compared them with Lamp1 immunostaining (Fig. 1 H-K). Control cells (Fig. 6A) contained multiple large endosomes, endolysosomes and smaller, denser lysosomes. Snap29 depletion (Fig. 6C) resulted in numerous small, electron-dense perinuclear lysosomes and cell-wise small endosomes. Ykt6 depletion (Fig. 6D) produced smaller than control but otherwise mature endolysosomes, indicating partial progression of maturation. In Syx12L-silenced cells (Fig. 6B, S5A), small endosomes and lysosomes comparable in size to those observed upon Snap29 depletion were still present, but in addition, strikingly large abnormal structures emerged. These consisted of multiple concentric membrane layers, often enclosing cytosol or smaller vesicles (Fig. 6B) and lipid droplets (Fig. S5A). These most probably correspond to the large Rab7- and Lamp1 positive structures with abnormally ‘thick’ membranes revealed by fluorescent microscopy (Fig. 1C, I). Similar structures were also observed upon silencing of other early endosomal proteins, such as Rab5 (Fig. 6E), Rbsn5 (Fig. 6F, S5B), or Vps8 (Fig. 6G), as well as in Vps8 mutant nephrocytes (Fig. S5 C-E). These structures lacked hallmarks of intraluminal membrane reorganization common for multilamellar late endosomes and instead appeared to arise through vesicle elongation and coiling, based on observation of membrane ends (Fig. S5C’) and enveloped cytoplasm. We therefore refer to them as endolysosomal swirls. Using a Rab2-pHluorin reporter, we found that swirls were Rab2-positive (Fig. 6H, I) but largely lacked the PI3P reporter FYVE-GFP (Fig. S5F) and the clathrin light chain - GFP marker (Fig S5G), confirming that they represent late stages of endolysosomal maturation. Notably, swirl-like structures were absent in Rab2-pHluorin expressing Snap29 (Fig. 6J) or Ykt6-depleted cells (Fig. 6K), suggesting the possibility that their formation may require these SNAREs.

**Figure 6.**
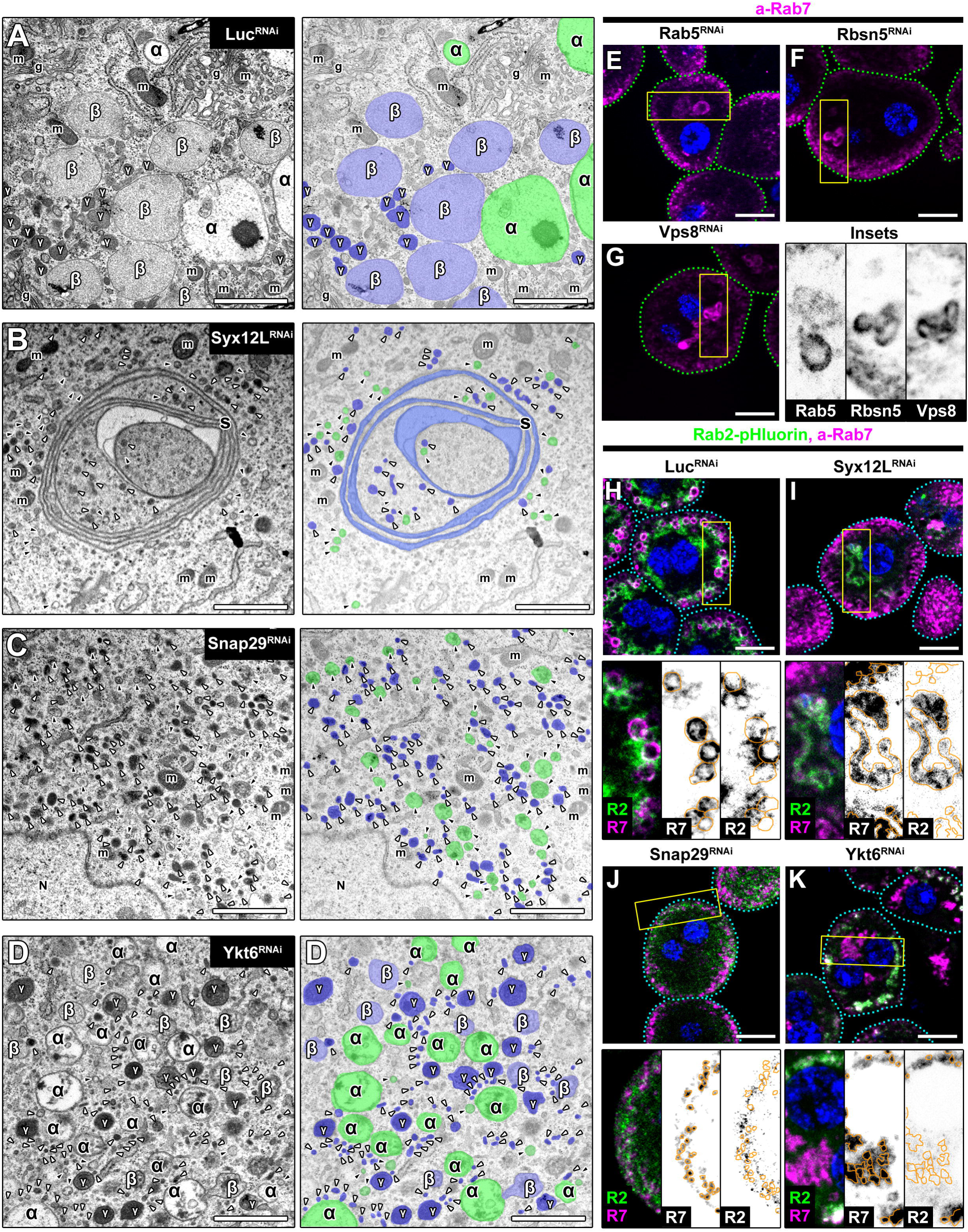
Endosome/lysosome maturation defects and formation of aberrant endo-lysosomal “swirls” upon bypass of early endosomal fusion. A-D) Transmission electron micrographs of deeper regions of nephrocytes. Scale bars: 1 μm. m, mitochondria; g, Golgi apparatus; α, late endosomes (colored green in false-color images); β, γ, lysosomes at different stages of degradation (colored light and dark blue). White arrowheads indicate small lysosome-like structures; black arrowheads indicate small endosomes. Syx12L (B), Snap29 (C), and Ykt6 (D) RNAi cells lack normally sized late endosomes and lysosomes, instead containing predominantly small vesicles, unlike controls (A). Note the presence of a large membrane swirl (S, colored blue in B) in Syx12L RNAi cells. Although Ykt6 RNAi cells contain smaller endolysosomal compartments than controls, they are larger than those in Syx12L or Snap29 RNAi cells. E–G) Inhibition of early endosomal Rab5 (E) or early tethering factors such as Rbsn5 (F) and the miniCORVET subunit Vps8 (G) also induces membrane swirl formation, revealed by anti-Rab7 staining. Blue: nuclei (DAPI). Cell outlines are indicated by dashed cyan lines. Scale bars: 10 μm. H–K) Lysosomes were labeled by expression of Rab2–pHluorin in nephrocytes. All three RNAi conditions (I-K) lack normally sized lysosomes, with Snap29 RNAi cells (J) showing the smallest. The Rab7^+^ membrane swirls observed in Syx12L RNAi cells (I) are Rab2-positive, confirming their lysosomal identity. Blue: nuclei (DAPI). Cell outlines are indicated by dashed cyan lines. Rab7^+^ structures are outlined by continuous orange lines in the black-and-white insets. Scale bars: 10 μm.

Together, these data indicate that bypassing early endosomal fusion can redirect trafficking toward aberrant late endolysosomal structures, whereas loss of Ykt6 favors accumulation of endolysosomes via a distinct pathway.

### Two additional SNARE complexes complement the Syx12L–Snap29–Ykt6 pathway in early endosome maturation

Because partial endosome maturation still occurs in Syx12L- or Ykt6-depleted nephrocytes, we performed epistasis analyses to define the relationships among the three identified SNAREs (Fig. 7). Simultaneous silencing of Syx12L and Ykt6 (Fig. 7D) abolished the formation of Rab7-positive endolysosomal swirls (Fig. 7A, G) and reduced the size of both Rab5- and Rab7-positive endosomes compared with Ykt6 single RNAi (Fig. 7B, H; Fig. S5H, I). This supports the existence of a common Syx12L–Snap29–Ykt6 pathway, while also indicating that Syx12L and Ykt6 can independently promote endosomal maturation toward distinct endpoints. Consistently, combined silencing of Snap29 with either Syx12L (Fig. 7E) or Ykt6 (Fig. 7F) reduced Rab5- and Rab7-positive endosome size relative to the respective single knockdowns (Fig. 7G, H; Fig. S5H, I). Snap29 RNAi also prevented swirl formation in Syx12L-depleted cells (Fig. 7E, G). These results place Snap29 as a central component shared by all early endosomal maturation routes in Drosophila.

**Figure 7.**
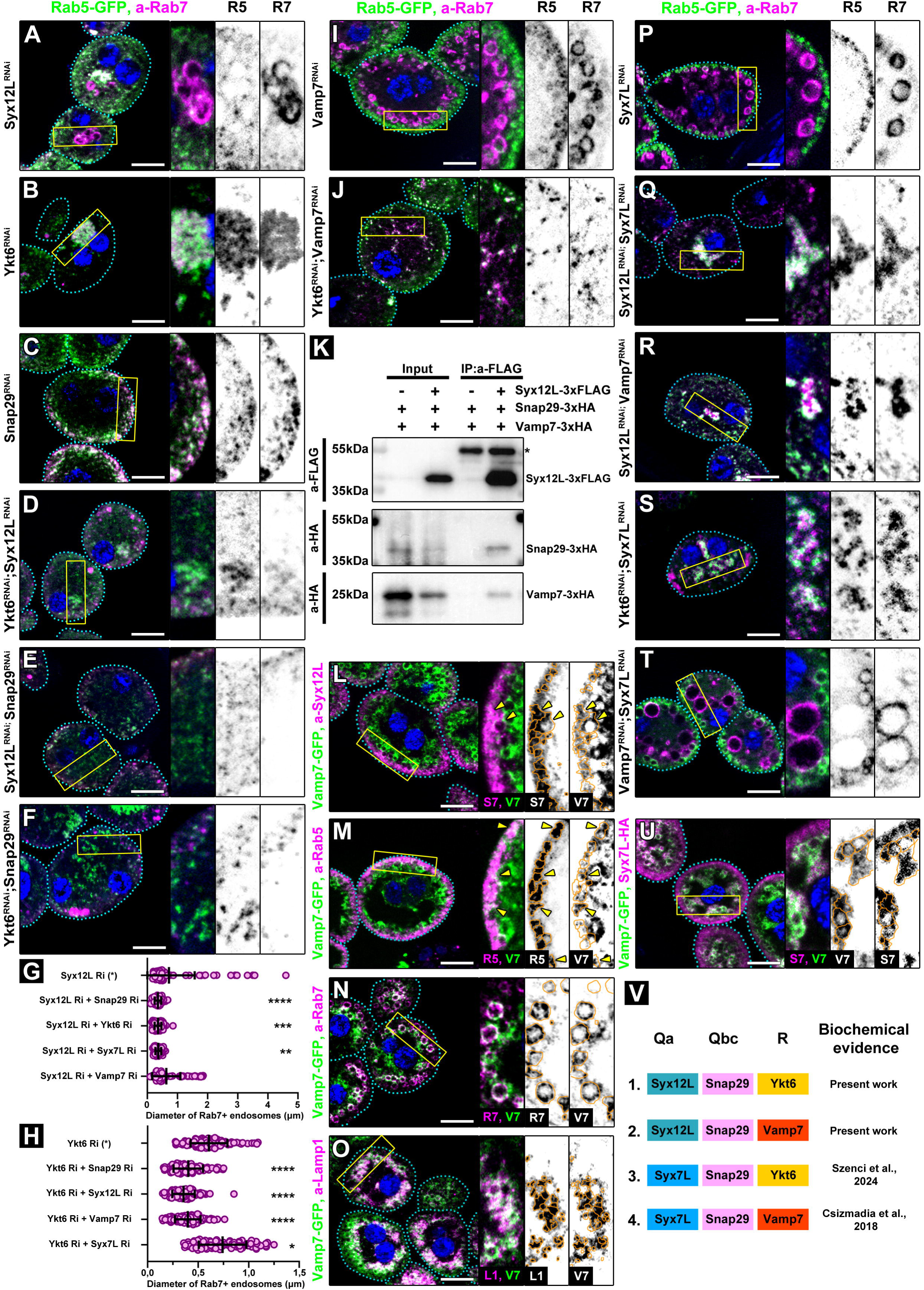
Parallel Syx12L- and Syx7L-based SNARE complexes drive endosome maturation. A–F) Double RNAi of Syx12L with Ykt6 (D) further reduces the size of small endosomes compared with Syx12L (A) or Ykt6 (B) single RNAi, similar to Syx12L (E) or Ykt6 (F) double RNAi in a Snap29 RNAi background (C). Notably, Snap29 and Ykt6 RNAi eliminate the membrane swirls characteristic of Syx12L RNAi cells (A, D, F), indicating that Syx12L-and Ykt6-containing SNARE complexes act in parallel. Blue: nuclei (DAPI). Cell outlines are indicated by dashed cyan lines. Scale bars: 10 μm. G) Quantification of Rab7^+^ endosome size shown in A, D, F, I and P. N = 100 endosomes from 10 cells. Asterisks indicate comparisons to Syx12L single RNAi. **** p < 0.0001, *** p < 0.001, ** p < 0.01. H) Quantification of Rab7^+^ endosome size shown in B, E, F, J and Q. N = 100 endosomes from 10 cells. Asterisks indicate comparisons to Ykt6 single RNAi. **** p < 0.0001, * p < 0.05. I, J) Double RNAi of Ykt6 with Vamp7 results in smaller endosomes compared with either Vamp7 (I) or Ykt6 (B) single RNAi, consistent with parallel routes of action. Blue: nuclei (DAPI). Cell outlines are indicated by dashed cyan lines. Scale bars: 10 μm. K) Co-immunoprecipitation from Drosophila S2R^+^ cells shows that Syx12L–3×FLAG precipitates Snap29–3×HA and Vamp7–3×HA. L–O) Vamp7–GFP predominantly localizes to Rab7^+^ late endosomes (L) and Lamp1^+^ lysosomes (M) in nephrocytes. Only minimal but detectable colocalization is observed with Rab5^+^ early endosomes and with Syx12L (L, M; yellow arrowheads). Blue: nuclei (DAPI). Cell outlines are indicated by dashed cyan lines. Syx12L^+^ (L), Rab5^+^ (M), Rab7^+^ (N) and Lamp1^+^ (O) structures are outlined by continuous orange lines in the black-and-white insets Scale bars: 10 μm. P, Q) Double RNAi of the Qa-SNAREs Syx12L and Syx7L (Q) results in smaller endosomes compared with single RNAi (A and P) and eliminates membrane swirls, consistent with parallel routes of action. Blue: nuclei (DAPI). Cell outlines are indicated by dashed cyan lines. Scale bars: 10 μm. R) Double RNAi of Syx12L with Vamp7 reduces swirl size but does not completely eliminate them, consistent with the action of a parallel Syx7L–Snap29–Ykt6 complex. Blue: nuclei (DAPI). Cell outlines are indicated by dashed cyan lines. Scale bars: 10 μm. S) Double RNAi of Syx7L with Ykt6 results in slightly larger late endosomes compared with Ykt6 single RNAi (B), but smaller compared with Syx7L single RNAi (S), consistent with the presence of a parallel Syx12L–Snap29–Vamp7 complex. Blue: nuclei (DAPI). Cell outlines are indicated by dashed cyan lines. Scale bars: 10 μm. T) Double RNAi of Syx7L with Vamp7 results in larger early endosomes compared with either Vamp7 or Syx7L single RNAi (R, S), consistent with the presence of a parallel Syx12L–Snap29–Ykt6 complex. Blue: nuclei (DAPI). Cell outlines are indicated by dashed cyan lines. Scale bars: 10 μm. U) Vamp7–GFP colocalizes with Syx7L–HA in nephrocytes. Blue: nuclei (DAPI). Cell outlines are indicated by dashed cyan lines. Syx7L-HA^+^ structures are outlined by continuous orange lines in the black-and-white insets Scale bars: 10 μm. V) Schematic illustrating the composition of the four endosomal Syx12L- and Syx7L-based SNARE complexes in flies, based on the current work and previous studies (*14, 56*).

In the absence of Ykt6, Syx12L and Snap29 must therefore act with an alternative R-SNARE. Our in vitro pull-down experiments indicated that Syx12L–Snap29 can associate with Vamp7 (Fig. 3J), which prompted us to test this interaction in vivo. Vamp7 silencing caused enlargement of Rab7-positive endosomes (Fig. 7I, Fig. S5I), consistent with our initial RNAi screen (Fig. 1A; Fig. S2G, I). Importantly, combined Vamp7 and Ykt6 silencing (Fig. 7J) reduced Rab5- and Rab7-positive endosome size compared with Ykt6 RNAi alone (Fig. 7B, H; Fig. S5I). Co-immunoprecipitation from S2 cells confirmed that FLAG–Syx12L forms a complex with both Snap29 and Vamp7 (Fig. 7K). Although Vamp7–GFP localized predominantly to Rab7-positive late endosomes and lysosomal layers, overlapping vesicles with Syx12L and Rab5 were also detected (Fig. 7L–O). Together, these data support a model in which, in the absence of Ykt6, a Syx12L–Snap29–Vamp7 complex can drive early endosomal maturation toward smaller endolysosomes.

Conversely, when Syx12L is absent, Snap29 and Ykt6 promote endosomes to mature into endolysosomal swirls, implying the involvement of an alternative Qa-SNARE. Given the close evolutionary relationship between Syx7L and Syx12L, and the documented ability of Syx7L/STX7 to form complexes with Snap29 and Ykt6 during crinophagy in flies and autophagy in mammalian cells (*45, 56*), we tested Syx7L as a candidate. Syx7L localized mainly to Rab7-positive late endosomes and lysosomal layers (Fig. 4C, D), with minimal but detectable overlap with Rab5-positive early endosomes (Fig. 4B), and its silencing resulted in enlarged late endosomes (Fig. 7P), consistent with our initial screen (Fig. S1F). Notably, simultaneous depletion of Syx12L and Syx7L (Fig. 7Q) abolished swirl formation. This indicates that, in the absence of Syx12L, a Syx7L–Snap29–Ykt6 complex drives the formation of endolysosomal swirls.

Previous work has shown that Syx7L–Snap29–Vamp7 complexes can also form in Drosophila (*14*) We found that Syx7L–HA colocalizes with Vamp7–GFP on endosome/lysosome-like vesicles in nephrocytes (Fig. 7U). To assess their role within the identified pathways, we analyzed Syx12L–Vamp7 and Ykt6–Syx7L double knockdowns. Syx12L–Vamp7 double RNAi (Fig. 7R) reduced swirl size but allowed the formation of larger Rab5- and Rab7-positive vesicles than Syx12L RNAi alone (Fig. 7G, R; Fig. S5H), indicating that the Syx7L–Snap29–Vamp7 complex functions downstream of the Syx7L–Snap29–Ykt6 complex during swirl formation. In contrast, Ykt6–Syx7L double RNAi (Fig. 7S) increased Rab5-positive endosome size compared with Ykt6 single RNAi (Fig. 7B, H; Fig. S5I), suggesting a maturation defect. This implies that the Syx7L–Snap29–Vamp7 complex functions downstream of the Syx12L–Snap29–Vamp7 complex. Consistently, Syx7L–Vamp7 double RNAi cells (Fig. 7T) contained significantly larger early endosomes than either single RNAi condition (Fig. 7B, I; Fig. S5J, K), whereas late endosomes showed a similar trend that did not reach statistical significance. Together, these data indicate that alternative Syx7L/Vamp7-containing complexes primarily act on early endosomes in parallel to the Syx12L–Snap29–Ykt6 pathway.

Collectively, our epistasis, localization, and interaction analyses identify three partially redundant yet mechanistically distinct SNARE-driven pathways operating during early endosome maturation, as well as a late endosomal pathway (Fig. 7V). The Syx12L–Snap29–Ykt6 pathway supports the earliest stages of maturation. In Ykt6-deficient cells, a Syx12L–Snap29–Vamp7 complex mediates limited progression toward small endolysosomes. Conversely, in Syx12L-deficient cells, a Syx7L–Snap29–Ykt6 complex drives maturation toward endolysosomal swirls. All three pathways converge downstream on a Syx7L–Snap29–Vamp7 complex to support further endosomal maturation.

### Distinct Rab5 and Rab7 requirements define the Ykt6- and Syx12L-independent maturation pathways

The existence of Syx12L- and Ykt6-independent pathways prompted us to ask whether these routes differ not only in their SNARE composition but also in their Rab GTPase requirements. Because Rab5 and Rab7 define key stages of endosomal maturation, we analyzed how perturbing each pathway affected their functional involvement.

To assess Rab5 involvement, we first depleted Rab5 and observed that swirl-like structures still formed (Fig. 6E), indicating that their formation is Rab5-independent. We next expressed constitutively active Rab5 (Rab5-CA), which remains GTP-bound and enhances Rab5-dependent fusion events (Fig. 8A–J). In control nephrocytes, Rab5-CA expression (Fig. 8F, U) increased the size of Rab5- and Rab7-positive endosomes, including Rab5/Rab7 double-positive structures. However, in Snap29-depleted cells (Fig. 8G, V), Rab5-CA failed to enlarge early endosomes, indicating that all Rab5-dependent fusion pathways require Snap29. In Syx12L-depleted cells (Fig. 8H, W), Rab5-CA did not increase swirl size, although Rab5-CA remained associated with swirl membranes, further supporting their endolysosomal identity and indicating that Syx12L-independent pathways operate largely independently of Rab5. In contrast, Rab5-CA markedly increased the size of Rab5- and Rab7-positive endosomes in Ykt6-depleted cells (Fig. 8I, X), demonstrating that the Ykt6-independent pathway is Rab5-dependent.

**Figure 8.**
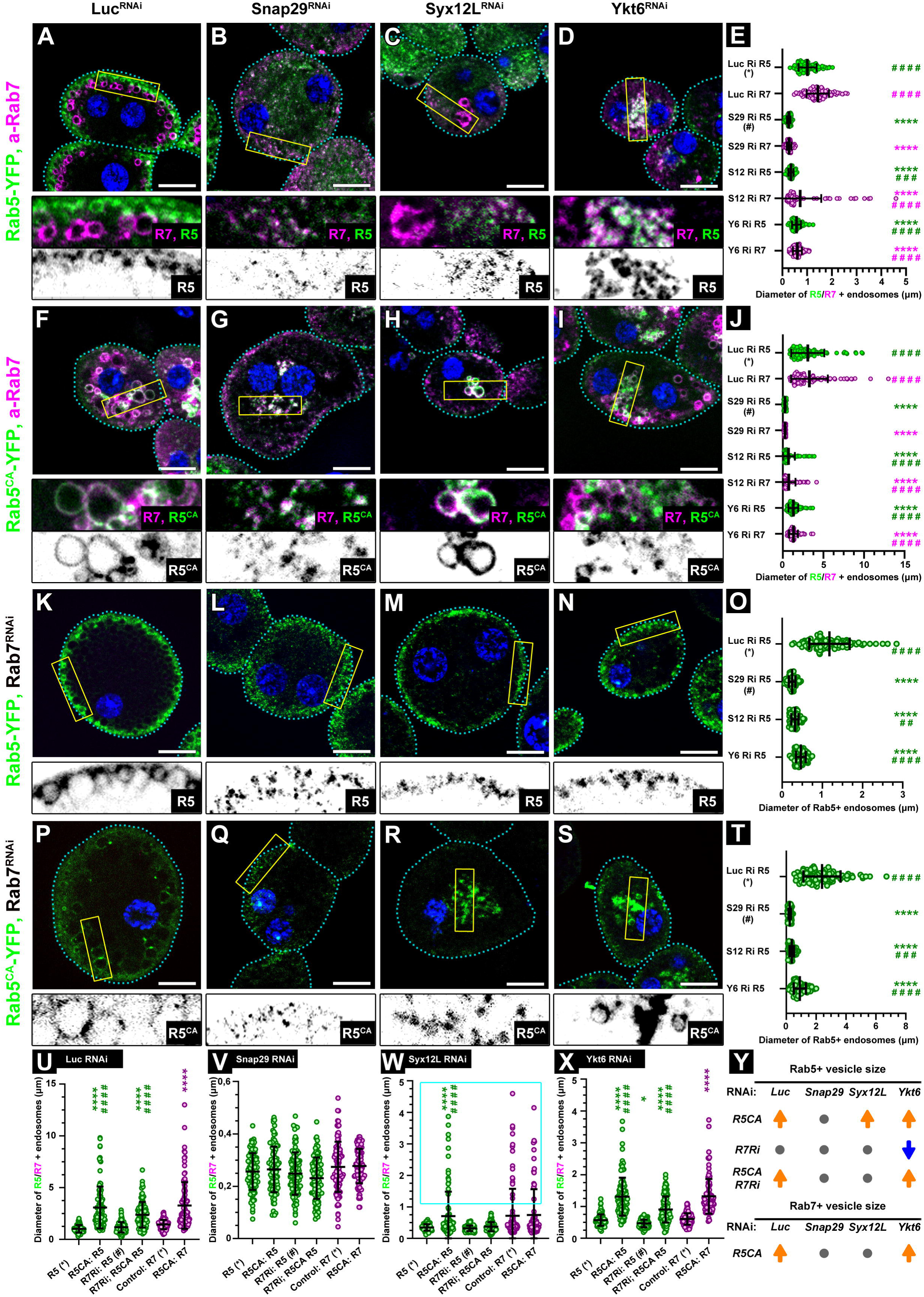
Rab5 activation and Rab7 depletion reveal distinct Rab dependencies of Syx12L-, Snap29-, and Ykt6-dependent pathways. A–D) Rab5-GFP^+^ early endosomes and Rab7^+^ late endosomes are reduced in size in Snap29 RNAi (B), Syx12L RNAi (C), and Ykt6 RNAi (D) nephrocytes compared to controls (A), with Snap29 RNAi cells showing the smallest compartments. Cell outlines are indicated by dashed cyan lines. Blue: nuclei (DAPI). Scale bars: 10 μm. E) Quantification of Rab5^+^ and Rab7^+^ endosome sizes shown in A–D. N = 100 endosomes from 10 cells. Asterisks (*) indicate comparisons to control; hashes (#) indicate comparisons to Snap29 RNAi. ****, #### p < 0.0001. F–I) Expression of constitutively active Rab5 (Rab5-CA-YFP) strongly increases early endosome size in control RNAi cells and moderately in Ykt6 RNAi cells (I), but not in Snap29 RNAi cells (G). Swirl structures observed in Syx12L RNAi cells (H) become Rab5-CA positive, but their size remains unchanged. Cell outlines are indicated by dashed cyan lines. Blue: nuclei (DAPI). Scale bars: 10 μm. These data indicate that some Rab5-dependent routes rely on Vamp7-containing complexes, whereas all Rab5-dependent routes require Snap29. Swirl formation occurs independently of Rab5 activity. J) Quantification of Rab5-CA^+^ and Rab7^+^ endosome sizes shown in F–I. N = 100 endosomes from 10 cells. Asterisks (*) indicate comparisons to control; hashes (#) indicate comparisons to Snap29 RNAi. ****, #### p < 0.0001. K–N) In a Rab7 RNAi background, Rab5-GFP^+^ early endosomes are reduced in size in Snap29 (L), Syx12L (M), and Ykt6 (N) RNAi cells compared to control (K), with Snap29 RNAi showing the smallest endosomes. Cell outlines are indicated by dashed cyan lines. Blue: nuclei (DAPI). Scale bars: 10 μm. O) Quantification of Rab5^+^ endosome size shown in K–N. N = 100 endosomes from 10 cells. Asterisks (*) indicate comparisons to control; hashes (#) indicate comparisons to Snap29 RNAi. ****, #### p < 0.0001; ###, p < 0.001. P–S) In a Rab7 RNAi background, Rab5-CA-YFP increases early endosome size in control RNAi cells and slightly in Ykt6 RNAi cells, but not in Syx12L or Snap29 RNAi cells. Note that Rab7 RNAi eliminates the swirls characteristic of Syx12L RNAi. Cell outlines are indicated by dashed cyan lines. Blue: nuclei (DAPI). Scale bars: 10 μm. These data indicate that Rab7-dependent routes also rely on Vamp7-containing complexes and that swirl formation is Rab7 dependent. T) Quantification of Rab5-CA^+^ endosome size shown in P–S. N = 100 endosomes from 10 cells. Asterisks (*) indicate comparisons to control; hashes (#) indicate comparisons to Snap29 RNAi. ****, #### p < 0.0001; ##, p < 0.01. U–X) Summary quantifications of Rab5^+^, Rab5-CA^+^, and Rab7^+^ endosome sizes in different RNAi backgrounds. U) Control RNAi (A,F,K,P), V) Snap29 RNAi (B,G,L,Q), W) Syx12L RNAi (C,H,M,R), X) Ykt6 RNAi (D,I,N,S). N = 100 endosomes from 10 cells. Asterisks (*) indicate comparisons to Rab5-GFP expressing single RNAi; hashes (#) indicate comparisons to Rab5-CA in Rab7 plus gene-of-interest double RNAi cells. ****, #### p < 0.0001; * p < 0.05. Y) Summary diagram of relative endosome sizes across conditions. Grey dot: no change; orange and blue arrows: size increase and decrease, respectively.

We next assessed Rab7 involvement (Fig. 8K–O). In control cells, Rab7 depletion did not change early endosome size, as expected (Fig. 8K). However, in all three SNARE knockdowns in a Rab7 RNAi background (Fig. 8L–O), early endosomes were smaller than in Rab7 RNAi alone, indicating that both Syx12L- and Ykt6-dependent pathways can engage Rab7. Notably, early endosomes were further reduced in size in Ykt6-depleted cells when Rab7 was also silenced (Fig. 8D, N, X). Similarly, when constitutively active Rab5 (Rab5-CA) was expressed in combination with Rab7 silencing (Fig. 8P–T), enlarged Rab5-positive endosomes formed only in Ykt6-depleted cells (Fig. 8S), and swirls were not observed even when Syx12L was silenced (Fig. 8R). These results indicate that the Syx12L-independent pathway and swirl formation require Rab7, whereas the Ykt6-independent pathway can proceed, at least partially, without Rab7.

Together, these experiments demonstrate that the three early endosomal maturation pathways differ not only in their SNARE composition but also in their Rab GTPase requirements. This strongly argues that these pathways are not simple compensatory mechanisms but represent distinct, regulated routes of early endosome maturation, likely engaging different tethering factors and regulatory inputs.

## DISCUSSION

Endosomal maturation has traditionally been viewed as a largely linear process, driven by sequential Rab GTPase conversion and the ordered deployment of tethering factors and SNARE complexes. While this framework has been invaluable, accumulating evidence - particularly from mammalian systems - has suggested that multiple fusion machineries may operate at the same trafficking step. However, extensive redundancy in these systems has made it difficult to disentangle core mechanisms from compensatory effects. By exploiting the reduced SNARE repertoire of Drosophila nephrocytes (*49–51*), our study reveals that early endosome maturation is executed by multiple, parallel SNARE-dependent pathways, each with distinct molecular requirements, functional consequences, and morphological outputs. Our data establish that early endosomal fusion in Drosophila does not rely on a single indispensable SNARE complex. Instead, at least three partially redundant SNARE assemblies promote maturation beyond the early endosomal stage (Fig. 9). These pathways converge downstream but differ markedly in their reliance on specific SNAREs, Rab GTPases, and recycling dynamics. Importantly, these pathways are not interchangeable backups; rather, they are all required for shaping endosomal architecture, recycling flux, and cargo handling.

**Figure 9.**
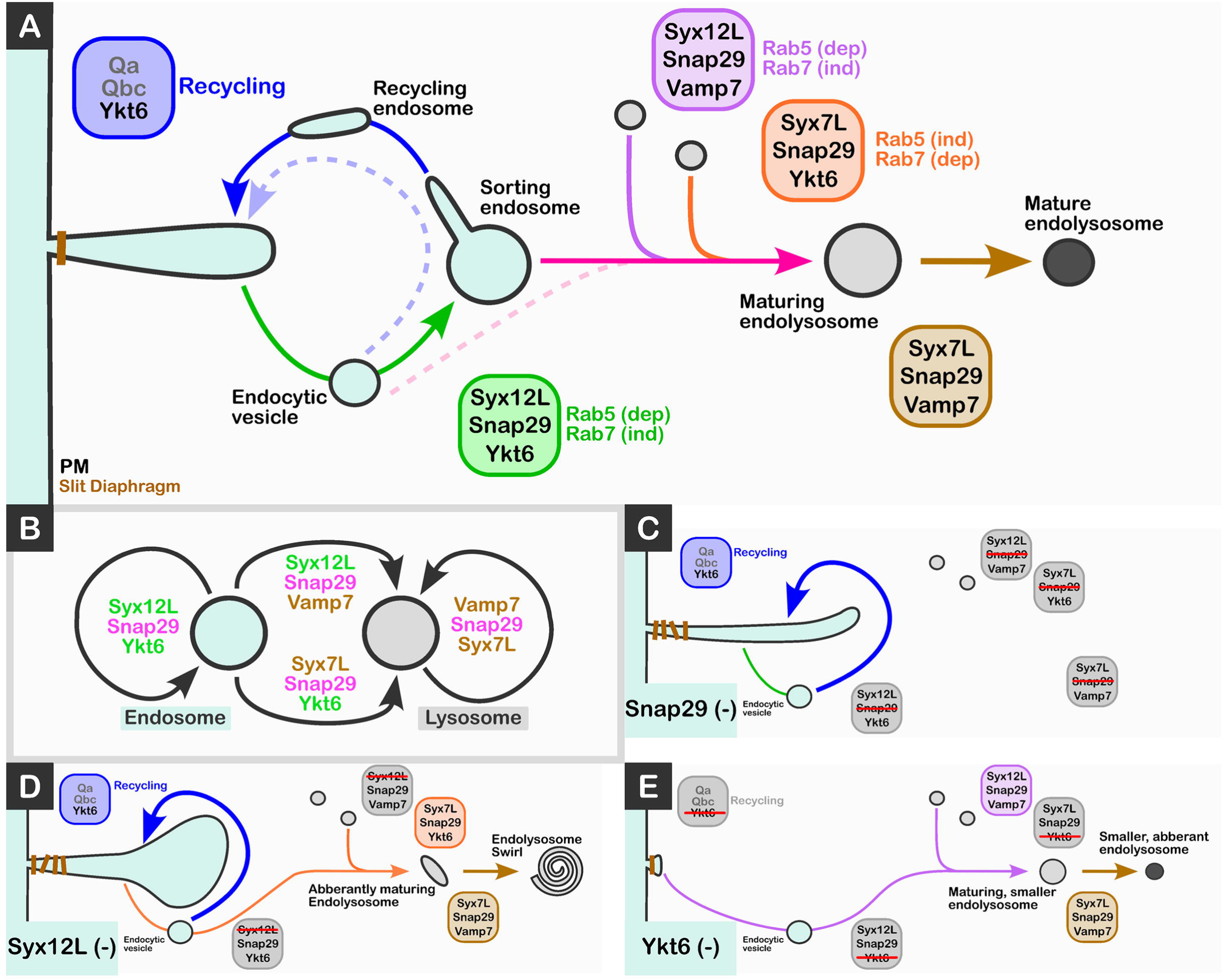
Proposed model of endosome maturation driven by Syx12L- and Syx7L-containing SNARE complexes in Drosophila nephrocytes. A) In control cells, the Syx12L–SNAP29–Ykt6 complex mediates the earliest Rab5-dependent fusions between early endocytic vesicles and early endosomes. Subsequent maturation involves both Rab5-dependent and -independent, and Rab7-independent and - dependent fusion events driven by Syx12L–SNAP29–Vamp7 and Syx7L–SNAP29–Ykt6 complexes, respectively. Fusions of late endosomes or lysosomes are executed primarily by the Syx7L–SNAP29–Vamp7 complex. Recycling endosomes are fused with the plasma membrane (PM) by a yet unidentified Ykt6-containing complex. Balanced recycling versus degradation maintains slit diaphragm (SD) integrity and lacunar depth. B) Simplified schematic of the SNARE network highlighting SNAP29 as a central shared component. C) In SNAP29 knockdown cells, none of the endosomal/lysosomal SNARE complexes are functional. As a result, endosomes and lysosomes remain small and unfused. A large fraction of endosomes is redirected into the recycling pathway, where Ykt6 mediates their fusion with the PM, leading to lacunar deepening and clogging of SDs. D) In Syx12L knockdown cells, only Syx7L-containing complexes remain functional. Early endosomes fail to undergo homotypic fusion and their fusion with more mature compartments is repressed. Maturation is not completely blocked: Syx7L–SNAP29–Ykt6 drives their conversion into small endolysosome-like structures, and Syx7L–SNAP29–Vamp7 mediates their homotypic fusion into endolysosomal “swirls.” In parallel, many unfused early endosomes are redirected into the recycling pathway, where Ykt6-dependent fusion with the PM deepens lacunae and disrupts SD architecture. E) In Ykt6 knockdown cells, homotypic early endosome fusion is lost due to inactivation of the Syx12L–SNAP29–Ykt6 complex, and fusion of early endosomes with more mature compartments is also impaired. Maturation is partially maintained through Vamp7-containing complexes, which generate smaller endolysosomes. Because Ykt6 is also required for recycling, its absence diverts most endosomal cargo toward degradation, leading to progressive loss of lacunae due to continuous uptake of their components.

The existence of parallel fusion routes explains why early endosomal maturation can proceed, albeit aberrantly, upon loss of individual fusion components, while simultaneously producing strikingly distinct phenotypes depending on which early-acting SNARE is disrupted. These findings argue against a strictly hierarchical SNARE cascade and instead support a model in which SNARE composition itself is a determinant of endosomal fate.

Among the SNAREs analyzed, Snap29 occupies a unique position. Genetic, morphological, and functional analyses demonstrate that all identified early endosomal fusion routes depend on Snap29. Loss of Snap29 abolishes compensatory fusion events, disrupts maturation at multiple stages, and produces phenotypes that cannot be bypassed through activation of alternative pathways. These findings place Snap29 at the core of early endosome maturation and its progression toward the late endosome–lysosome system in Drosophila.

This central role is consistent with extensive prior evidence implicating Snap29 across lysosome-related pathways. Snap29 is required for autophagosome–lysosome fusion through interactions with Syx17 and Vamp7, and for secretory granule maturation and crinophagy via complexes containing Syx7L (previously termed Syx13) together with Vamp7 or Ykt6 (*14, 16, 45, 54, 56, 59, 60, 62, 75, 84*). In addition, Snap29 mutations cause defects in eye pigment granule formation, a lysosome-related organelle in flies (*54, 85*). Consistent with this view, Snap29-deficient cells contain lysosomes that remain small and fail to undergo homotypic or heterotypic fusion.

This system-wide role is consistent with Snap29’s identity as a TM-domain–lacking Qbc-SNARE, which enables its availability to participate in multiple complexes because it is soluble and not membrane-bound. Beyond its biochemical versatility, our data indicates that Snap29 acts as a central integrator of endolysosomal fusion, coordinating parallel pathways rather than serving a single step. This framework provides a mechanistic explanation for the severe endolysosomal defects observed upon SNAP29 loss in humans, such as in CEDNIK syndrome, where Snap29 dysfunction cannot be compensated by alternative SNARE assemblies (*86, 87*).

A notable outcome of our analysis is the clarification of the roles of Syntaxin7 family members in Drosophila. In mammalian cells, STX7 is generally regarded as a late endosomal/lysosomal Qa-SNARE (*35–38, 45*), whereas STX12 has been described to function at early endosomes (*34*). In contrast, in Drosophila, Syx12L (previously termed Syx7) was originally assigned an early endosomal role (*52, 53*), while Syx7L (previously termed Syx13) has been implicated in a specialized late endolysosomal fusion pathway - secretory granule–lysosome fusion (crinophagy) - suggesting a primary function at later stages of the pathway (*14, 56*).

Our functional, genetic, and localization data demonstrate that Drosophila Syx12L (formerly Syx7) operates predominantly at early endosomes, where it mediates fusion events preceding late endosomal maturation. In contrast, late endosomal fusion is mediated by Syx7L-containing complexes (formerly Syx13), supporting a reassignment of functional orthology between Drosophila and mammalian Syntaxin family members. Although the apparent mismatch between mammalian and Drosophila Syntaxin7 family assignments has persisted in the literature (*57*), it has not been explicitly addressed. Based on our findings and previous evolutionary analyses (*49, 50*), we propose that Syntaxin7 family proteins in vertebrates underwent paralogous compensation following gene duplication, likely associated with functional inactivation of the ancestral Syx7′b lineage due to insertion of a Harbinger transposon (*78*). In contrast, Drosophila Syx12L and Syx7L retained their ancestral functions, corresponding functionally to human STX12 and STX7, respectively.

One of the most striking insights from this study is the tight coupling between early endosomal fusion and endosomal recycling. Disruption of early endosome fusion—either by SNARE or tethering factor depletion—does not simply stall maturation but instead reroutes membrane traffic toward recycling pathways. This is manifested in nephrocytes by elongated lacunar channels.

These observations indicate that early endosome fusion acts as a rate-limiting checkpoint that determines whether internalized membrane proceeds toward degradation or is rapidly returned to the plasma membrane. When fusion is impaired, endocytic vesicles are preferentially shunted into recycling routes, preserving membrane flux but altering compartment morphology and cargo distribution. Based on our nephrocyte data and independent studies in wing cells (*65*), Ykt6 is required for endosomal recycling. These findings suggest that Ykt6 plays a particularly prominent role in endocytic processes, acting not only in endosomal fusion but also in restraining excessive recycling, thereby maintaining balance between degradative and recycling pathways.

Our epistasis and Rab5 activation experiments further reveal that the identified SNARE pathways differ fundamentally in their Rab requirements. While the Ykt6-independent pathway (Syx12L–Snap29–Vamp7 complex) relies on Rab5 activity, the Syx12L-independent route (Ykt6–Snap29–Syx7L) can bypass Rab5 and proceed in its absence. Rab7 dependence is likewise pathway-specific. Certain maturation routes retain fusion competence even when Rab7 function is compromised. These findings challenge simplified models of Rab5-to-Rab7 conversion as an obligatory step in endosome maturation. Instead, they support a model in which SNARE identity can partially uncouple fusion from canonical Rab logic, allowing maturation to proceed through alternative routes (Fig. 9).

The formation of multilayered endolysosomal swirls represents a distinctive morphological outcome of impaired early endosome fusion. Our ultrastructural, genetic, and marker analyses indicate that these structures arise not from intraluminal vesicle accumulation but from elongation and coiling of endolysosomal membranes, likely driven by unbalanced fusion events and membrane remodeling. Their dependence on specific SNARE combinations, and their absence when both Syx12L and Syx7L are removed, supports the idea that they represent a defined alternative maturation route rather than a nonspecific pathology. The appearance of similar structures upon perturbation of early endosomal tethers (miniCORVET or Rbsn5) or Rab5 further reinforces the conclusion that swirls reflect a general consequence of bypassing canonical early fusion steps. These structures thus provide a morphological readout of altered SNARE pathway engagement.

Together, our findings support a revised model of early endosome maturation in which multiple SNARE-dependent pathways operate in parallel, non-interchangeable ways, each capable of driving fusion to a limited extent and each associated with distinct Rab usage, recycling behavior, and morphological outcomes. Rather than serving as simple backups, these pathways act together, providing the endosomal system with flexibility, robustness, and regulatory capacity. Such an organization allows cells to adapt endosomal trafficking to physiological demands, maintain membrane homeostasis, and buffer against perturbations.

A longstanding question in membrane trafficking is whether SNAREs themselves act as targeting signals that confer vesicle identity (*76, 88–90*). Recent biochemical and structural studies have challenged this view, suggesting that beyond the QabcR rule of complex formation (*90*), SNARE pairing is often permissive, and that the surrounding molecular context plays a dominant role in specifying productive fusion events (*76, 89, 91–93*). Given that Qa, Qb, Qc, and R SNARE families have undergone hundreds of millions of years of independent evolution (*49, 50, 94*), the conservation of such interaction promiscuity argues for its functional importance. Our study provides a clear in vivo example of this principle: early endosome maturation in Drosophila relies on partially overlapping SNARE complexes that differ by only a single Qa or R component, yet these drive distinct fusion outcomes within a robust network. Rather than acting as strict determinants of vesicle identity, SNAREs therefore appear to function as permissive fusion modules that expand the range of possible trafficking routes. In this view, vesicle identity is not encoded by a single SNARE, but emerges from the combinatorial assembly of multiple factors, allowing trafficking pathways to remain flexible while preserving overall system robustness.

In conclusion, our study demonstrates that early endosome maturation is not governed by a single linear SNARE cascade but by a network of partially overlapping fusion machineries (Fig. 9). By defining the composition, regulation, and consequences of these pathways, we provide a framework for understanding how endosomal fate decisions are encoded at the molecular level.

## MATERIALS AND METHODS

### Molecular cloning

For pull-down experiments, DNA constructs encoding Drosophila SNAREs were generated by PCR using primers listed in Table S1. PCR products were cloned into the BamHI–XhoI sites of the pETMBP vector, except for the Syx7 SNARE domain, which was cloned into the BamHI–NotI sites of pETARA (*95*). Inserts were assembled using the NEBuilder HiFi DNA Assembly Cloning Kit (New England Biolabs, Massachusetts, USA). DH5α competent *E. coli* cells (New England Biolabs, Massachusetts, USA) were transformed and plated on LB–ampicillin agar plates. After overnight growth at 37 °C, single colonies were used to inoculate 4 ml LB cultures, and plasmids were purified using the GeneJet Plasmid Miniprep Kit (Thermo Fisher Scientific, Massachusetts, USA).

For immunoprecipitation experiments, the coding region of Syx7 was amplified from Drosophila cDNA (primers in Table S1). To generate N-terminally 3×FLAG-tagged Syntaxin7, the insert was cloned into the NotI–Acc65I sites of pUAST-attB-Flag-dTSC2 WT (Addgene #111807). For N-terminally 3×HA-tagged Syntaxin7, the insert was cloned into the Acc65I–XbaI sites of pUAST-3xHA-V5-RpL10Ab (Addgene #125223). All constructs were verified by Sanger sequencing (Microsynth AG, Switzerland). UAS-VAMP7-3xHA, UAS-SNAP29-3xHA, pMT-GAL4, and UAS-Ykt6-3xHA constructs were described previously (*59, 62*).

### Protein production and purification

BL21(DE3) *E. coli* cells were transformed with pulldown constructs and grown at 37 °C to OD600 = 0.4. Protein expression was induced with 500 µM IPTG and cultures were incubated overnight at 18 °C. Cells were harvested by centrifugation at 3315 g for 5 min (Beckman JLA-9.1000 rotor, Beckman Coulter Life Sciences, Indianapolis, IN, USA), resuspended in lysis buffer (50 mM Na2HPO4 pH 8.0, 300 mM NaCl, 20 mM imidazole, 0.1% Triton X-100, 2 mM β-mercaptoethanol, 10 mg/ml lysozyme), and lysed by freeze–thaw cycles followed by sonication. Lysates were clarified at 48 400 g for 30 min at 4 °C. Supernatants were incubated with Ni Sepharose Excel resin (Cytiva, Chicago, IL, USA) for 1 h at 4 °C with rotation. Bound proteins were washed sequentially and eluted in elution buffer (20 mM Tris-HCl pH 8.0, 200 mM NaCl, 400 mM imidazole, 10% glycerol, 2 mM β-mercaptoethanol). Eluates were concentrated if needed (Amicon Ultra-15, 10 kDa MWCO, Millipore, Massachusetts, USA), supplemented with 2 mM TCEP, aliquoted, and stored at −80 °C.

### GST pull-down

GST pull-down assays were performed using Glutathione Sepharose High-Performance resin (Cytiva, Chicago, IL, USA). GST-Syx7-SNARE-6×His was bound to resin for 1 h at 4 °C. After washing, prey proteins were added and incubated for 30 min at 4 °C. Resin was washed three times and bound proteins were eluted in Laemmli buffer, boiled, and analyzed by SDS-PAGE followed by Coomassie Brilliant Blue staining.

### S2R^+^ cell culture, transfection and immunoprecipitation

S2R^+^ cells (DGRC #150, RRID:CVCL Z831) were maintained at 26 °C in Insect XPress medium (Lonza) with 10% FBS and 1% Penicillin–Streptomycin. For IP experiments, 7.5 × 10^6^ cells were transfected using the calcium phosphate method. Cells were co-transfected with pMT-GAL4 and protein expression was induced with 500 µM CuSO_4_ 24 h post-transfection. Cells were lysed in Triton X-100 buffer and cleared at 20 000 g. Inputs were reserved, and lysates were incubated with anti-HA agarose (MBL-561-8) or anti-FLAG M2 agarose (Millipore A2220) for 3 h at 4 °C. Beads were washed and eluted in Laemmli buffer. Proteins were separated by SDS-PAGE and transferred to PVDF membranes using the Bio-Rad Trans-Blot Turbo system.

### Antibodies

Details of antibodies are listed in Table S1. Anti-Cubilin, anti-Lamp1, anti-Sns and anti-Syx12L (previously Avl/Syx7) were kindly provided by Joaquim Culi (*96*), Andreas Jenny (*97*), Michael Krahn (*98*) and David Bilder (*52*), respectively. Anti-Snap29 and anti-GFP were already in our hands (*62*) ; all other antibodies were obtained from commercial suppliers.

### Fly work and treatments

Flies were maintained at 25[°C on standard minimal food. Wandering third-instar (L3) larvae of both sexes were used for all experiments, except for those involving the X-linked Ykt6-HA transgene, for which only female progeny were analyzed to ensure consistent expression levels. Sources and details of fly stocks are listed in Table S1. Sources and details of fly stocks are listed in Table S1. UAS-GFP-VAMP7, UAS-pHluorinSE-Rab2, and UAS-Syx13-HA were kindly provided by Amy Kiger, Ole Kjaerulff, and Fen-Biao Gao, respectively (*57, 70, 99*). UAS-Ykt6.HA and vps8[1] were already in our hands (*30, 62*), and all other lines were obtained from stock centers. Detailed genotypes of experimental animals are listed in Table S2. FlyBase was used for gene and stock annotation (*100*).

Silver nitrate uptake was performed as described previously (*30*). Briefly, adult flies were allowed to lay eggs on food containing 0.005% AgNO_3_ (20 g yeast in 35 ml of 1.5% agar). Garland nephrocytes were dissected in PBS and fixed in 4% formaldehyde in PBS for 30 min at room temperature (RT). Images were acquired using a Plan-Neofluar 40×/0.75 NA objective on an AxioImager Z1 microscope with an AxioCam ICc camera (Zeiss). Before imaging, samples were exposed to UV light for 10–15 s using an HBO 100 lamp and a DAPI filter set to enhance contrast of intracellular silver inclusions (*30*).

### Immunohistochemistry

Nephrocytes from wandering L3 larvae were dissected in PBS and fixed in 4% formaldehyde in PBS for 45 min at RT. Samples were washed three times for 10 min and permeabilized in PBS containing 0.1% Triton X-100 (PBTX) for 15 min (*30*).

For slit diaphragm visualization, samples were heat-fixed (*98*). Nephrocytes were dissected in ice-cold M3 medium, transferred to boiling 0.03% Triton X-100 in PBS, and incubated for 22 s in boiling water. Samples were then processed in 0.03% PBTX.

Fixed samples were blocked in 5% fetal bovine serum (FBS) in PBTX for 30 min at RT and incubated with primary antibodies overnight at 4 °C. After washing, samples were incubated in PBTX containing 4% NaCl for 15 min to reduce nonspecific binding, followed by secondary antibody incubation for 3 h at RT. Samples were mounted in DAPI-containing VECTASHIELD.

### Channel diffusion assay

Nephrocytes were fixed in ice-cold 4% formaldehyde in PBS for 5 min to stop endocytosis. Samples were incubated with Alexa Fluor 647–BSA in M3 medium for 15 min at RT, followed by a second fixation for 45 min. Samples were washed and mounted in DAPI-containing VECTASHIELD (Vector) (*101*).

### Uptake assay

Nephrocytes were dissected in ice-cold M3 medium and incubated with FITC-avidin (1:100) and, where indicated, fluorescent dextran (10 kDa; 1:20) for 5 min (early endosomes) or 30 min (late endosomes/endolysosomes) at RT. For tracer clearance assays, samples were chased in tracer-free medium for 10 min before fixation after the 30 min pulse. Samples were fixed in 4% formaldehyde for 45 min, washed, and mounted.

### Fluorescence imaging

Images were acquired at RT using an AxioImager.M2 microscope with an ApoTome2 unit, a Plan-Apochromat 63×/1.40 oil objective, and an Orca Flash 4.0 LT sCMOS camera. Imaging settings were identical for all comparable experiments. Images were processed using Zeiss Efficient Navigation 2 and assembled in Adobe Photoshop CS4.

### Electron microscopy

Nephrocytes were fixed overnight at 4 °C in 3.2% paraformaldehyde, 1% glutaraldehyde, 1% sucrose, and 0.028% CaCl_2_ in 0.1 N sodium cacodylate buffer (pH 7.4). Samples were post-fixed in 0.5% osmium tetroxide for 1 h and half-saturated uranyl acetate for 30 min, dehydrated in graded ethanol, and embedded in Durcupan. Ultrathin sections were stained with Reynolds’ lead citrate and examined on a JEM-1011 transmission electron microscope at 80 kV.

For tannic acid impregnation (*102*), samples were treated after osmification with 1% tannic acid in 0.1 M cacodylate buffer (pH 7.4) for 30 min at RT, and uranyl acetate staining was omitted.

### Quantification and statistical analysis

Fluorescent structures were quantified from original, unmodified single optical sections using ImageJ/Fiji. Signal thresholds for each channel were set consistently by the same person across all samples. Cells were randomly selected for analysis, and only cells with nuclei present in the focal plane were included to ensure sampling of both perinuclear and peripheral regions, except for measurements of slit diaphragm surface density and colocalization. Each experiment was performed at least twice on different days.

Endosome size measurements were obtained for Rab5-GFP/YFP, Rab5-CA-YFP, anti-Rab7, and anti-Lamp1–positive compartments by measuring vesicle diameters using the Line tool in ImageJ. For each genotype, 3–6 animals were analyzed, and 10 endosomes were measured per cell from 10 cells, yielding n = 100 endosomes per condition.

Pairwise colocalization between reporter proteins and antibody stainings was quantified using the Threshold Overlap Method. Both channels were thresholded identically, and the area fraction of each cell covered by the fluorescent signal was measured. Overlapping signal was determined using the Image Calculator (AND function). Threshold overlap scores (TOS) were calculated according to Sheng et al. (*73*) from 12 cells per genotype and condition (n = 12), using samples from 4–7 animals.

Fluorescent tracer uptake (FITC–avidin and dextran) was quantified by measuring the area fraction of each cell covered by tracer signal using uniform threshold settings. Cell outlines were determined by enhancing autofluorescence. For each genotype and condition, 4–7 animals were analyzed and fluorescence was quantified in 13-15 cells (n = 13-15).

Lacunar depth (BSA–Alexa Fluor 647) was estimated by outlining the total cell area and the area defined by the lacuna endpoints. The difference in radii of the corresponding calculated circles from the area was used as an approximation of lacunar depth. Measurements were obtained from 15 cells per genotype from 4–6 animals (n = 15).

Surface density of slit diaphragm components (Pyd and Sns) was quantified from cortical sections of heat-fixed, antibody-stained nephrocytes. Using the Line tool in ImageJ, a line (6–15 μm, depending on cell size) was drawn perpendicular to slit diaphragm stripes. Peak numbers were extracted from Plot Profile outputs and normalized to a 10 μm line length. Measurements were obtained from 12 cells per genotype using samples from 4–7 animals (n = 12).

Maximum signal depth of slit diaphragm components (Pyd and Sns) was measured in medial optical sections by drawing a line perpendicular to the cell surface and recording the depth of detectable signal. For each genotype, 12 cells from 4–7 animals were analyzed (n = 12).

Statistical analyses were performed using GraphPad Prism. Normality was assessed using the D’Agostino–Pearson test. For non-normally distributed datasets, Mann-Whitney test (if two datasets were compared) or Kruskal–Wallis tests with Dunn’s multiple comparisons were used. For normally distributed datasets, Student’s T-test (if two datasets were compared) or one-way ANOVA with appropriate post hoc tests was applied. P value thresholds were defined as * or #, p < 0.05; ** or ##, p < 0.01; *** or ###, p < 0.001; **** or ####, p < 0.0001. Figure legends specify reference groups for all comparisons.

Dot plots display individual data points with mean ± standard deviation. Full statistical details are provided in Table S3.

## ACKNOWLEDGEMNETS

We thank S. Pálfia, M. Truszka, and I. Répássy for technical assistance and colleagues listed in Materials and Methods for providing reagents.

## Data and materials availability

All data needed to evaluate the conclusions in the paper are present in the paper and/or the Supplementary Materials.

## Author contributions

**D.H.:** Conceptualization, Formal analysis, Funding acquisition, Investigation, Methodology, Validation, Visualization and Writing – original draft, Writing – review & editing. **M.M.:** Conceptualization, Investigation, Methodology, Validation, Visualization and Writing – original draft, Writing – review & editing. **A.R.:** Investigation, Methodology and Writing – original draft. **I**.B.: Investigation. **D.B.:** Investigation. **A.N.:** Investigation, Writing – review & editing. **V.B.:** Formal analysis and Investigation. **Zs. S.-V.:** Investigation and Methodology. **G.J.:** Funding acquisition, Methodology and Resources, Writing – review & editing. **P.L.:** Conceptualization, Funding acquisition, Investigation, Methodology, Project administration, Resources, Supervision, Validation, Visualization and Writing – original draft, Writing – review & editing.

## SUPPLEMENTAL INFORMATION

**Figure S1.** Representative micrographs of the SNARE RNAi screen: Qa, Qb, and Qc SNAREs

**Figure S2.** Representative micrographs and quantification of the SNARE RNAi screen: Qbc and R SNAREs

**Figure S3:** Additional nephrocyte data (uptake and TEM)

**Figure S4:** The filtration system is defective in Syx12L, Snap29 and Ykt6 depleted nephrocytes, and additional lacunar depth data

**Figure S5:** Additional characterization of endolysosomal “swirl” structures and additional quantification data to Figure 7.

**Table S1.** Fly stocks, antibodies, and reagents used in this study.

**Table S2.** Genotypes of animals used in this study.

**Table S3.** Statistical details.

## REFERENCES

1. T. C. Sudhof, J. E. Rothman, Membrane fusion: grappling with SNARE and SM proteins. Science (New York, N.Y.) 323, 474–477 (2009).

2. D. Kummel, C. Ungermann, Principles of membrane tethering and fusion in endosome and lysosome biogenesis. Current opinion in cell biology 29, 61–66 (2014).

3. W. Hong, S. Lev, Tethering the assembly of SNARE complexes. Trends in cell biology 24, 35–43 (2014).

4. W. Wickner, R. Schekman, Membrane fusion. Nature structural & molecular biology 15, 658–664 (2008).

5. M. Zick, W. T. Wickner, A distinct tethering step is vital for vacuole membrane fusion. eLife 3, e03251 (2014).

6. D. G. McEwan, D. Popovic, A. Gubas, S. Terawaki, H. Suzuki, D. Stadel, F. P. Coxon, D. Miranda de Stegmann, S. Bhogaraju, K. Maddi, A. Kirchof, E. Gatti, M. H. Helfrich, S. Wakatsuki, C. Behrends, P. Pierre, I. Dikic, PLEKHM1 regulates autophagosome-lysosome fusion through HOPS complex and LC3/GABARAP proteins. Molecular cell 57, 39–54 (2015).

7. S. Pankiv, E. A. Alemu, A. Brech, J. A. Bruun, T. Lamark, A. Overvatn, G. Bjorkoy, T. Johansen, FYCO1 is a Rab7 effector that binds to LC3 and PI3P to mediate microtubule plus end-directed vesicle transport. The Journal of cell biology 188, 253–269 (2010).

8. R. van der Kant, A. Fish, L. Janssen, H. Janssen, S. Krom, N. Ho, T. Brummelkamp, J. Carette, N. Rocha, J. Neefjes, Late endosomal transport and tethering are coupled processes controlled by RILP and the cholesterol sensor ORP1L. Journal of cell science 126, 3462–3474 (2013).

9. H. J. Balderhaar, C. Ungermann, CORVET and HOPS tethering complexes–coordinators of endosome and lysosome fusion. Journal of cell science 126, 1307–1316 (2013).

10. J. A. Solinger, A. Spang, Tethering complexes in the endocytic pathway: CORVET and HOPS. The FEBS journal 280, 2743–2757 (2013).

11. P. Lorincz, S. Toth, P. Benko, Z. Lakatos, A. Boda, G. Glatz, M. Zobel, S. Bisi, K. Hegedus, S. Takats, G. Scita, G. Juhasz, Rab2 promotes autophagic and endocytic lysosomal degradation. The Journal of cell biology 216, 1937–1947 (2017).

12. N. Fujita, W. Huang, T. H. Lin, J. F. Groulx, S. Jean, J. Nguyen, Y. Kuchitsu, I. Koyama-Honda, N. Mizushima, M. Fukuda, A. A. Kiger, Genetic screen in Drosophila muscle identifies autophagy-mediated T-tubule remodeling and a Rab2 role in autophagy. eLife 6, e23367 (2017).

13. H. Kajiho, Y. Kajiho, E. Frittoli, S. Confalonieri, G. Bertalot, G. Viale, P. P. Di Fiore, A. Oldani, M. Garre, G. V. Beznoussenko, A. Palamidessi, M. Vecchi, P. Chavrier, F. Perez, G. Scita, RAB2A controls MT1-MMP endocytic and E-cadherin polarized Golgi trafficking to promote invasive breast cancer programs. EMBO reports 17, 1061–1080 (2016).

14. T. Csizmadia, P. Lőrincz, K. Hegedűs, S. Széplaki, P. Lőw, G. Juhász, Molecular mechanisms of developmentally programmed crinophagy in Drosophila. The Journal of cell biology 217, 361–374 (2018).

15. Z. Wang, G. Miao, X. Xue, X. Guo, C. Yuan, Z. Wang, G. Zhang, Y. Chen, D. Feng, J. Hu, H. Zhang, The Vici Syndrome Protein EPG5 Is a Rab7 Effector that Determines the Fusion Specificity of Autophagosomes with Late Endosomes/Lysosomes. Molecular cell 63, 781–795 (2016).

16. P. Jiang, T. Nishimura, Y. Sakamaki, E. Itakura, T. Hatta, T. Natsume, N. Mizushima, The HOPS complex mediates autophagosome-lysosome fusion through interaction with syntaxin 17. Molecular biology of the cell 25, 1327–1337 (2014).

17. C. G. Angers, A. J. Merz, HOPS interacts with Apl5 at the vacuole membrane and is required for consumption of AP-3 transport vesicles. Molecular biology of the cell 20, 4563–4574 (2009).

18. M. S. Pols, C. ten Brink, P. Gosavi, V. Oorschot, J. Klumperman, The HOPS proteins hVps41 and hVps39 are required for homotypic and heterotypic late endosome fusion. Traffic (Copenhagen, Denmark) 14, 219–232 (2013).

19. H. J. Balderhaar, J. Lachmann, E. Yavavli, C. Brocker, A. Lurick, C. Ungermann, The CORVET complex promotes tethering and fusion of Rab5/Vps21-positive membranes. Proceedings of the National Academy of Sciences of the United States of America 110, 3823–3828 (2013).

20. R. L. Plemel, B. T. Lobingier, C. L. Brett, C. G. Angers, D. P. Nickerson, A. Paulsel, D. Sprague, A. J. Merz, Subunit organization and Rab interactions of Vps-C protein complexes that control endolysosomal membrane traffic. Molecular biology of the cell 22, 1353–1363 (2011).

21. C. Brocker, A. Kuhlee, C. Gatsogiannis, H. J. Balderhaar, C. Honscher, S. Engelbrecht-Vandre, C. Ungermann, S. Raunser, Molecular architecture of the multisubunit homotypic fusion and vacuole protein sorting (HOPS) tethering complex. Proceedings of the National Academy of Sciences of the United States of America 109, 1991–1996 (2012).

22. A. Lürick, D. Kümmel, C. Ungermann, Multisubunit tethers in membrane fusion. Current Biology 28, R417–R420 (2018).

23. D. F. Seals, G. Eitzen, N. Margolis, W. T. Wickner, A. Price, A Ypt/Rab effector complex containing the Sec1 homolog Vps33p is required for homotypic vacuole fusion. Proceedings of the National Academy of Sciences of the United States of America 97, 9402–9407 (2000).

24. A. E. Wurmser, T. K. Sato, S. D. Emr, New component of the vacuolar class C-Vps complex couples nucleotide exchange on the Ypt7 GTPase to SNARE-dependent docking and fusion. The Journal of cell biology 151, 551–562 (2000).

25. K. Peplowska, D. F. Markgraf, C. W. Ostrowicz, G. Bange, C. Ungermann, The CORVET tethering complex interacts with the yeast Rab5 homolog Vps21 and is involved in endo-lysosomal biogenesis. Developmental cell 12, 739–750 (2007).

26. S. E. Rieder, S. D. Emr, A novel RING finger protein complex essential for a late step in protein transport to the yeast vacuole. Molecular biology of the cell 8, 2307–2327 (1997).

27. R. van der Kant, C. T. Jonker, R. H. Wijdeven, J. Bakker, L. Janssen, J. Klumperman, J. Neefjes, Characterization of the mammalian CORVET and HOPS complexes and their modular restructuring for endosome specificity. The Journal of biological chemistry, (2015).

28. E. D. Perini, R. Schaefer, M. Stoter, Y. Kalaidzidis, M. Zerial, Mammalian CORVET is required for fusion and conversion of distinct early endosome subpopulations. Traffic (Copenhagen, Denmark) 15, 1366–1389 (2014).

29. J. Lachmann, E. Glaubke, P. S. Moore, C. Ungermann, The Vps39-like TRAP1 is an effector of Rab5 and likely the missing Vps3 subunit of human CORVET. Cellular logistics 4, e970840 (2014).

30. P. Lorincz, Z. Lakatos, A. Varga, T. Maruzs, Z. Simon-Vecsei, Z. Darula, P. Benko, G. Csordas, M. Lippai, I. Ando, K. Hegedus, K. F. Medzihradszky, S. Takats, G. Juhasz, MiniCORVET is a Vps8-containing early endosomal tether in Drosophila. eLife 5, e14226 (2016).

31. P. Lorincz, L. A. Kenez, S. Toth, V. Kiss, A. Varga, T. Csizmadia, Z. Simon-Vecsei, G. Juhasz, Vps8 overexpression inhibits HOPS-dependent trafficking routes by outcompeting Vps41/Lt. eLife 8, e45631 (2019).

32. D. Fasshauer, R. B. Sutton, A. T. Brunger, R. Jahn, Conserved structural features of the synaptic fusion complex: SNARE proteins reclassified as Q-and R-SNAREs. Proceedings of the national academy of sciences 95, 15781–15786 (1998).

33. R. Jahn, R. H. Scheller, SNAREs--engines for membrane fusion. Nature reviews. Molecular cell biology 7, 631–643 (2006).

34. D. Brandhorst, D. Zwilling, S. O. Rizzoli, U. Lippert, T. Lang, R. Jahn, Homotypic fusion of early endosomes: SNAREs do not determine fusion specificity. Proceedings of the National Academy of Sciences 103, 2701–2706 (2006).

35. R. J. Advani, H.-R. Bae, J. B. Bock, D. S. Chao, Y.-C. Doung, R. Prekeris, J.-S. Yoo, R. H. Scheller, Seven Novel Mammalian SNARE Proteins Localize to Distinct Membrane Compartments*. Journal of Biological Chemistry 273, 10317–10324 (1998).

36. W. Antonin, C. Holroyd, R. Tikkanen, S. Höning, R. Jahn, The R-SNARE Endobrevin/VAMP-8 Mediates Homotypic Fusion of Early Endosomes and Late Endosomes. Molecular biology of the cell 11, 3289–3298 (2000).

37. S. H. Wong, Y. Xu, T. Zhang, W. Hong, Syntaxin 7, a Novel Syntaxin Member Associated with the Early Endosomal Compartment*. Journal of Biological Chemistry 273, 375–380 (1998).

38. P. R. Pryor, B. M. Mullock, N. A. Bright, M. R. Lindsay, S. R. Gray, S. C. Richardson, A. Stewart, D. E. James, R. C. Piper, J. P. Luzio, Combinatorial SNARE complexes with VAMP7 or VAMP8 define different late endocytic fusion events. EMBO reports 5, 590–595 (2004).

39. R. J. Advani, B. Yang, R. Prekeris, K. C. Lee, J. Klumperman, R. H. Scheller, Vamp-7 Mediates Vesicular Transport from Endosomes to Lysosomes. Journal of Cell Biology 146, 765–776 (1999).

40. I. Dingjan, P. T. A. Linders, D. R. J. Verboogen, N. H. Revelo, M. t. Beest, G. v. d. Bogaart, Endosomal and Phagosomal SNAREs. Physiological Reviews 98, 1465–1492 (2018).

41. V. Atlashkin, V. Kreykenbohm, E. L. Eskelinen, D. Wenzel, A. Fayyazi, G. Fischer von Mollard, Deletion of the SNARE vti1b in mice results in the loss of a single SNARE partner, syntaxin 8. Molecular and cellular biology 23, 5198–5207 (2003).

42. C. Bollmann, S. Schöning, K. Kotschnew, J. Grosse, N. Heitzig, G. Fischer von Mollard, Primary neurons lacking the SNAREs vti1a and vti1b show altered neuronal development. Neural Development 17, 12 (2022).

43. A. J. Kunwar, M. Rickmann, B. Backofen, S. M. Browski, J. Rosenbusch, S. Schöning, T. Fleischmann, K. Krieglstein, G. Fischer von Mollard, Lack of the endosomal SNAREs vti1a and vti1b led to significant impairments in neuronal development. Proceedings of the National Academy of Sciences 108, 2575–2580 (2011).

44. A. Music, B. Tejeda-González, D. M. Cunha, G. Fischer von Mollard, S. Hernández-Pérez, P. K. Mattila, The SNARE protein Vti1b is recruited to the sites of BCR activation but is redundant for antigen internalisation, processing and presentation. Frontiers in Cell and Developmental Biology **Volume** 10 -2022, (2022).

45. T. Matsui, P. Jiang, S. Nakano, Y. Sakamaki, H. Yamamoto, N. Mizushima, Autophagosomal YKT6 is required for fusion with lysosomes independently of syntaxin 17. The Journal of cell biology 217, 2633–2645 (2018).

46. F. Jian, S. Wang, R. Tian, Y. Wang, C. Li, Y. Li, S. Wang, C. Fang, C. Ma, Y. Rong, The STX17-SNAP47-VAMP7/VAMP8 complex is the default SNARE complex mediating autophagosome–lysosome fusion. Cell research, 1–18 (2024).

47. L. J. Davis, N. A. Bright, J. R. Edgar, M. D. J. Parkinson, L. Wartosch, J. Mantell, A. A. Peden, J. P. Luzio, Organelle tethering, pore formation and SNARE compensation in the late endocytic pathway. Journal of cell science 134, (2021).

48. T. Fujiwara, T. Mishima, T. Kofuji, T. Chiba, K. Tanaka, A. Yamamoto, K. Akagawa, Analysis of Knock-Out Mice to Determine the Role of HPC-1/Syntaxin 1A in Expressing Synaptic Plasticity. The Journal of Neuroscience 26, 5767 (2006).

49. T. H. Kloepper, C. N. Kienle, D. Fasshauer, An elaborate classification of SNARE proteins sheds light on the conservation of the eukaryotic endomembrane system. Molecular biology of the cell 18, 3463–3471 (2007).

50. T. H. Kloepper, C. N. Kienle, D. Fasshauer, SNAREing the Basis of Multicellularity: Consequences of Protein Family Expansion during Evolution. Molecular Biology and Evolution 25, 2055–2068 (2008).

51. J. T. Littleton, A Genomic Analysis of Membrane Trafficking and Neurotransmitter Release in Drosophila. Journal of Cell Biology 150, F77–F82 (2000).

52. H. Lu, D. Bilder, Endocytic control of epithelial polarity and proliferation in Drosophila. Nature cell biology 7, 1232–1239 (2005).

53. H. A. Morrison, H. Dionne, T. E. Rusten, A. Brech, W. W. Fisher, B. D. Pfeiffer, S. E. Celniker, H. Stenmark, D. Bilder, Regulation of early endosomal entry by the Drosophila tumor suppressors Rabenosyn and Vps45. Molecular biology of the cell 19, 4167–4176 (2008).

54. E. Morelli, P. Ginefra, V. Mastrodonato, G. V. Beznoussenko, T. E. Rusten, D. Bilder, H. Stenmark, A. A. Mironov, T. Vaccari, Multiple functions of the SNARE protein Snap29 in autophagy, endocytic, and exocytic trafficking during epithelial formation in Drosophila. Autophagy, 0 (2014).

55. Y. Yamazaki, C. Schönherr, G. K. Varshney, M. Dogru, B. Hallberg, R. H. Palmer, Goliath family E3 ligases regulate the recycling endosome pathway via VAMP3 ubiquitylation. The EMBO journal 32, 524–537 (2013).

56. G. Szenci, G. Glatz, S. Takáts, G. Juhász, The Ykt6–Snap29–Syx13 SNARE complex promotes crinophagy via secretory granule fusion with Lamp1 carrier vesicles. Scientific reports 14, 3200 (2024).

57. Y. Lu, Z. Zhang, D. Sun, S. T. Sweeney, F. B. Gao, Syntaxin 13, a genetic modifier of mutant CHMP2B in frontotemporal dementia, is required for autophagosome maturation. Molecular cell 52, 264–271 (2013).

58. M. A. Akbar, S. Ray, H. Kramer, The SM protein Car/Vps33A regulates SNARE-mediated trafficking to lysosomes and lysosome-related organelles. Molecular biology of the cell 20, 1705–1714 (2009).

59. S. Takats, P. Nagy, A. Varga, K. Pircs, M. Karpati, K. Varga, A. L. Kovacs, K. Hegedus, G. Juhasz, Autophagosomal Syntaxin17-dependent lysosomal degradation maintains neuronal function in Drosophila. The Journal of cell biology 201, 531–539 (2013).

60. S. Takats, K. Pircs, P. Nagy, A. Varga, M. Karpati, K. Hegedus, H. Kramer, A. L. Kovacs, M. Sass, G. Juhasz, Interaction of the HOPS complex with Syntaxin 17 mediates autophagosome clearance in Drosophila. Molecular biology of the cell 25, 1338–1354 (2014).

61. Z. Lakatos, P. Lőrincz, Z. Szabó, P. Benkő, L. A. Kenéz, T. Csizmadia, G. Juhász, Sec20 is Required for Autophagic and Endocytic Degradation Independent of Golgi-ER Retrograde Transport. Cells 8, (2019).

62. S. Takats, G. Glatz, G. Szenci, A. Boda, G. V. Horvath, K. Hegedus, A. L. Kovacs, G. Juhasz, Non-canonical role of the SNARE protein Ykt6 in autophagosome-lysosome fusion. PLoS genetics 14, e1007359 (2018).

63. L. Zhou, X. Xue, K. Yang, Z. Feng, M. Liu, J. C. Pastor-Pareja, Convergence of secretory, endosomal, and autophagic routes in trans-Golgi-associated lysosomes. The Journal of cell biology 222, (2023).

64. T. Csizmadia, A. Dósa, E. Farkas, B. V. Csikos, E. A. Kriska, G. Juhász, P. Lőw, Developmental program-independent secretory granule degradation in larval salivary gland cells of Drosophila. Traffic (Copenhagen, Denmark) 23, 568–586 (2022).

65. K. Linnemannstöns, L. Witte, P. Karuna M, J. C. Kittel, A. Danieli, D. Müller, L. Nitsch, M. Honemann-Capito, F. Grawe, A. Wodarz, Ykt6-dependent endosomal recycling is required for Wnt secretion in the Drosophila wing epithelium. *Development (Cambridge*, England) 147, dev185421 (2020).

66. S. Fujii, K. Kurokawa, R. Inaba, N. Hiramatsu, T. Tago, Y. Nakamura, A. Nakano, T. Satoh, A. K. Satoh, Recycling endosomes attach to the trans-side of Golgi stacks in Drosophila and mammalian cells. Journal of cell science 133, (2020).

67. D. M. Ward, J. Pevsner, M. A. Scullion, M. Vaughn, J. Kaplan, Syntaxin 7 and VAMP-7 are soluble N-ethylmaleimide-sensitive factor attachment protein receptors required for late endosome-lysosome and homotypic lysosome fusion in alveolar macrophages. Molecular biology of the cell 11, 2327–2333 (2000).

68. D. Hargitai, A. Nagy, I. Bodor, G. Szenci, H. Laczkó-Dobos, A. Bhattacharjee, N. Neuhauser, S. Takáts, G. Juhász, P. Lőrincz, HOPS-dependent vesicle tethering lock inhibits endolysosomal fusions and autophagosome secretion upon the loss of Syntaxin17. Sci Adv 11, eadu9605 (2025).

69. A. Boda, P. Lőrincz, S. Takáts, T. Csizmadia, S. Tóth, A. L. Kovács, G. Juhász, Drosophila Arl8 is a general positive regulator of lysosomal fusion events. Biochimica et Biophysica Acta (BBA)-Molecular Cell Research 1866, 533–544 (2019).

70. V. K. Lund, K. L. Madsen, O. Kjaerulff, Drosophila Rab2 controls endosome-lysosome fusion and LAMP delivery to late endosomes. Autophagy 14, 1520–1542 (2018).

71. Y. Fu, J. Y. Zhu, F. Zhang, A. Richman, Z. Zhao, Z. Han, Comprehensive functional analysis of Rab GTPases in Drosophila nephrocytes. Cell Tissue Res 368, 615–627 (2017).

72. T. Maruzs, D. Feil-Börcsök, E. Lakatos, G. Juhász, A. Blastyák, D. Hargitai, S. Jean, P. Lőrincz, G. Juhász, Interaction of the sorting nexin 25 homologue Snazarus with Rab11 balances endocytic and secretory transport and maintains the ultrafiltration diaphragm in nephrocytes. Molecular biology of the cell 34, ar87 (2023).

73. H. Sheng, W. Stauffer, H. N. Lim, Systematic and general method for quantifying localization in microscopy images. Biology open 5, 1882–1893 (2016).

74. K. Lang, J. Milosavljevic, H. Heinkele, M. Chen, L. Gerstner, D. Spitz, S. Kayser, M. Helmstädter, G. Walz, M. Köttgen, A. Spracklen, J. Poulton, T. Hermle, Selective endocytosis controls slit diaphragm maintenance and dynamics in Drosophila nephrocytes. eLife 11, e79037 (2022).

75. E. Itakura, C. Kishi-Itakura, N. Mizushima, The hairpin-type tail-anchored SNARE syntaxin 17 targets to autophagosomes for fusion with endosomes/lysosomes. Cell 151, 1256–1269 (2012).

76. S. Koike, R. Jahn, SNAREs define targeting specificity of trafficking vesicles by combinatorial interaction with tethering factors. Nature communications 10, 1608 (2019).

77. T. Vaccari, H. Lu, R. Kanwar, M. E. Fortini, D. Bilder, Endosomal entry regulates Notch receptor activation in Drosophila melanogaster. The Journal of cell biology 180, 755–762 (2008).

78. J. J. Smith, K. Sumiyama, C. T. Amemiya, A living fossil in the genome of a living fossil: Harbinger transposons in the coelacanth genome. Mol Biol Evol 29, 985–993 (2012).

79. M. Plooster, G. Rossi, M. S. Farrell, J. C. McAfee, J. L. Bell, M. Ye, G. H. Diering, H. Won, S. L. Gupton, P. Brennwald, Schizophrenia-Linked Protein tSNARE1 Regulates Endosomal Trafficking in Cortical Neurons. The Journal of neuroscience : the official journal of the Society for Neuroscience 41, 9466–9481 (2021).

80. P. Wen, F. Zhang, Y. Fu, J. Y. Zhu, Z. Han, Exocyst Genes Are Essential for Recycling Membrane Proteins and Maintaining Slit Diaphragm in Drosophila Nephrocytes. Journal of the American Society of Nephrology : JASN 31, 1024–1034 (2020).

81. H. Weavers, S. Prieto-Sanchez, F. Grawe, A. Garcia-Lopez, R. Artero, M. Wilsch-Brauninger, M. Ruiz-Gomez, H. Skaer, B. Denholm, The insect nephrocyte is a podocyte-like cell with a filtration slit diaphragm. Nature 457, 322–326 (2009).

82. S. Zhuang, H. Shao, F. Guo, R. Trimble, E. Pearce, S. M. Abmayr, Sns and Kirre, the Drosophila orthologs of Nephrin and Neph1, direct adhesion, fusion and formation of a slit diaphragm-like structure in insect nephrocytes. *Development (Cambridge*, England) 136, 2335–2344 (2009).

83. F. Zhang, Y. Zhao, Y. Chao, K. Muir, Z. Han, Cubilin and amnionless mediate protein reabsorption in Drosophila nephrocytes. Journal of the American Society of Nephrology : JASN 24, 209–216 (2013).

84. D. Zheng, M. Tong, S. Zhang, Y. Pan, Y. Zhao, Q. Zhong, X. Liu, Human YKT6 forms priming complex with STX17 and SNAP29 to facilitate autophagosome-lysosome fusion. Cell reports 43, (2024).

85. D. M. Dean, L. E. Codd, R. Constanza, X. M. Segel, purpleoid1, a classic Drosophila eye color mutation, is an allele of the t-SNARE-encoding gene SNAP29. microPublication Biology 2025, (2025).

86. D. Fuchs-Telem, H. Stewart, D. Rapaport, J. Nousbeck, A. Gat, M. Gini, Y. Lugassy, S. Emmert, K. Eckl, H. C. Hennies, O. Sarig, D. Goldsher, B. Meilik, A. Ishida-Yamamoto, M. Horowitz, E. Sprecher, CEDNIK syndrome results from loss-of-function mutations in SNAP29. The British journal of dermatology 164, 610–616 (2011).

87. E. Sprecher, A. Ishida-Yamamoto, M. Mizrahi-Koren, D. Rapaport, D. Goldsher, M. Indelman, O. Topaz, I. Chefetz, H. Keren, J. O’Brien T, D. Bercovich, S. Shalev, D. Geiger, R. Bergman, M. Horowitz, H. Mandel, A mutation in SNAP29, coding for a SNARE protein involved in intracellular trafficking, causes a novel neurocutaneous syndrome characterized by cerebral dysgenesis, neuropathy, ichthyosis, and palmoplantar keratoderma. American journal of human genetics 77, 242–251 (2005).

88. S. Koike, R. Jahn, SNARE proteins: zip codes in vesicle targeting? The Biochemical journal 479, 273–288 (2022).

89. S. Koike, R. Jahn, Rab GTPases and phosphoinositides fine-tune SNAREs dependent targeting specificity of intracellular vesicle traffic. Nature communications 15, 2508 (2024).

90. D. Yadav, A. Hacisuleyman, M. Dergai, D. Khalifeh, L. A. Abriata, M. D. Peraro, D. Fasshauer, A look beyond the QR code of SNARE proteins. Protein science : a publication of the Protein Society 33, e5158 (2024).

91. M. L. Schwartz, D. P. Nickerson, B. T. Lobingier, R. L. Plemel, M. Duan, C. G. Angers, M. Zick, A. J. Merz, Sec17 (alpha-SNAP) and an SM-tethering complex regulate the outcome of SNARE zippering in vitro and in vivo. eLife 6, (2017).

92. A. Orr, H. Song, S. F. Rusin, A. N. Kettenbach, W. Wickner, HOPS catalyzes the interdependent assembly of each vacuolar SNARE into a SNARE complex. Molecular biology of the cell 28, 975–983 (2017).

93. H. Song, A. S. Orr, M. Lee, M. E. Harner, W. T. Wickner, HOPS recognizes each SNARE, assembling ternary trans-complexes for rapid fusion upon engagement with the 4th SNARE. eLife 9, e53559 (2020).

94. V. L. Koumandou, B. Wickstead, M. L. Ginger, M. van der Giezen, J. B. Dacks, M. C. Field, Molecular paleontology and complexity in the last eukaryotic common ancestor. Critical reviews in biochemistry and molecular biology 48, 373–396 (2013).

95. G. Glatz, G. Gogl, A. Alexa, A. Remenyi, Structural mechanism for the specific assembly and activation of the extracellular signal regulated kinase 5 (ERK5) module. The Journal of biological chemistry 288, 8596–8609 (2013).

96. A. Atienza-Manuel, V. Castillo-Mancho, S. De Renzis, J. Culi, M. Ruiz-Gómez, Endocytosis mediated by an atypical CUBAM complex modulates slit diaphragm dynamics in nephrocytes. Development (Cambridge, England) 148, (2021).

97. N. Chaudhry, M. Sica, S. Surabhi, D. S. Hernandez, A. Mesquita, A. Selimovic, A. Riaz, L. Lescat, H. Bai, G. C. MacIntosh, A. Jenny, Lamp1 mediates lipid transport, but is dispensable for autophagy in Drosophila. Autophagy 18, 2443–2458 (2022).

98. F. Hochapfel, L. Denk, G. Mendl, U. Schulze, C. Maaßen, Y. Zaytseva, H. Pavenstädt, T. Weide, R. Rachel, R. Witzgall, Distinct functions of Crumbs regulating slit diaphragms and endocytosis in Drosophila nephrocytes. Cellular and Molecular Life Sciences 74, 4573–4586 (2017).

99. S. Jean, S. Cox, S. Nassari, A. A. Kiger, Starvation-induced MTMR13 and RAB21 activity regulates VAMP8 to promote autophagosome–lysosome fusion. EMBO reports 16, 297–311 (2015).

100. A. Öztürk-Çolak, S. J. Marygold, G. Antonazzo, H. Attrill, D. Goutte-Gattat, V. K. Jenkins, B. B. Matthews, G. Millburn, G. dos Santos, C. J. Tabone, C. FlyBase, FlyBase: updates to the Drosophila genes and genomes database. Genetics 227, iyad211 (2024).

101. J. Milosavljevic, C. Lempicki, K. Lang, H. Heinkele, L. L. Kampf, C. Leroy, M. Chen, L. Gerstner, D. Spitz, M. Wang, A. U. Knob, S. Kayser, M. Helmstädter, G. Walz, M. R. Pollak, T. Hermle, Nephrotic Syndrome Gene TBC1D8B Is Required for Endosomal Maturation and Nephrin Endocytosis in Drosophila. Journal of the American Society of Nephrology 33, 2174–2193 (2022).

102. T. Kosaka, K. Ikeda, Reversible blockage of membrane retrieval and endocytosis in the garland cell of the temperature-sensitive mutant of Drosophila melanogaster, shibirets1. The Journal of cell biology 97, 499–507 (1983).

