## Supplemental File for "Multiple overlapping SNARE complexes drive endosome maturation in Drosophila nephrocytes"

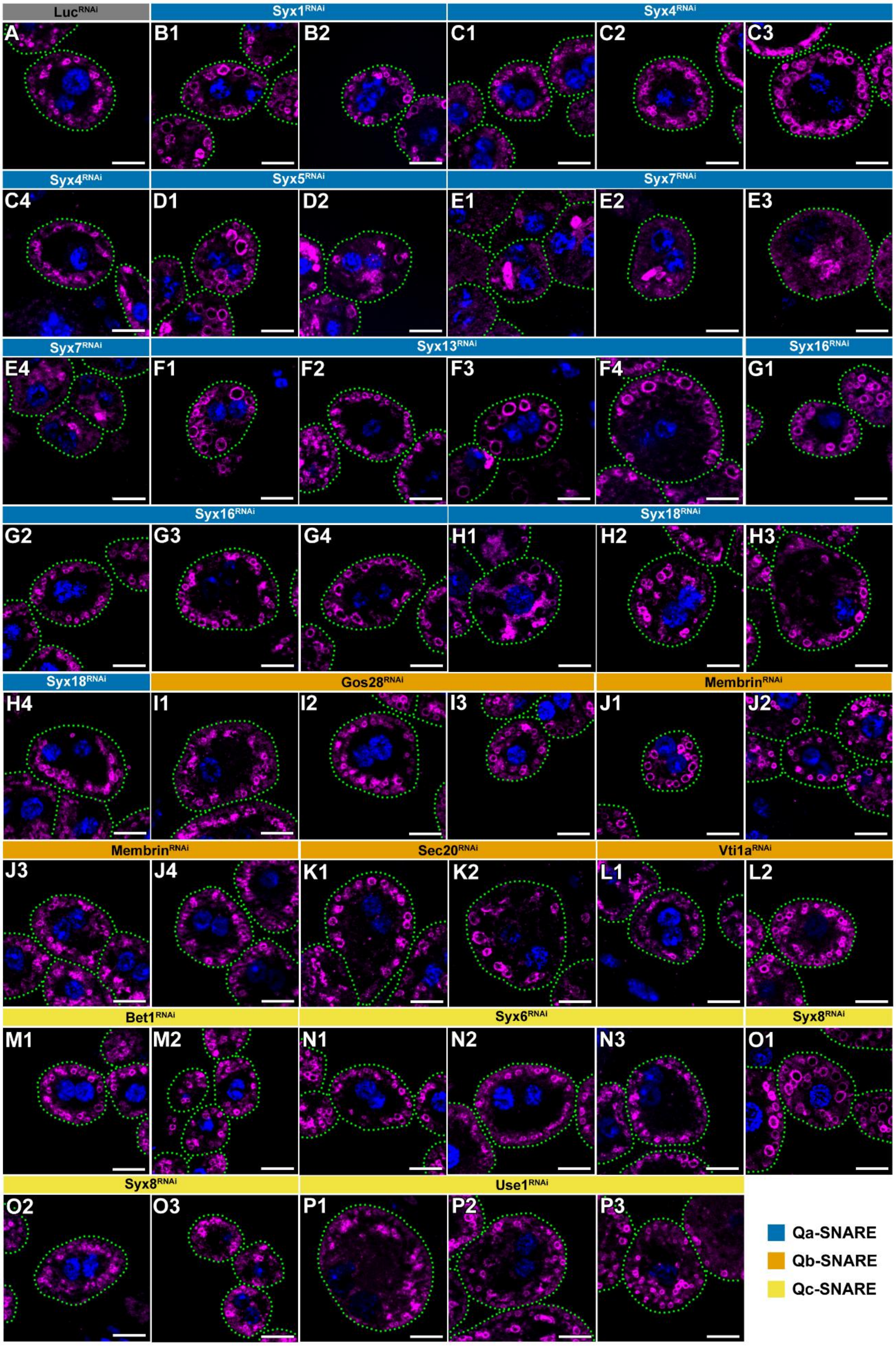

Luc RNAi
Syx1 RNAi
Syx4 RNAi
Syx5 RNAi
Syx7 RNAi
Syx13 RNAi
Syx16 RNAi
Syx18 RNAi
Gos28 RNAi
Membrin RNAi
Sec20 RNAi
Vti1a RNAi
Bet1 RNAi
Syx6 RNAi
Syx8 RNAi
Use1 RNAi
A
B1
B2
C1
C2
C3
C4
D1
D2
E1
E2
E3
E4
F1
F2
F3
F4
G1
G2
G3
G4
H1
H2
H3
H4
I1
I2
I3
J1
J2
J3
J4
K1
K2
L1
L2
M1
M2
N1
N2
N3
O1
O2
O3
P1
P2
P3
Qa-SNARE
Qb-SNARE
Qc-SNARE

**Figure S1. Representative micrographs of the SNARE RNAi screen: Qa, Qb, and Qc SNAREs**

A1–P3) Representative images from independent RNAi lines targeting Qa, Qb, and Qc SNAREs. Magenta and blue channels show $Rab7^+$ late endosomes and nuclei (DAPI), respectively. Cell outlines are indicated by dashed cyan lines. Scale bars: 10 µm.

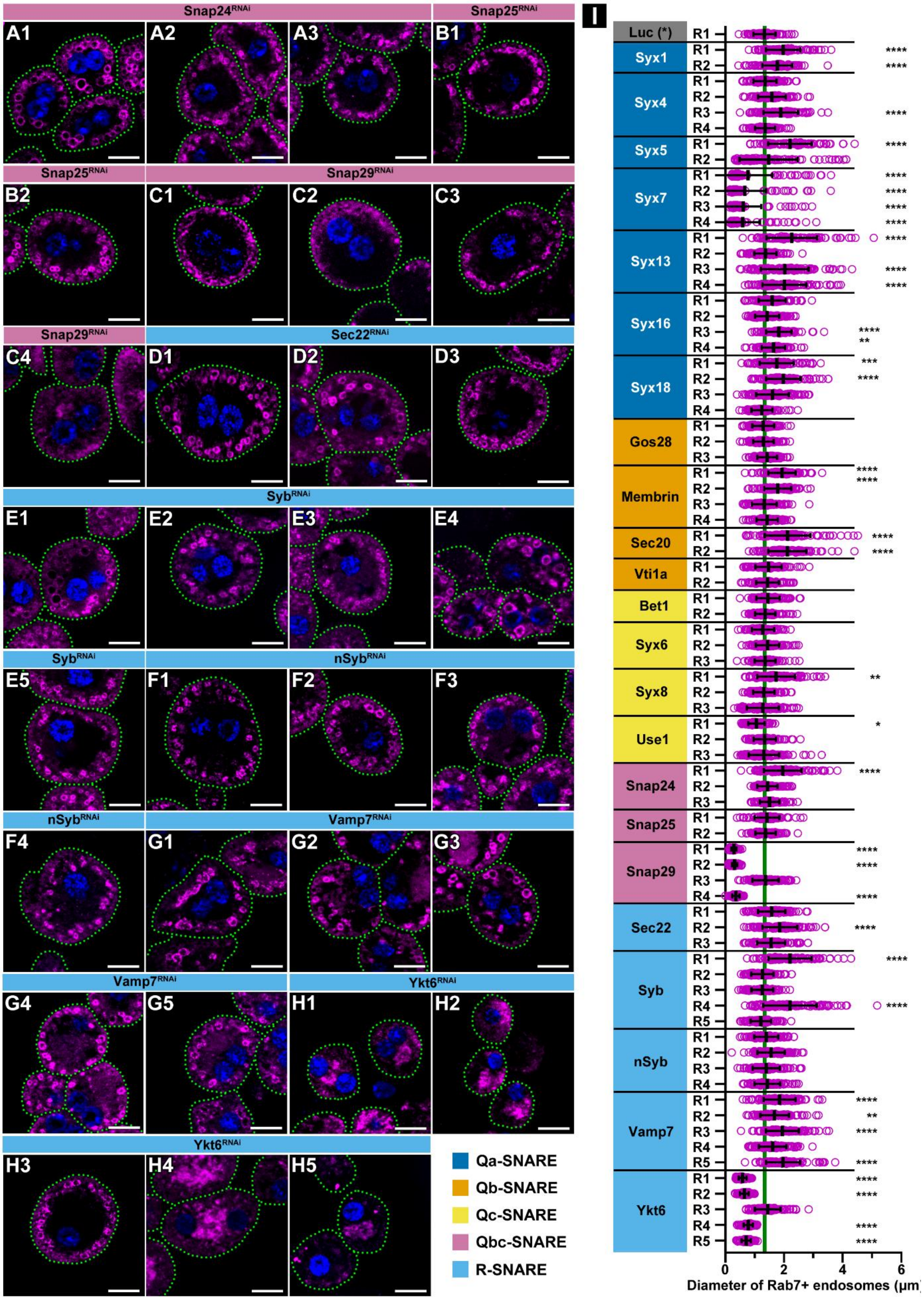

Snap24RNAi
Snap25RNAi
A1
A2
A3
B1
Snap25RNAi
Snap29RNAi
B2
C1
C2
C3
Snap29RNAi
Sec22RNAi
C4
D1
D2
D3
SybRNAi
E1
E2
E3
E4
SybRNAi
nSybRNAi
E5
F1
F2
F3
nSybRNAi
Vamp7RNAi
F4
G1
G2
G3
Vamp7RNAi
Ykt6RNAi
G4
G5
H1
H2
Ykt6RNAi
H3
H4
H5
Qa-SNARE
Qb-SNARE
Qc-SNARE
Qbc-SNARE
R-SNARE
I
Luc (*)
Syx1
Syx4
Syx5
Syx7
Syx13
Syx16
Syx18
Gos28
Membrin
Sec20
Vti1a
Bet1
Syx6
Syx8
Use1
Snap24
Snap25
Snap29
Sec22
Syb
nSyb
Vamp7
Ykt6
R1
R2
R3
R4
R5
0
2
4
6
Diameter of Rab7+ endosomes (μm)

**Figure S2. Representative micrographs and quantification of the SNARE RNAi screen: Qbc and R SNAREs**

A1–H5) Representative images from independent RNAi lines targeting Qbc and R SNAREs (continuation of Figure S1). Magenta and blue channels show $Rab7^+$ late endosomes and nuclei (DAPI), respectively. Cell outlines are indicated by dashed cyan lines. Scale bars: 10 µm.

I) Quantification of $Rab7^+$ endosome size shown in Figure S1 and A1–H5. N = 100 endosomes from 10 cells. Asterisks indicate comparisons to control (Luciferase RNAi, Fig. S1A). **** $p < 0.0001$; ** $p < 0.01$; * $p < 0.05$. Dashed line indicates mean endosome size in control cells.

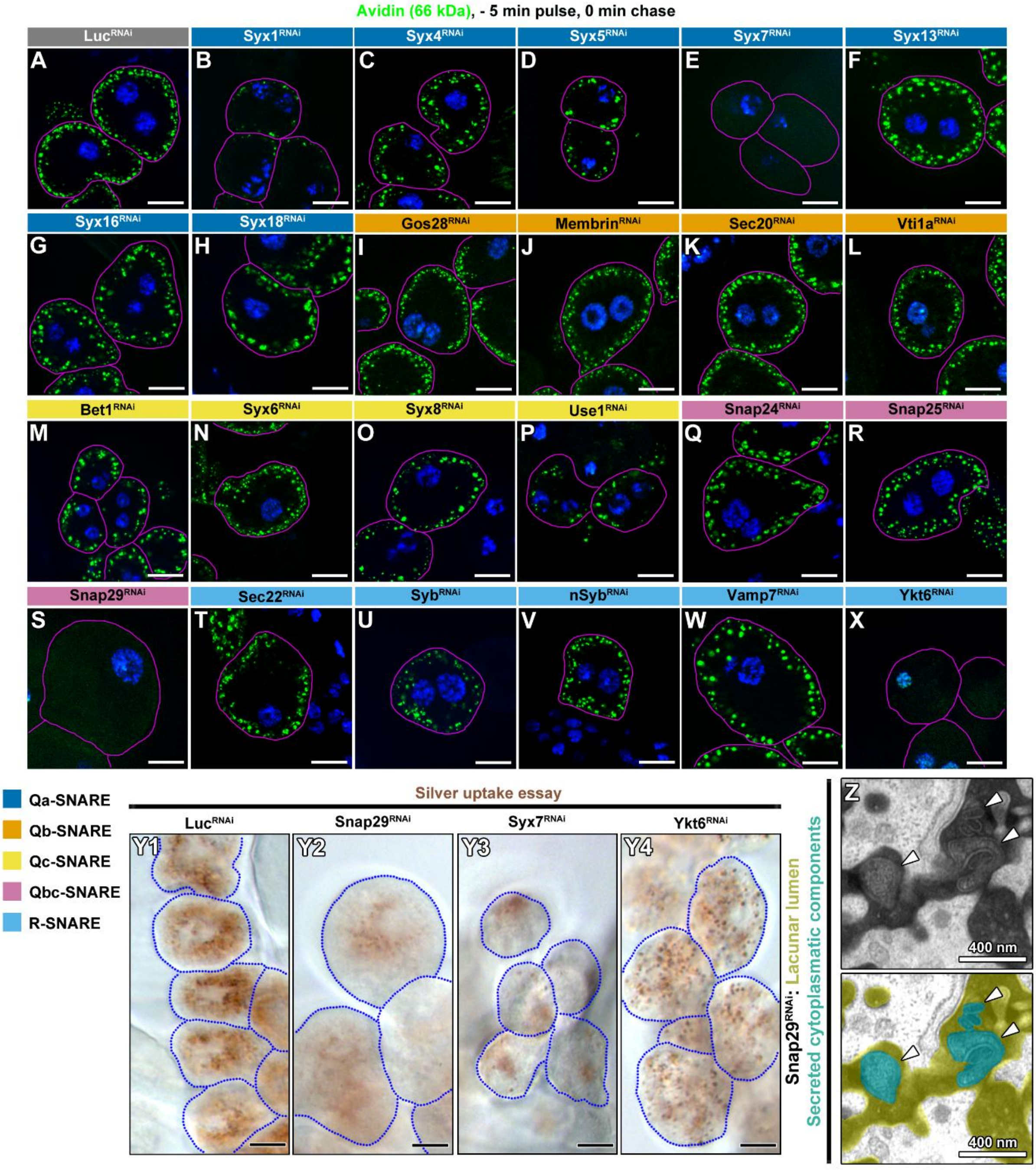

Avidin (66 kDa), - 5 min pulse, 0 min chase
Luc RNAi
Syx1 RNAi
Syx4 RNAi
Syx5 RNAi
Syx7 RNAi
Syx13 RNAi
Syx16 RNAi
Syx18 RNAi
Gos28 RNAi
Membrin RNAi
Sec20 RNAi
Vti1a RNAi
Bet1 RNAi
Syx6 RNAi
Syx8 RNAi
Use1 RNAi
Snap24 RNAi
Snap25 RNAi
Snap29 RNAi
Sec22 RNAi
Syb RNAi
nSyb RNAi
Vamp7 RNAi
Ykt6 RNAi
A
B
C
D
E
F
G
H
I
J
K
L
M
N
O
P
Q
R
S
T
U
V
W
X
Qa-SNARE
Qb-SNARE
Qc-SNARE
Qbc-SNARE
R-SNARE
Silver uptake essay
Luc RNAi
Snap29 RNAi
Syx7 RNAi
Ykt6 RNAi
Y1
Y2
Y3
Y4
Snap29 RNAi: Lacunar lumen
Secreted cytoplasmatic components
Z
400 nm
400 nm

**Figure S3: Additional nephrocyte data (uptake and TEM)**

A-X) Representative images from RNAi-based avidin uptake screen. Green and blue channels show internalized avidin and nuclei (DAPI), respectively. Cell outlines are indicated by magenta lines. Scale bars: 10 µm.

Y) Silver nitrate uptake and transport to lysosomes proceed normally in control nephrocytes (Y1), as indicated by brown cytosolic silver inclusions. In contrast, internalized silver is reduced in Syx7 (Y2), Snap29 (Y3), and Ykt6 (Y4) RNAi cells, although inclusions remain detectable in Ykt6 RNAi cells (Y4). Scale bars: 10 µm.

Z) Higher-magnification view of the lacunar lumen in Snap29 RNAi nephrocytes shown in Fig. 5C. The lumen is marked by electron-dense TA, and secreted autophagosomes (arrowheads) are visible.

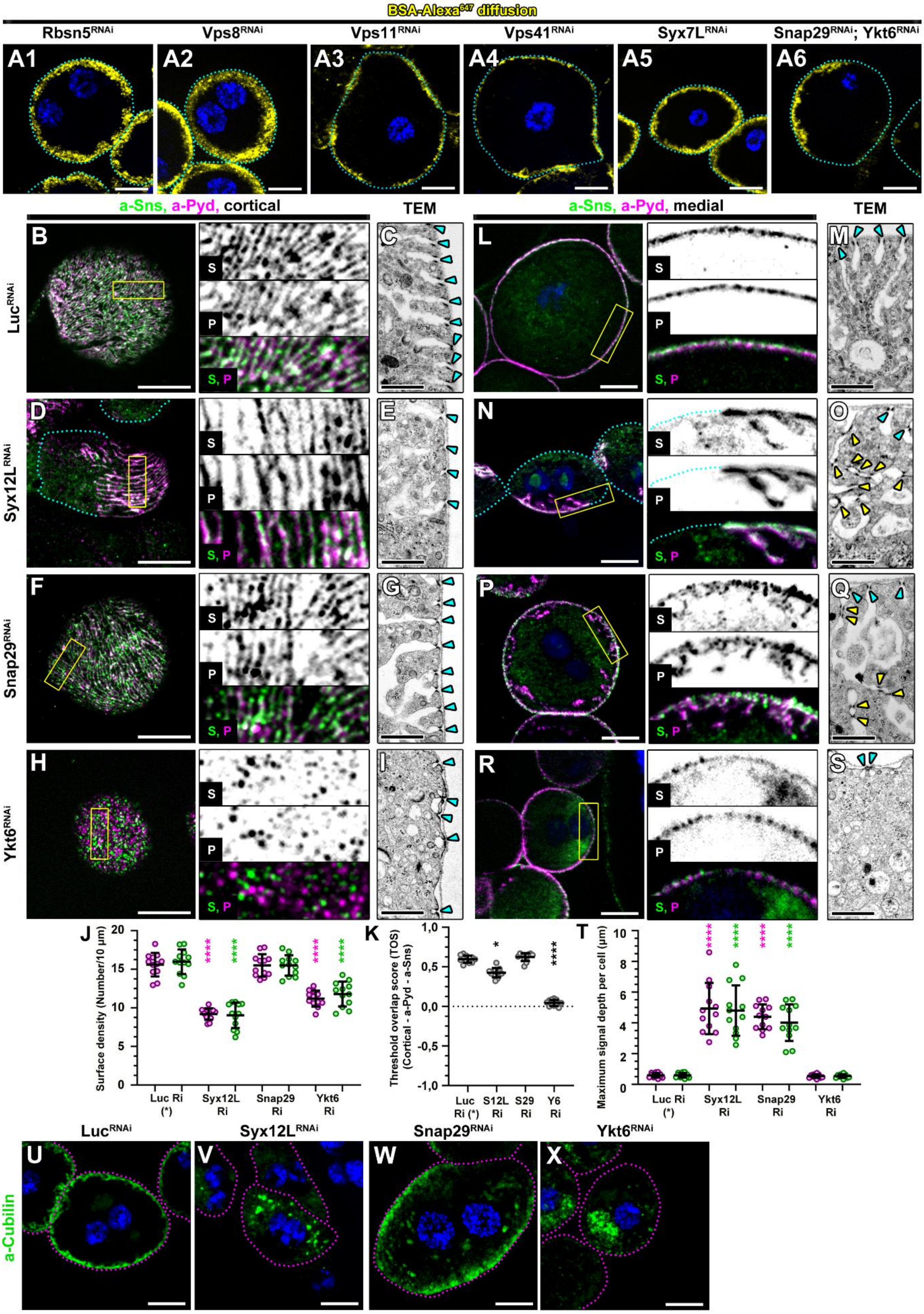

BSA-Alexa647 diffusion
Rbsn5RNAi
Vps8RNAi
Vps11RNAi
Vps41RNAi
Syx7LRNAi
Snap29RNAi; Ykt6RNAi
A1
A2
A3
A4
A5
A6
a-Sns, a-Pyd, cortical
TEM
a-Sns, a-Pyd, medial
TEM
LucRNAi
Syx12LRNAi
Snap29RNAi
Ykt6RNAi
S
P
S, P
Surface density (Number/10 μm)
Luc Ri (*)
Syx12L Ri
Snap29 Ri
Ykt6 Ri
Threshold overlap score (TOS) (Cortical - a-Pyd - a-Sns)
Luc Ri (*)
S12L Ri
S29 Ri
Y6 Ri
Maximum signal depth per cell (μm)
LucRNAi
Syx12LRNAi
Snap29RNAi
Ykt6RNAi
a-Cubilin

**Figure S4: The filtration system is defective in Syx12L, Snap29 and Ykt6 depleted nephrocytes, and additional lacunar depth data**

A1–A6) Channel diffusion assay using BSA–Alexa647 shows increased lacunar depth in early endosomal tether-depleted cells (Rbsn5 RNAi, A1; Vps8 RNAi, A2) compared with control (Fig. 5F). In contrast, lacunar depth is reduced in HOPS subunit RNAi cells (Vps11, A3; Vps41, A4). In Syx7L RNAi cells (A5); formerly Syx13), lacunar depth remains similar to control. In Snap29/Ykt6 double RNAi cells (A6) lacunar depth is reduced, similar to Ykt6 single RNAi (Fig. 5I). Blue: nuclei (DAPI). Cell outlines are indicated by dashed cyan lines.

B–S) Slit diaphragm (SD) morphology was examined by fluorescence microscopy in nephrocytes immunostained for the SD protein Sticks and stones (Sns) and the SD adaptor Polychaetoid (Pyd), and by transmission electron microscopy (TEM). Cortical sections (B–I) show SD patterning at the cell surface, and medial sections (L–S) show cross-sectional views of SD units and deeper regions of nephrocytes. Cyan arrowheads indicate cortical SDs, and yellow arrowheads indicate ectopic SDs. In controls (B, C, L, M), only cortical SDs are present, exhibiting a characteristic fingerprint-like pattern in which Sns and Pyd colocalize. In all three examined SNARE RNAi conditions, SD morphology is dramatically altered (D–I, N–S), although in distinct ways.

The fingerprint-like pattern becomes irregular in Syx12L (D, E) and Snap29 (F, G) RNAi cells, and ectopic SDs appear as intracellular Sns- and Pyd-positive protrusions in Syx12L (N) and Snap29 (P) RNAi cells, also detected by TEM (O, Q). Note that the density of cortical SDs is reduced in Syx12L RNAi cells (D). In contrast, the fingerprint-like SD pattern is completely abolished in Ykt6 RNAi cells (H, I), leaving an irregular dot-like distribution in which Sns and Pyd show minimal colocalization. Ectopic SDs are not observed (R, S). Note the absence of lacunar invaginations in Ykt6-depleted cells (I, S). Blue: nuclei (DAPI). Cell outlines are indicated by cyan lines in D and N. Scale bars: 10 μm (fluorescence) and 1 μm (TEM).

J) Quantification of the surface density of SD components shown in A, C, E, and G reveals reduced cortical SDs in Syx12L and Ykt6 RNAi cells. Asterisks (*) indicate comparisons to control. **** $p < 0.0001$.

K) Quantification of colocalization of cortical SD components using the Threshold Overlap Score (TOS) from A, C, E, and G shows a slight decrease in Syx12L and a dramatic decrease in Ykt6 RNAi cells. Asterisks (*) indicate comparisons to control. $p < 0.05$; **** $p < 0.0001$.

T) Quantification of SD signal depth from K, M, O, and Q shows ectopic SDs in Syx12L and Snap29 RNAi cells. Asterisks (*) indicate comparisons to control. **** $p < 0.0001$.

U–X) Lacunar morphology examined by Cubilin immunostaining reveals increased lacunar channel depth in Syx12L (V) and Snap29 (W) RNAi cells compared with controls (U). Note the intense internal Cubilin signal in Syx12L cells (V), corresponding to lacunar dilations observed in Fig. 5B, G. In contrast, in Ykt6 RNAi cells (X) the cortical Cubilin signal is negligible, and Cubilin accumulates in numerous small vesicles, indicating defective transport to the plasma membrane. Blue channel shows nuclei (DAPI); cell outlines are indicated by magenta lines. Scale bars: 10 μm.

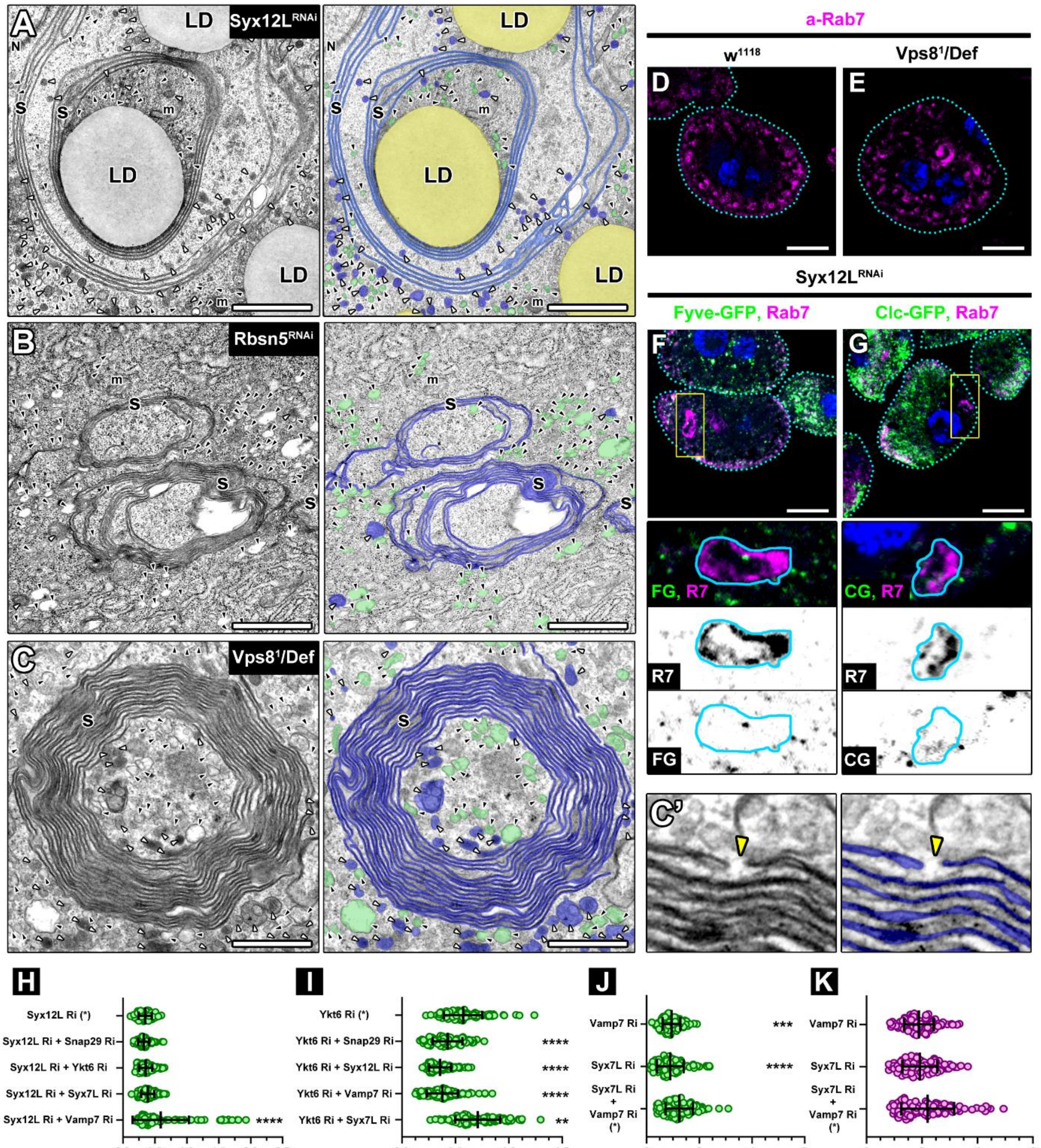

A
Syx12L RNAi
LD
N
S
m
B
Rbsn5 RNAi
C
Vps8[1]/Def
a-Rab7
D
w[1118]
E
Vps8[1]/Def
Syx12L RNAi
Fyve-GFP, Rab7
Clc-GFP, Rab7
F
G
FG, R7
CG, R7
R7
FG
CG
C'
H
Syx12L Ri (*)
Syx12L Ri + Snap29 Ri
Syx12L Ri + Ykt6 Ri
Syx12L Ri + Syx7L Ri
Syx12L Ri + Vamp7 Ri
****
Diameter of Rab5+ endosomes (µm)
I
Ykt6 Ri (*)
Ykt6 Ri + Snap29 Ri
Ykt6 Ri + Syx12L Ri
Ykt6 Ri + Vamp7 Ri
Ykt6 Ri + Syx7L Ri
****
**
Diameter of Rab5+ endosomes (µm)
J
Vamp7 Ri
Syx7L Ri
Syx7L Ri + Vamp7 Ri (*)
***
****
Diameter of Rab5+ endosomes (µm)
K
Vamp7 Ri
Syx7L Ri
Syx7L Ri + Vamp7 Ri (*)
Diameter of Rab7+ endosomes (µm)

**Figure S5: Additional characterization of endolysosomal "swirl" structures and additional quantification data to Figure 7.**

A) Representative electron micrograph of a membrane swirl (S) in a Syx12L-depleted nephrocyte. Scale bar: 1 µm. m, mitochondria; LD, lipid droplets (orange). White arrowheads indicate small lysosome-like structures (blue); black arrowheads indicate small endosomes (green).

B, C) TEM reveals swirl structures in Rbsn5 RNAi (B) and Vps8 mutant (C) nephrocytes. Scale bar: 1 µm. m, mitochondria. White arrowheads indicate small lysosome-like structures (blue); black arrowheads indicate small endosomes (green). Inset (C′) shows that the outer double membrane of the swirl has closed ends and is not fully continuous (yellow arrowheads).

D, E) Rab7-positive membrane swirls are present in Vps8 mutant nephrocytes (E) but not in controls (D). Blue: nuclei (DAPI). Cell outlines are indicated by dashed cyan lines. Scale bars: 10 µm.

F, G) Swirls in Syx12L RNAi cells are negative for early endosomal markers Fyve-GFP (F) and clathrin light chain (Clc)-GFP (G). Blue: nuclei (DAPI). Cell outlines are indicated by dashed cyan lines; swirl outlines are shown by continuous cyan lines in insets. Scale bars: 10 µm.

H) Quantification of $Rab5^+$ endosome size shown in Figure 7A, D, F, I and P. N = 100 endosomes from 10 cells. Asterisks indicate comparisons to Syx12L single RNAi. **** $p < 0.0001$.

I) Quantification of $Rab5^+$ endosome size shown in Figure 7B, E, F, J and Q. N = 100 endosomes from 10 cells. Asterisks indicate comparisons to Ykt6 single RNAi. **** $p < 0.0001$, ** $p < 0.01$.

J, K) Quantification of $Rab5^+$ (J) and $Rab7^+$(K) endosome size shown in Figure 7R-T. N = 100 endosomes from 10 cells. Asterisks indicate comparisons to Syx7L; Vamp7 double RNAi. **** $p < 0.0001$, *** $p < 0.001$.

**Table S1. Fly stocks, antibodies, and reagents used in this study.**

| Reagent or resource | Source/Details | ID |
|---|---|---|
| ***Drosophila* stocks** | | |
| Df(3L)ED211 | Kyoto Stock Center | FBst0313146 |
| Prospero-Gal4 | Bloomington Drosophila Stock Center [BDSC] | FBst0080572 |
| UAS-Bet1[GD2721] | Vienna Drosophila Resource Center (VDRC) | FBst0471070 |
| UAS-Bet1[HMJ22351] | Bloomington Drosophila Stock Center [BDSC] | FBst0058269 |
| UAS-Dcr-2.D | Bloomington Drosophila Stock Center [BDSC] | FBst0024650 |
| UAS-EGFP-Clc | Bloomington Drosophila Stock Center [BDSC] | FBst0007109 |
| UAS-GFP-myc-2xFYVE | Bloomington Drosophila Stock Center [BDSC] | FBst0042712 |
| UAS-GFP-Rab5 | Bloomington Drosophila Stock Center [BDSC] | FBst0043336 |
| UAS-GFP-VAMP7 | Kindly provided by Amy Kiger (*1*) | FBgn0266186 |
| UAS-Gos28[GD3051] | Vienna Drosophila Resource Center (VDRC) | FBst0450412 |
| UAS-Gos28[HMS01203] | Bloomington Drosophila Stock Center [BDSC] | FBst0034724 |
| UAS-Gos28[KK107479] | Vienna Drosophila Resource Center (VDRC) | FBst0472163 |
| UAS-hsp-Hsap\STX12.HA.1 | Kyoto Stock Center | - |
| UAS-hsp-Hsap\STX7.HA.1 | Kyoto Stock Center | - |
| UAS-Luciferase[JF01355] | Bloomington Drosophila Stock Center [BDSC] | FBst0031603 |
| UAS-Membrin[4780R-3] | National Institute of Genetics Fly Stocks (NIG-FLY) Mishima, Japan | - |
| UAS-Membrin[GD2313] | Vienna Drosophila Resource Center (VDRC) | FBst0465629 |
| UAS-Membrin[GLC01633] | Discontinued | FBtp0090271 |
| UAS-Membrin[KK100283] | Vienna Drosophila Resource Center (VDRC) | FBst0481093 |
| UAS-nSyb[GD17382] | Vienna Drosophila Resource Center (VDRC) | FBst0468331 |

| | | |
|---|---|---|
| UAS-nSyb[GD17382] | Vienna Drosophila Resource Center (VDRC) | FBst0468332 |
| UAS-nSyb[JF03417] | Bloomington Drosophila Stock Center [BDSC] | FBst0031983 |
| UAS-nSyb[KK109893] | Vienna Drosophila Resource Center (VDRC) | FBst0476389 |
| UAS-pHluorinSE-Rab2 | Kindly provided by Ole Kjaerulff (*2*) | FBal0346047 |
| UASp-YFP.Rab5 | Bloomington Drosophila Stock Center [BDSC] | FBst0009775 |
| UASp-YFP.Rab5.Q88L | Bloomington Drosophila Stock Center [BDSC] | FBst0009774 |
| UAS-Rab5[GL01872] | Bloomington Drosophila Stock Center [BDSC] | FBst0067877 |
| UAS-Rab7[JF02377] | Bloomington Drosophila Stock Center [BDSC] | FBst0027051 |
| UAS-Rbsn5[HMC04769] | Bloomington Drosophila Stock Center [BDSC] | FBst0057459 |
| UAS-Sec20[HMS01172] | Bloomington Drosophila Stock Center [BDSC] | FBst0034693 |
| UAS-Sec20[KK107334] | Vienna Drosophila Resource Center (VDRC) | FBst0472138 |
| UAS-Sec22[GD3175] | Vienna Drosophila Resource Center (VDRC) | FBst0471620 |
| UAS-Sec22[HMS01238] | Bloomington Drosophila Stock Center [BDSC] | FBst0034893 |
| UAS-Sec22[KK108519] | Vienna Drosophila Resource Center (VDRC) | FBst0472639 |
| UAS-Sec5[JF02676] | Bloomington Drosophila Stock Center [BDSC] | FBst0027526 |
| UAS-Snap24[GD4050] | Vienna Drosophila Resource Center (VDRC) | FBst0467689 |
| UAS-Snap24[JF03146] | Bloomington Drosophila Stock Center [BDSC] | FBst0028719 |
| UAS-Snap24[KK101630] | Vienna Drosophila Resource Center (VDRC) | FBst0480021 |
| UAS-Snap25[HMS01367] | Bloomington Drosophila Stock Center [BDSC] | FBst0034377 |
| UAS-Snap25[JF02615] | Bloomington Drosophila Stock Center [BDSC] | FBst0027306 |
| UAS-Snap29[GD7222] | Vienna Drosophila Resource Center (VDRC) | FBst0453044 |
| UAS-Snap29[HMC03467] | Bloomington Drosophila Stock Center [BDSC] | FBst0051893 |
| UAS-Snap29[JF01883] | Bloomington Drosophila Stock Center [BDSC] | FBst0025862 |

| | | |
|---|---|---|
| UAS-Snap29[KK108034] | Vienna Drosophila Resource Center (VDRC) | FBst0479760 |
| UAS-Syb[12210R-1] | National Institute of Genetics Fly Stocks (NIG-FLY) Mishima, Japan | FBst1086468 |
| UAS-Syb[GD4534] | Vienna Drosophila Resource Center (VDRC) | FBst0458569 |
| UAS-Syb[HMS01678] | Bloomington Drosophila Stock Center [BDSC] | FBst0038234 |
| UAS-Syb[HMS01987] | Bloomington Drosophila Stock Center [BDSC] | FBst0039067 |
| UAS-Syb[KK113351] | Vienna Drosophila Resource Center (VDRC) | FBst0474786 |
| UAS-Syx1[31136R-1] | National Institute of Genetics Fly Stocks (NIG-FLY) Mishima, Japan | - |
| UAS-Syx1[31136R-2 ] | National Institute of Genetics Fly Stocks (NIG-FLY) Mishima, Japan | - |
| UAS-Syx13.HA | Lu et al., 2013 (*3*) | FBtp0093144 |
| UAS-Syx13[GD2449] | Vienna Drosophila Resource Center (VDRC) | FBst0471035 |
| UAS-Syx13[HMS01723] | Bloomington Drosophila Stock Center [BDSC] | FBst0038525 |
| UAS-Syx13[JF01920] | Bloomington Drosophila Stock Center [BDSC] | FBst0027984 |
| UAS-Syx13[KK111650] | Vienna Drosophila Resource Center (VDRC) | FBst0474301 |
| UAS-Syx16[GD3694] | Vienna Drosophila Resource Center (VDRC) | FBst0471181 |
| UAS-Syx16[HMC03430] | Bloomington Drosophila Stock Center [BDSC] | FBst0051856 |
| UAS-Syx16[JF01924] | Bloomington Drosophila Stock Center [BDSC] | FBst0025884 |
| UAS-Syx16[KK108039] | Vienna Drosophila Resource Center (VDRC) | FBst0481191 |
| UAS-Syx18[13626R-2] | National Institute of Genetics Fly Stocks (NIG-FLY) Mishima, Japan | FBst1086931 |
| UAS-Syx18[GD241] | Vienna Drosophila Resource Center (VDRC) | FBst0450389 |
| UAS-Syx18[JF02263] | Bloomington Drosophila Stock Center [BDSC] | FBst0026721 |
| UAS-Syx18[KK101345] | Vienna Drosophila Resource Center (VDRC) | FBst0476941 |
| UAS-Syx4[GD8646] | Vienna Drosophila Resource Center (VDRC) | FBst0459577 |
| UAS-Syx4[HMS02771] | Bloomington Drosophila Stock Center [BDSC] | FBst0044054 |
| UAS-Syx4[JF01460] | Bloomington Drosophila Stock Center [BDSC] | FBst0031667 |
| UAS-Syx4[KK111728] | Vienna Drosophila Resource Center (VDRC) | FBst0474335 |

| | | |
|---|---|---|
| UAS-Syx5[GD1743] | Vienna Drosophila Resource Center (VDRC) | FBst0462587 |
| UAS-Syx5[JF03330] | Bloomington Drosophila Stock Center [BDSC] | FBst0029397 |
| UAS-Syx6[GD422] | Vienna Drosophila Resource Center (VDRC) | FBst0451699 |
| UAS-Syx6[JF03125] | Bloomington Drosophila Stock Center [BDSC] | FBst0028505 |
| UAS-Syx6[KK109340] | Vienna Drosophila Resource Center (VDRC) | FBst0476628 |
| UAS-Syx7[5081R-4] | National Institute of Genetics Fly Stocks (NIG-FLY) Mishima, Japan | - |
| UAS-Syx7[GD2767] | Vienna Drosophila Resource Center (VDRC) | FBst0470004 |
| UAS-Syx7[JF02436] | Bloomington Drosophila Stock Center [BDSC] | FBst0029546 |
| UAS-Syx7[KK101990] | Vienna Drosophila Resource Center (VDRC) | FBst0479086 |
| UAS-Syx8[GD2494] | Vienna Drosophila Resource Center (VDRC) | FBst0467104 |
| UAS-Syx8[JF02038] | Bloomington Drosophila Stock Center [BDSC] | FBst0026013 |
| UAS-Syx8[KK101612] | Vienna Drosophila Resource Center (VDRC) | FBst0478837 |
| UAS-Use1[GD2382] | Vienna Drosophila Resource Center (VDRC) | FBst0464643 |
| UAS-Use1[GLC01442] | Bloomington Drosophila Stock Center [BDSC] | FBst0043253 |
| UAS-Use1[KK102950] | Vienna Drosophila Resource Center (VDRC) | FBst0471893 |
| UAS-Vamp7[1599R-1] | National Institute of Genetics Fly Stocks (NIG-FLY) Mishima, Japan | FBal0272351 |
| UAS-Vamp7[GD4531] | Vienna Drosophila Resource Center (VDRC) | FBst0450940 |
| UAS-Vamp7[GL01524] | Bloomington Drosophila Stock Center [BDSC] | FBst0043543 |
| UAS-Vamp7[HMS01762] | Bloomington Drosophila Stock Center [BDSC] | FBst0038300 |
| UAS-Vamp7[KK107576] | Vienna Drosophila Resource Center (VDRC) | FBst0480543 |
| UAS-Vps11[KK102566] | Vienna Drosophila Resource Center (VDRC) | FBst0479241 |
| UAS-Vps41[18028R-2] | National Institute of Genetics Fly Stocks (NIG-FLY) Mishima, Japan | FBst1088273 |
| UAS-Vps8[KK100319] | Vienna Drosophila Resource Center (VDRC) | FBst0477778 |
| UAS-Vti1a[GD2233] | Vienna Drosophila Resource Center (VDRC) | FBst0466305 |
| UAS-Vti1a[HMS01727] | Bloomington Drosophila Stock Center [BDSC] | FBst0038526 |
| UAS-Ykt6.HA | Takáts et al, 2018 (*4*) | FBal0359940 |

| | | |
|---|---|---|
| UAS-Ykt6[1515R-1] | National Institute of Genetics Fly Stocks (NIG-FLY) Mishima, Japan | FBst1087524 |
| UAS-Ykt6[GD8927] | Vienna Drosophila Resource Center (VDRC) | FBst0453488 |
| UAS-Ykt6[HMJ21032] | Bloomington Drosophila Stock Center [BDSC] | FBst0050937 |
| UAS-Ykt6[HMS01778] | Bloomington Drosophila Stock Center [BDSC] | FBst0038314 |
| UAS-Ykt6[KK101343] | Vienna Drosophila Resource Center (VDRC) | FBst0477474 |
| Vps8[1] | Lőrincz et al., 2016 (*5*) | FBal0320420 |
| white[1118] | Bloomington Drosophila Stock Center [BDSC] | FBst0003605 |
| **Chemicals and other reagents** | | |
| Durcupan | Merck KGaA | 44614 |
| Tannic Acid | Mallinckrodt | 126420 |
| Formaldehyde solution | Merck KGaA | 252549 |
| FITC-Avidin | Thermo Fisher Scientific Inc. | 434411 |
| Lead citrate | Thermo Fisher Scientific Inc. | A10701.22 |
| anti-FLAG M2 affinity agarose gel beads | Millipore | A2220 |
| albumin from bovine serum (BSA) Alexa Fluor 647 | Thermo Fisher Scientific Inc. | A34785 |
| Sodium cacodylate trihydrate | Merck KGaA | C0250 |
| Calcium chloride | Merck KGaA | C1016 |
| 4′,6-Diamidino-2-phenylindole dihydrochloride (DAPI) | Merck KGaA | D8417 |
| Fetal Bovine Serum (FBS) | Merck KGaA | F4135 |
| Glutaraldehyde solution | Merck KGaA | G7776 |
| VECTASHIELD® Antifade Mounting Medium | Vector Laboratories Inc. | H-1000-10 |
| anti-HA rabbit agarose beads | Medical & Biological Laboratories CO., LTD. | MBL-561-8 |
| Osmium tetroxide | Merck KGaA | O5500 |
| 10 mM Phosphate Buffered Saline (PBS) ph 7.4 | Merck KGaA | P-3813 |
| Paraformaldehyde | Merck KGaA | P6148 |
| Sucrose | Merck KGaA | S0389 |

| | | |
|---|---|---|
| Shield and Sang M3 Insect Medium | Merck KGaA | S8398 |
| Uranyl Acetate EM Solution | TAAB | U001/S/2/10 |
| Immobilon Western Chemiluminescent HRP substrate | Millipore | WBKLS0500 |
| Triton X-100 | Merck KGaA | X100 |
| AgNO3 | Merck KGaA | 209139 |
| Dextran (10,000 MW) - AlexaFluor™ 568 | Eugene, Oregon | D22912 |
| HEPES | Merck KGaA | 83264 |
| $Na_2HPO_4$ | Merck KGaA | 567547 |
| NaCl | Merck KGaA | S5886 |
| $CuSO_4$ | Merck KGaA | 1.02780 |
| EDTA | Merck KGaA | E9884 |
| Tris hydrochloride | Merck KGaA | 10812846001 |
| cOmplete™, Mini, EDTA-free Protease Inhibitor Cocktail | Merck KGaA | 11836170001 |
| PMSF | Merck KGaA | PMSF-RO |
| Laemmli buffer | Merck KGaA | S3401 |
| 0.1% SDS | Sigma-Aldrich | 151-21-3 |
| Immun-Blot® PVDF Membrane | Bio-Rad Laboratories, Inc. | 1620174 |
| isopropylβ-D-1-thiogalactopyranoside (IPTG) | Merck KGaA | I6758 |
| Imidazole | Merck KGaA | I5513 |
| β-mercaptoethanol | Merck KGaA | 444203 |
| Lysozyme | Merck KGaA | 10837059001 |
| Ni Sepharose™ excel histidine-tagged protein purification resin | Cytiva, Chicago, IL, USA | 17371201 |
| Glycerol | Merck KGaA | G5516 |
| Amicon® Ultra Centrifugal Filter, 10 kDa MWCO | Merck KGaA | UFC8010 |
| Tris(2-carboxyethyl)phosphine hydrochloride (TCEP) | Merck KGaA | C4706 |
| Glutathione Sepharose™ High | Cytiva, Chicago, IL, USA | 17527901 |

| | | |
|---|---|---|
| Performance GST-tagged protein purification resin | | |
| Coomassie Brilliant Blue R | Merck KGaA | B7920 |
| **Software, Hardware and Services** | | |
| Adobe Photoshop CS4 | Adobe Inc. | SCR_014199 |
| Beckman JLA 9.1000 rotor | Beckman Coulter Life Sciences, Indianapolis, IN, USA | - |
| ChemiDoc™ Imaging System | BioRad | 12003153 |
| GraphPad Prism | GraphPad Software Inc. | SCR_002798 |
| ImageJ | ImageJ | SCR_003070 |
| iTEM | Olympus Soft Imaging Solutions GmbH, Münster, Germany | - |
| Sanger sequencing | Microsynth AG, Switzerland | - |
| Zeiss Efficient Navigation 2 software | Carl Zeiss AG | SCR_013672 |
| **Antibodies** | | |
| Alexa Fluor$^{TM}$ 568 donkey anti-mouse (IC: 1:1000) | Thermo Fisher Scientific Inc. | AB_11180865 |
| Dylight$^{TM}$ 488 goat anti-Guinea Pig (IC 1:600) | Thermo Fisher Scientific Inc. | SA5-10094 |
| Alexa Fluor$^{TM}$ 568 goat anti-rat (IC: 1:1000) | Thermo Fisher Scientific Inc. | AB_2534121 |
| Alexa Fluor$^{TM}$ 568 donkey anti-rabbit (IC: 1:1000) | Thermo Fisher Scientific Inc. | AB_2534017 |
| Dylight$^{TM}$ 488 goat anti-rabbit (IC: 1:1000) | Thermo Fisher Scientific Inc. | AB_2630356 |
| Dylight$^{TM}$ 488 goat anti-mouse (IC 1:600) | Thermo Fisher Scientific Inc. | AB_844397 |
| Alexa Fluor$^{TM}$ 488 goat anti-chicken (IC: 1:600) | Thermo Fisher Scientific Inc. | AB_2534096 |
| Alexa Fluor$^{TM}$ 488 donkey anti-rat (IC: 1:600) | Thermo Fisher Scientific Inc. | AB_2535794 |
| anti-Cubilin rat (IC: 1:200) | Atienza-Manuel, et al. 2021. (*6*) | - |

| | | |
|---|---|---|
| anti-FLAG mouse (WB: 1:2000) | Sigma | A9044 |
| anti-GFP Guinea-Pig (IC: 1:500) | Takats et al 2013. (*7*) | - |
| anti-HA rabbit (IC: 1:100) | Merck KGaA | AB_260070 |
| anti-HA rabbit (WB: 1:2500) | Proteintech | 51064-2-AP |
| anti-HA rat (IC: 1:100) | Roche Holding AG | AB_2314622 |
| anti-Lamp1 rabbit (IC: 1:1000) | Chaudhry et al. 2022. (*8*) | - |
| anti-mouse HRP (WB: 1:10000) | Sigma | AP308P |
| anti-Pyd mouse (IC: 1:400) | Developmental Studies Hybridoma Bank | AB_2618043 |
| anti-Rab5 rabbit (IC: 1:100) | Abcam Limited | AB_882240 |
| anti-Rab7 mouse (IC: 1:10) | Developmental Studies Hybridoma Bank | AB_2722471 |
| anti-rabbit HRP (WB: 1:10000) | Millipore | AP187P |
| anti-rat HRP (WB: 1:4000) | Sigma | A9037 |
| anti-Snap29 rat (IC: 1:300, WB: 1:2500) | Takats et al 2013. (*7*) | - |
| anti-Sns chicken (IC: 1:1000) | Hochapfel et al., 2017. (*9*) | - |
| Anti-Syx12L (Avl) | Lu and Bilder 2005. (*10*) | - |
| **cDNA, vectors and kits** | | |
| Syntaxin7 cDNA template | Drosophila Genomics Resource Center | DGRC_4835 |
| GeneJET Plasmid Miniprep Kit | Thermo Fischer Scientific Inc, Massachusetts, USA | K0502 |
| metallothionein-Gal4 (pMT-Gal4) plasmid | Drosophila Genomics Resource Center | DGRC_1042 |
| NEBuilder® HiFi DNA Assembly Cloning Kit | New England Biolabs Inc. Massachusetts, USA | E5520S |
| pETARA vector | Glatz et al. 2013. (*11*) | - |
| pETMBP vector | Glatz et al. 2013. (*11*) | - |
| pUAST-3xHA-V5-RpL10Ab | Addgene Inc. | 125223 |
| pUAST-attB-Flag-dTSC2 WT vector | Addgene Inc. | 111807 |
| **Cell lines and maintenance** | | |
| Ampicillin | Merck KGaA | A5354 |
| BL21 (DE3) E. coli cells | Merck KGaA | LGC604012 |

<table>
<tr><td>DH5-α competent E. coli cells</td><td colspan="2">New England Biolabs Inc. Massachusetts, USA</td><td>-</td></tr>
<tr><td>10% Fetal bovine serum (FBS)</td><td colspan="2">Euro Clone</td><td>ECS0183L</td></tr>
<tr><td>Insect-XPRESS™ Protein-free Insect Cell Medium</td><td colspan="2">Lonza</td><td>12-730Q</td></tr>
<tr><td>LB Broth (Miller) medium</td><td colspan="2">Merck KGaA</td><td>L2542</td></tr>
<tr><td>Penicillin-Streptomycin</td><td colspan="2">Thermo Fischer Scientific Inc, Massachusetts, USA</td><td>15140122</td></tr>
<tr><td>S2R+ Drosophila cell line - Stock 150</td><td colspan="2">Drosophila Genomics Resource Center</td><td>CVCL_Z831</td></tr>
<tr><td colspan="4">Primers used</td></tr>
<tr><td rowspan="2">Syx7 SNARE domain</td><td>1.</td><td>CGAGAATCTTTATTTTCAGGGATCCCTGCAGGCCCTCGAGG</td><td rowspan="12"></td></tr>
<tr><td>2.</td><td>GGTGGTGATGGTGGCGGCCGCTCGCCTTGCGCAGATTTTCC</td></tr>
<tr><td rowspan="2">Vamp7 SNARE</td><td>1.</td><td>CGAGAATCTTTATTTTCAGGGATCCACCATTTCGCGAGTACATGGTCAG</td></tr>
<tr><td>2.</td><td>TGGTGGTGGTGGTGCTCGAGCTTCCAAAACATTTGACGAGCCAAA</td></tr>
<tr><td rowspan="2">Ykt6 SNARE</td><td>1.</td><td>CGAGAATCTTTATTTTCAGGGATCCCCGCTGACGAAAATGCAAAAC</td></tr>
<tr><td>2.</td><td>TGGTGGTGGTGGTGCTCGAGGGTGAAGCTGCAGCAGGAG</td></tr>
<tr><td rowspan="2">SNAP29 Full length</td><td>1.</td><td>CGAGAATCTTTATTTTCAGGGATCCATGGCCCATAACTACCTGCA</td></tr>
<tr><td>2.</td><td>TGGTGGTGGTGGTGCTCGAGAAGCTTGCTCATGTCCTTGTTCT</td></tr>
<tr><td rowspan="2">Syx7 for FLAG-tag</td><td>1.</td><td>GAATTCAGATCTGCGGCCGCCATGGACTTACAGCATATGGAG</td></tr>
<tr><td>2.</td><td>TCCTCTAGAGGTACCCTCGACCTAATTCTTGAACTGAAAAACGAG</td></tr>
<tr><td rowspan="2">Syx7 for HA-tag</td><td>1.</td><td>CCAGACTACGCTGCCGGTACCATGGACTTACAGCATATGGAG</td></tr>
<tr><td>2.</td><td>CTTCACAAAGATCCTCTAGACTAATTCTTGAACTGAAAAACGAG</td></tr>
</table>

**Table S2. Genotypes of animals used in this study.**

| Figure | | Drosophila melanogaster genotype |
|---|---|---|
| Fig1. | B | white[1118]/(y); UAS-Dcr-2.D/+; UAS-Luciferase[JF01355]/Prospero-Gal4, UAS-GFP-Rab5 |
| | C | white[1118]/(y); UAS-Dcr-2.D/UAS-Syx7[KK101990]; +/Prospero-Gal4, UAS-GFP-Rab5 |
| | D | white[1118]/(y); UAS-Dcr-2.D/+; UAS-Snap29[JF01883]/Prospero-Gal4, UAS-GFP-Rab5 |
| | E | white[1118]/(y); UAS-Dcr-2.D/UAS-Ykt6[KK101343]; +/Prospero-Gal4, UAS-GFP-Rab5 |
| | H | white[1118]/(y); UAS-Dcr-2.D/+; UAS-Luciferase[JF01355]/Prospero-Gal4 |
| | I | white[1118]/(y); UAS-Dcr-2.D/UAS-Syx7[KK101990]; +/Prospero-Gal4 |
| | J | white[1118]/(y); UAS-Dcr-2.D/+; UAS-Snap29[JF01883]/Prospero-Gal4 |
| | K | white[1118]/(y); UAS-Dcr-2.D/UAS-Ykt6[KK101343]; +/Prospero-Gal4 |
| S1. | A | white[1118]/(y); UAS-Dcr-2.D/+;UAS-Luciferase[JF01355] /Prospero-Gal4 |
| | B1 | white[1118]/(y); UAS-Dcr-2.D/UAS-Syx1[31136R-1] ; +/Prospero-Gal4 |
| | B2 | white[1118]/(y); UAS-Dcr-2.D/UAS-Syx1[31136R-2 ] ; +/Prospero-Gal4 |
| | C1 | white[1118]/(y); UAS-Dcr-2.D/UAS-Syx4[KK111728]; +/Prospero-Gal4 |
| | C2 | white[1118]/(y); UAS-Dcr-2.D/+;UAS-Syx4[JF01460]/Prospero-Gal4 |
| | C3 | white[1118]/(y); UAS-Dcr-2.D/UAS-Syx4[GD8646]; +/Prospero-Gal4 |
| | C4 | white[1118]/(y); UAS-Dcr-2.D/UAS-Syx4[HMS02771]; +/Prospero-Gal4 |
| | D1 | white[1118]/(y); UAS-Dcr-2.D/+;UAS-Syx5[JF03330]/Prospero-Gal4 |
| | D2 | white[1118]/(y); UAS-Dcr-2.D/+;UAS-Syx5[GD1743]/Prospero-Gal4 |
| | E1 | white[1118]/(y); UAS-Dcr-2.D/UAS-Syx7[KK101990]; +/Prospero-Gal4 |
| | E2 | white[1118]/(y); UAS-Dcr-2.D/+;UAS-Syx7[JF02436]/Prospero-Gal4 |
| | E3 | white[1118]/(y); UAS-Dcr-2.D/+;UAS-Syx7[5081R-4]/Prospero-Gal4 |
| | E4 | white[1118]/(y); UAS-Dcr-2.D/+;UAS-Syx7[GD2767]/Prospero-Gal4 |
| | F1 | white[1118]/(y); UAS-Dcr-2.D/UAS-Syx13[KK111650]; +/Prospero-Gal4 |
| | F2 | white[1118]/(y); UAS-Dcr-2.D/UAS-Syx13[HMS01723]; +/Prospero-Gal4 |
| | F3 | white[1118]/(y); UAS-Dcr-2.D/+;UAS-Syx13[GD2449]/Prospero-Gal4 |
| | F4 | white[1118]/(y); UAS-Dcr-2.D/+;UAS-Syx13[JF01920]/Prospero-Gal4 |
| | G1 | white[1118]/(y); UAS-Dcr-2.D/UAS-Syx16[KK108039]; +/Prospero-Gal4 |
| | G2 | white[1118]/(y); UAS-Dcr-2.D/UAS-Syx16[HMC03430]; +/Prospero-Gal4 |
| | G3 | white[1118]/(y); UAS-Dcr-2.D/+;UAS-Syx16[JF01924]/Prospero-Gal4 |
| | G4 | white[1118]/(y); UAS-Dcr-2.D/+;UAS-Syx16[GD3694]/Prospero-Gal4 |

| | | |
|---|---|---|
| | H1 | white[1118]/(y); UAS-Dcr-2.D/+;UAS-Syx18[13626R-2]/Prospero-Gal4 |
| | H2 | white[1118]/(y); UAS-Dcr-2.D/UAS-Syx18[KK101345]; +/Prospero-Gal4 |
| | H3 | white[1118]/(y); UAS-Dcr-2.D/+;UAS-Syx18[JF02263]/Prospero-Gal4 |
| | H4 | white[1118]/(y); UAS-Dcr-2.D/+;UAS-Syx18[GD241]/Prospero-Gal4 |
| | I1 | white[1118]/(y); UAS-Dcr-2.D/+;UAS-Gos28[GD3051]/Prospero-Gal4 |
| | I2 | white[1118]/(y); UAS-Dcr-2.D/+;UAS-Gos28[HMS01203]/Prospero-Gal4 |
| | I3 | white[1118]/(y); UAS-Dcr-2.D/UAS-Gos28[KK107479]; +/Prospero-Gal4 |
| | J1 | white[1118]/(y); UAS-Dcr-2.D/UAS-Membrin[KK100283]; +/Prospero-Gal4 |
| | J2 | white[1118]/(y); UAS-Dcr-2.D/+;UAS-Membrin[4780R-3]/Prospero-Gal4 |
| | J3 | white[1118]/(y); UAS-Dcr-2.D/+;UAS-Membrin[GD2313]/Prospero-Gal4 |
| | J4 | white[1118]/(y); UAS-Dcr-2.D/+;UAS-Membrin[GLC01633]/Prospero-Gal4 |
| | K1 | white[1118]/(y); UAS-Dcr-2.D/UAS-Sec20[KK107334]; +/Prospero-Gal4 |
| | K2 | white[1118]/(y); UAS-Dcr-2.D/+;UAS-Sec20[HMS01172]/Prospero-Gal4 |
| | L1 | white[1118]/(y); UAS-Dcr-2.D/UAS-Vti1a[GD2233]; +/Prospero-Gal4 |
| | L2 | white[1118]/(y); UAS-Dcr-2.D/UAS-Vti1a[HMS01727]; +/Prospero-Gal4 |
| | M1 | white[1118]/(y); UAS-Dcr-2.D/UAS-Bet1[HMJ22351]; +/Prospero-Gal4 |
| | M2 | white[1118]/(y); UAS-Dcr-2.D/+;UAS-Bet1[GD2721]/Prospero-Gal4 |
| | N1 | white[1118]/(y); UAS-Dcr-2.D/UAS-Syx6[KK109340]; +/Prospero-Gal4 |
| | N2 | white[1118]/(y); UAS-Dcr-2.D/+;UAS-Syx6[JF03125]/Prospero-Gal4 |
| | N3 | white[1118]/(y); UAS-Dcr-2.D/UAS-Syx6[GD422]; +/Prospero-Gal4 |
| | O1 | white[1118]/(y); UAS-Dcr-2.D/UAS-Syx8[GD2494]; +/Prospero-Gal4 |
| | O2 | white[1118]/(y); UAS-Dcr-2.D/+;UAS-Syx8[JF02038]/Prospero-Gal4 |
| | O3 | white[1118]/(y); UAS-Dcr-2.D/UAS-Syx8[KK101612]; +/Prospero-Gal4 |
| | P1 | white[1118]/(y); UAS-Dcr-2.D/+;UAS-Use1[GD2382]/Prospero-Gal4 |
| | P2 | white[1118]/(y); UAS-Dcr-2.D/+;UAS-Use1[GLC01442]/Prospero-Gal4 |
| | P3 | white[1118]/(y); UAS-Dcr-2.D/UAS-Use1[KK102950]; +/Prospero-Gal4 |
| **S2.** | A1 | white[1118]/(y); UAS-Dcr-2.D/+;UAS-Snap24[JF03146]/Prospero-Gal4 |
| | A2 | white[1118]/(y); UAS-Dcr-2.D/+;UAS-Snap24[GD4050]/Prospero-Gal4 |
| | A3 | white[1118]/(y); UAS-Dcr-2.D/UAS-Snap24[KK101630]; +/Prospero-Gal4 |
| | B1 | white[1118]/(y); UAS-Dcr-2.D/+;UAS-Snap25[HMS01367]/Prospero-Gal4 |
| | B2 | white[1118]/(y); UAS-Dcr-2.D/+;UAS-Snap25[JF02615]/Prospero-Gal4 |
| | C1 | white[1118]/(y); UAS-Dcr-2.D/+;UAS-Snap29[JF01883]/Prospero-Gal4 |
| | C2 | white[1118]/(y); UAS-Dcr-2.D/UAS-Snap29[HMC03467]; +/Prospero-Gal4 |
| | C3 | white[1118]/(y); UAS-Dcr-2.D/UAS-Snap29[GD7222]; +/Prospero-Gal4 |

| | | |
|---|---|---|
| | C4 | white[1118]/(y); UAS-Dcr-2.D/UAS-Snap29[KK108034] ; +/Prospero-Gal4 |
| | D1 | white[1118]/(y); UAS-Dcr-2.D/+;UAS-Sec22[HMS01238]/Prospero-Gal4 |
| | D2 | white[1118]/(y); UAS-Dcr-2.D/UAS-Sec22[KK108519]; +/Prospero-Gal4 |
| | D3 | white[1118]/(y); UAS-Dcr-2.D/UAS-Sec22[GD3175]; +/Prospero-Gal4 |
| | E1 | white[1118]/(y); UAS-Dcr-2.D/+;UAS-Syb[12210R-1]/Prospero-Gal4 |
| | E2 | white[1118]/(y); UAS-Dcr-2.D/+;UAS-Syb[HMS01678]/Prospero-Gal4 |
| | E3 | white[1118]/(y); UAS-Dcr-2.D/+;UAS-Syb[GD4534]/Prospero-Gal4 |
| | E4 | white[1118]/(y); UAS-Dcr-2.D/UAS-Syb[KK113351]; +/Prospero-Gal4 |
| | E5 | white[1118]/(y); UAS-Dcr-2.D/UAS-Syb[HMS01987]; +/Prospero-Gal4 |
| | F1 | white[1118]/(y); UAS-Dcr-2.D/UAS-nSyb[KK109893]; +/Prospero-Gal4 |
| | F2 | white[1118]/(y); UAS-Dcr-2.D/+;UAS-nSyb[GD17382]/Prospero-Gal4 |
| | F3 | white[1118]/(y); UAS-Dcr-2.D/UAS-nSyb[GD17382]; +/Prospero-Gal4 |
| | F4 | white[1118]/(y); UAS-Dcr-2.D/+;UAS-nSyb[JF03417]/Prospero-Gal4 |
| | G1 | white[1118]/(y); UAS-Dcr-2.D/UAS-Vamp7[KK107576]; +/Prospero-Gal4 |
| | G2 | white[1118]/(y); UAS-Dcr-2.D/UAS-Vamp7[HMS01762]; +/Prospero-Gal4 |
| | G3 | white[1118]/(y); UAS-Dcr-2.D/+;UAS-Vamp7[GL01524]/Prospero-Gal4 |
| | G4 | white[1118]/(y); UAS-Dcr-2.D/+;UAS-Vamp7[GD4531]/Prospero-Gal4 |
| | G5 | white[1118]/(y); UAS-Dcr-2.D/UAS-Vamp7[1599R-1]; +/Prospero-Gal4 |
| | H1 | white[1118]/(y); UAS-Dcr-2.D/UAS-Ykt6[HMJ21032]; +/Prospero-Gal4 |
| | H2 | white[1118]/(y); UAS-Dcr-2.D/UAS-Ykt6[GD8927]; +/Prospero-Gal4 |
| | H3 | white[1118]/(y); UAS-Dcr-2.D/UAS-Ykt6[HMS01778]; +/Prospero-Gal4 |
| | H4 | white[1118]/(y); UAS-Dcr-2.D/UAS-Ykt6[1515R-1]; +/Prospero-Gal4 |
| | H5 | white[1118]/(y); UAS-Dcr-2.D/UAS-Ykt6[KK101343]; +/Prospero-Gal4 |
| **Fig2.** | D, L, Q | white[1118]/(y); UAS-Dcr-2.D/+;UAS-Luciferase[JF01355] /Prospero-Gal4 |
| | F, M, R | white[1118]/(y); UAS-Dcr-2.D/UAS-Syx7[KK101990]; +/Prospero-Gal4 |
| | H, N, S | white[1118]/(y); UAS-Dcr-2.D/+;UAS-Snap29[JF01883]/Prospero-Gal4 |
| | I, O, T | white[1118]/(y); UAS-Dcr-2.D/UAS-Ykt6[KK101343]; +/Prospero-Gal4 |
| **S3.** | A | white[1118]/(y); UAS-Dcr-2.D/+;UAS-Luciferase[JF01355] /Prospero-Gal4 |
| | B | white[1118]/(y); UAS-Dcr-2.D/UAS-Syx1[31136R-1] ; +/Prospero-Gal4 |
| | C | white[1118]/(y); UAS-Dcr-2.D/UAS-Syx4[GD8646]; +/Prospero-Gal4 |
| | D | white[1118]/(y); UAS-Dcr-2.D/+;UAS-Syx5[GD1743]/Prospero-Gal4 |
| | E | white[1118]/(y); UAS-Dcr-2.D/UAS-Syx7[KK101990]; +/Prospero-Gal4 |
| | F | white[1118]/(y); UAS-Dcr-2.D/+;UAS-Syx13[JF01920]/Prospero-Gal4 |

| | | |
|---|---|---|
| | G | white[1118]/(y); UAS-Dcr-2.D/UAS-Syx16[KK108039]; +/Prospero-Gal4 |
| | H | white[1118]/(y); UAS-Dcr-2.D/+;UAS-Syx18[JF02263]/Prospero-Gal4 |
| | I | white[1118]/(y); UAS-Dcr-2.D/UAS-Gos28[KK107479]; +/Prospero-Gal4 |
| | J | white[1118]/(y); UAS-Dcr-2.D/+;UAS-Membrin[4780R-3]/Prospero-Gal4 |
| | K | white[1118]/(y); UAS-Dcr-2.D/UAS-Sec20[KK107334]; +/Prospero-Gal4 |
| | L | white[1118]/(y); UAS-Dcr-2.D/UAS-Vti1a[HMS01727]; +/Prospero-Gal4 |
| | M | white[1118]/(y); UAS-Dcr-2.D/UAS-Bet1[HMJ22351]; +/Prospero-Gal4 |
| | N | white[1118]/(y); UAS-Dcr-2.D/UAS-Syx6[KK109340]; +/Prospero-Gal4 |
| | O | white[1118]/(y); UAS-Dcr-2.D/UAS-Syx8[GD2494]; +/Prospero-Gal4 |
| | P | white[1118]/(y); UAS-Dcr-2.D/UAS-Use1[KK102950]; +/Prospero-Gal4 |
| | Q | white[1118]/(y); UAS-Dcr-2.D/+;UAS-Snap24[JF03146]/Prospero-Gal4 |
| | R | white[1118]/(y); UAS-Dcr-2.D/+;UAS-Snap25[JF02615]/Prospero-Gal4 |
| | S | white[1118]/(y); UAS-Dcr-2.D/+;UAS-Snap29[JF01883]/Prospero-Gal4 |
| | T | white[1118]/(y); UAS-Dcr-2.D/UAS-Sec22[KK108519]; +/Prospero-Gal4 |
| | U | white[1118]/(y); UAS-Dcr-2.D/+;UAS-Syb[12210R-1]/Prospero-Gal4 |
| | V | white[1118]/(y); UAS-Dcr-2.D/+;UAS-nSyb[JF03417]/Prospero-Gal4 |
| | W | white[1118]/(y); UAS-Dcr-2.D/+;UAS-Vamp7[GL01524]/Prospero-Gal4 |
| | X | white[1118]/(y); UAS-Dcr-2.D/UAS-Ykt6[KK101343]; +/Prospero-Gal4 |
| | Y1 | white[1118]/(y); UAS-Dcr-2.D/+;UAS-Luciferase[JF01355] /Prospero-Gal4 |
| | Y2 | white[1118]/(y); UAS-Dcr-2.D/+;UAS-Snap29[JF01883]/Prospero-Gal4 |
| | Y3 | white[1118]/(y); UAS-Dcr-2.D/UAS-Syx7[KK101990]; +/Prospero-Gal4 |
| | Y4 | white[1118]/(y); UAS-Dcr-2.D/UAS-Ykt6[KK101343]; +/Prospero-Gal4 |
| | Z | white[1118]/(y); UAS-Dcr-2.D/+;UAS-Snap29[JF01883]/Prospero-Gal4 |
| **Fig3.** | A, B | white[1118]/(y); UAS-Dcr-2.D/+; +/Prospero-Gal4, UAS-GFP-Rab5 |
| | C | white[1118]/UAS-Ykt6.HA; UAS-Dcr-2.D/+; +/Prospero-Gal4, UAS-GFP-Rab5 |
| | D, E, H | white[1118]/(y); +/+; +/+ |
| | F, I | white[1118]/UAS-Ykt6.HA; UAS-Dcr-2.D/+; +/Prospero-Gal4 |
| **Fig4.** | B | white[1118]/(y); UAS-Dcr-2.D/UAS-Syx13.HA; +/Prospero-Gal4, UAS-GFP-Rab5 |
| | C, D | white[1118]/(y); UAS-Dcr-2.D/UAS-Syx13.HA; +/Prospero-Gal4 |
| | E | white[1118]/(y); UAS-Dcr-2.D/+;UAS-Luciferase[JF01355] /Prospero-Gal4 |
| | F | white[1118]/(y); UAS-Dcr-2.D/+; UAS-hsp-Hsap\STX7.HA.1/Prospero-Gal4 |
| | G | white[1118]/(y); UAS-Dcr-2.D/+; UAS-hsp-Hsap\STX12.HA.1/Prospero-Gal4 |
| | H | white[1118]/(y); UAS-Dcr-2.D/UAS-Syx7[KK101990];+/Prospero-Gal4 |

| | | |
|---|---|---|
| | I | white[1118]/(y); UAS-Dcr-2.D/UAS-Syx7[KK101990]; UAS-hsp-Hsap\STX7.HA.1/Prospero-Gal4 |
| | J | white[1118]/(y); UAS-Dcr-2.D/UAS-Syx7[KK101990]; UAS-hsp-Hsap\STX12.HA.1/Prospero-Gal4 |
| | K | white[1118]/(y); UAS-Dcr-2.D/UAS-Syx13[KK111650];+/Prospero-Gal4 |
| | L | white[1118]/(y); UAS-Dcr-2.D/UAS-Syx13[KK111650]; UAS-hsp-Hsap\STX7.HA.1/Prospero-Gal4 |
| | M | white[1118]/(y); UAS-Dcr-2.D/UAS-Syx13[KK111650]; UAS-hsp-Hsap\STX12.HA.1/Prospero-Gal4 |
| | O | white[1118]/(y); UAS-Dcr-2.D/+;UAS-Luciferase[JF01355] /Prospero-Gal4 |
| | P | white[1118]/(y); UAS-Dcr-2.D/+; UAS-hsp-Hsap\STX7.HA.1/Prospero-Gal4 |
| | Q | white[1118]/(y); UAS-Dcr-2.D/+; UAS-hsp-Hsap\STX12.HA.1/Prospero-Gal4 |
| | R | white[1118]/(y); UAS-Dcr-2.D/UAS-Syx7[KK101990];+/Prospero-Gal4 |
| | S | white[1118]/(y); UAS-Dcr-2.D/UAS-Syx7[KK101990]; UAS-hsp-Hsap\STX7.HA.1/Prospero-Gal4 |
| | T | white[1118]/(y); UAS-Dcr-2.D/UAS-Syx7[KK101990]; UAS-hsp-Hsap\STX12.HA.1/Prospero-Gal4 |
| **Fig5.** | A, F | white[1118]/(y); UAS-Dcr-2.D/+;UAS-Luciferase[JF01355] /Prospero-Gal4 |
| | B, G | white[1118]/(y); UAS-Dcr-2.D/UAS-Syx7[KK101990]; +/Prospero-Gal4<br>Note: Syx7 was renamed to Syx12L in this paper |
| | C, H | white[1118]/(y); UAS-Dcr-2.D/+;UAS-Snap29[JF01883]/Prospero-Gal4 |
| | D, I | white[1118]/(y); UAS-Dcr-2.D/UAS-Ykt6[KK101343]; +/Prospero-Gal4 |
| | J | white[1118]/(y); UAS-Dcr-2.D/+; UAS-Sec5[JF02676]/Prospero-Gal4 |
| | K | white[1118]/(y); UAS-Dcr-2.D/+;UAS-Sec5[JF02676]/UAS-Snap29[JF01883],Prospero-Gal4 |
| **S4.** | A1 | white[1118]/(y); UAS-Dcr-2.D/UAS-Rbsn5[HMC04769]; +/Prospero-Gal4 |
| | A2 | white[1118]/(y); UAS-Dcr-2.D/UAS-Vps8[KK100319] ; +/Prospero-Gal4 |
| | A3 | white[1118]/(y); UAS-Dcr-2.D/UAS-Vps11[KK102566] ; +/Prospero-Gal4 |
| | A4 | white[1118]/(y); UAS-Dcr-2.D/UAS-Vps41[18028R-2]; +/Prospero-Gal4 |
| | A5 | white[1118]/(y); UAS-Dcr-2.D/UAS-Syx13[KK111650]; +/Prospero-Gal4<br>Note: Syx13 was renamed to Syx7L in this paper |
| | A6 | white[1118]/(y); UAS-Dcr-2.D/UAS-Ykt6[KK101343];UAS-Snap29[JF01883]/Prospero-Gal4 |
| | B, C, L, M, U | white[1118]/(y); UAS-Dcr-2.D/+;UAS-Luciferase[JF01355] /Prospero-Gal4 |
| | D, E, N, O, V | white[1118]/(y); UAS-Dcr-2.D/UAS-Syx7[KK101990]; +/Prospero-Gal4<br>Note: Syx7 was renamed to Syx12L in this paper |
| | F, G, P, Q, W | white[1118]/(y); UAS-Dcr-2.D/+;UAS-Snap29[JF01883]/Prospero-Gal4 |
| | H, I, R, S, X | white[1118]/(y); UAS-Dcr-2.D/UAS-Ykt6[KK101343]; +/Prospero-Gal4 |

| | | |
|---|---|---|
| Fig6. | A | white[1118]/(y); UAS-Dcr-2.D/+;UAS-Luciferase[JF01355] /Prospero-Gal4 |
| | B | white[1118]/(y); UAS-Dcr-2.D/UAS-Syx7[KK101990]; +/Prospero-Gal4 Note: Syx7 was renamed to Syx12L in this paper |
| | C | white[1118]/(y); UAS-Dcr-2.D/+;UAS-Snap29[JF01883]/Prospero-Gal4 |
| | D | white[1118]/(y); UAS-Dcr-2.D/UAS-Ykt6[KK101343]; +/Prospero-Gal4 |
| | E | white[1118]/(y); UAS-Dcr-2.D/UAS-Rab5[GL01872]; +/Prospero-Gal4 |
| | F | white[1118]/(y); UAS-Dcr-2.D/UAS-Rbsn5[HMC04769]; +/Prospero-Gal4 |
| | G | white[1118]/(y); UAS-Dcr-2.D/UAS-Vps8[KK100319] ; +/Prospero-Gal4 |
| | H | white[1118]/(y); UAS-pHluorinSE-Rab2/+;UAS-Luciferase[JF01355] /Prospero-Gal4 |
| | I | white[1118]/(y); UAS-pHluorinSE-Rab2/UAS-Syx7[KK101990]; +/Prospero-Gal4 Note: Syx7 was renamed to Syx12L in this paper |
| | J | white[1118]/(y); UAS-pHluorinSE-Rab2/+;UAS-Snap29[JF01883]/Prospero-Gal4 |
| | K | white[1118]/(y); UAS-pHluorinSE-Rab2/UAS-Ykt6[KK101343]; +/Prospero-Gal4 |
| S5. | A | white[1118]/(y); UAS-Dcr-2.D/UAS-Syx7[KK101990]; +/Prospero-Gal4 Note: Syx7 was renamed to Syx12L in this paper |
| | B | white[1118]/(y); UAS-Dcr-2.D/UAS-Rbsn5[HMC04769]; +/Prospero-Gal4 |
| | C, E | white[1118]/(y); +/+; Vps8[1]/Df(3L)ED211 |
| | D | white[1118]/(y); +/+; +/+ |
| | F | white[1118]/(y); UAS-GFP-myc-2xFYVE/UAS-Syx7[KK101990]; +/Prospero-Gal4 Note: Syx7 was renamed to Syx12L in this paper |
| | G | white[1118]/(y); UAS-EGFP-Clc/UAS-Syx7[KK101990]; +/Prospero-Gal4 Note: Syx7 was renamed to Syx12L in this paper |
| Fig7. | A | white[1118]/(y); UAS-Dcr-2.D/UAS-Syx7[KK101990]; +/Prospero-Gal4, UAS-GFP-Rab5 Note: Syx7 was renamed to Syx12L in this paper |
| | B | white[1118]/(y); UAS-Dcr-2.D/UAS-Ykt6[KK101343]; +/Prospero-Gal4, UAS-GFP-Rab5 |
| | C | white[1118]/(y); UAS-Dcr-2.D/+; UAS-Snap29[JF01883]/Prospero-Gal4, UAS-GFP-Rab5 |
| | D | white[1118]/(y); UAS-Dcr-2.D/UAS-Ykt6[KK101343]; UAS-Syx7[JF02436]/Prospero-Gal4, UAS-GFP-Rab5 Note: Syx7 was renamed to Syx12L in this paper |
| | E | white[1118]/(y); UAS-Dcr-2.D/UAS-Syx7[KK101990]; UAS-Snap29[JF01883]/Prospero-Gal4, UAS-GFP-Rab5 Note: Syx7 was renamed to Syx12L in this paper |
| | F | white[1118]/(y); UAS-Dcr-2.D/UAS-Ykt6[KK101343]; UAS-Snap29[JF01883]/Prospero-Gal4, UAS-GFP-Rab5 |
| | I | white[1118]/(y); UAS-Dcr-2.D/+; UAS-Vamp7[GL01524]/Prospero-Gal4, UAS-GFP-Rab5 |
| | J | white[1118]/(y); UAS-Dcr-2.D/UAS-Ykt6[KK101343]; UAS-Vamp7[GL01524]/Prospero-Gal4, UAS-GFP-Rab5 |
| | L, M, N, O | white[1118]/(y); UAS-Dcr-2.D/UAS-GFP-VAMP7; +/Prospero-Gal4 |
| | P | white[1118]/(y); UAS-Dcr-2.D/+; UAS-Syx13[JF01920]/Prospero-Gal4, UAS-GFP-Rab5 Note: Syx13 was renamed to Syx7L in this paper |

| | | |
|---|---|---|
| | Q | white[1118]/(y); UAS-Dcr-2.D/UAS-Syx7[KK101990]; UAS-Syx13[JF01920]/Prospero-Gal4, UAS-GFP-Rab5 Note: Syx7 and Syx13 were renamed to Syx12L and Syx7L in this paper, respectively. |
| | R | white[1118]/(y); UAS-Dcr-2.D/UAS-Syx7[KK101990]; UAS-Vamp7[GL01524]/Prospero-Gal4, UAS-GFP-Rab5 Note: Syx7 was renamed to Syx12L in this paper |
| | S | white[1118]/(y); UAS-Dcr-2.D/UAS-Ykt6[KK101343]; UAS-Syx13[JF01920]/Prospero-Gal4, UAS-GFP-Rab5 Note: Syx13 was renamed to Syx7L in this paper |
| | T | white[1118]/(y); UAS-Dcr-2.D/UAS-Vamp7[1599R-1]; UAS-Syx13[JF01920]/Prospero-Gal4, UAS-GFP-Rab5 Note: Syx13 was renamed to Syx7L in this paper |
| | U | white[1118]/(y); UAS-Syx13.HA/UAS-GFP-VAMP7; +/Prospero-Gal4 Note: Syx13 was renamed to Syx7L in this paper |
| **Fig8.** | A | white[1118]/(y); +/UASp-YFP.Rab5; UAS-Luciferase[JF01355] /Prospero-Gal4 |
| | B | white[1118]/(y); +/UASp-YFP.Rab5; UAS-Snap29[JF01883]/Prospero-Gal4 |
| | C | white[1118]/(y); UAS-Syx7[KK101990]/UASp-YFP.Rab5; +/Prospero-Gal4 Note: Syx7 was renamed to Syx12L in this paper |
| | D | white[1118]/(y); UAS-Ykt6[KK101343]/UASp-YFP.Rab5; +/Prospero-Gal4 |
| | F | white[1118]/(y); +/UASp-YFP.Rab5.Q88L; UAS-Luciferase[JF01355] /Prospero-Gal4 |
| | G | white[1118]/(y); +/UASp-YFP.Rab5.Q88L; UAS-Snap29[JF01883]/Prospero-Gal4 |
| | H | white[1118]/(y); UAS-Syx7[KK101990]/UASp-YFP.Rab5.Q88L; +/Prospero-Gal4 Note: Syx7 was renamed to Syx12L in this paper |
| | I | white[1118]/(y); UAS-Ykt6[KK101343]/UASp-YFP.Rab5.Q88L; +/Prospero-Gal4 |
| | K | white[1118]/(y); +/UASp-YFP.Rab5; UAS-Luciferase[JF01355] /Prospero-Gal4, UAS-Rab7[JF02377] |
| | L | white[1118]/(y); +/UASp-YFP.Rab5; UAS-Snap29[JF01883]/Prospero-Gal4, UAS-Rab7[JF02377] |
| | M | white[1118]/(y); UAS-Syx7[KK101990]/UASp-YFP.Rab5; +/Prospero-Gal4, UAS-Rab7[JF02377] Note: Syx7 was renamed to Syx12L in this paper |
| | N | white[1118]/(y); UAS-Ykt6[KK101343]/UASp-YFP.Rab5; +/Prospero-Gal4, UAS-Rab7[JF02377] |
| | P | white[1118]/(y); +/UASp-YFP.Rab5.Q88L; UAS-Luciferase[JF01355] /Prospero-Gal4, UAS-Rab7[JF02377] |
| | Q | white[1118]/(y); +/UASp-YFP.Rab5.Q88L; UAS-Snap29[JF01883]/Prospero-Gal4, UAS-Rab7[JF02377] |
| | R | white[1118]/(y); UAS-Syx7[KK101990]/UASp-YFP.Rab5.Q88L; +/Prospero-Gal4, UAS-Rab7[JF02377] Note: Syx7 was renamed to Syx12L in this paper |
| | S | white[1118]/(y); UAS-Ykt6[KK101343]/UASp-YFP.Rab5.Q88L; +/Prospero-Gal4, UAS-Rab7[JF02377] |

**Table S3. Statistical details.**

| Figure 1 | | | | | |
|---|---|---|---|---|---|
| **Panel A: Inset from (Supplementary Figure 2 Panel I)** | | | | | |
| **Panel F** | | | | | |
| **Measured:** | Diameter of Rab5-GFP+ endosomes (micrometer) | | | | |
| **Statistical test:** | Kruskal-Wallis test with Dunn's multiple comparisons test | | | | |
| **Simplified genotype** | **N** | **Mean** | **Standard deviation** | **P-value** | |
| | | | | **Compared to Luc Ri** | **Compared to Snap29 Ri** |
| Luc Ri | 100 | 1,041 | 0,3794 | - | <0,0001 |
| Syx7 Ri | 100 | 0,3446 | 0,1163 | <0,0001 | 0,0111 |
| Snap29 Ri | 100 | 0,2627 | 0,08403 | <0,0001 | - |
| Ykt6 Ri | 100 | 0,5871 | 0,2065 | <0,0001 | <0,0001 |
| **Panel G** | | | | | |
| **Measured:** | Threshold Overlap Score (TOS) between Rab5-GFP and a-Rab7 per cell | | | | |
| **Statistical test:** | Ordinary one-way ANOVA with Tukey's multiple comparisons test | | | | |
| **Simplified genotype** | **N** | **Mean** | **Standard deviation** | **P-value** | |
| | | | | **Compared to Luc Ri** | |
| Luc Ri | 12 | -0,0056 | 0,04477 | - | |
| Syx7 Ri | 12 | 0,5105 | 0,09546 | <0,0001 | |
| Snap29 Ri | 12 | 0,5924 | 0,1585 | <0,0001 | |
| Ykt6 Ri | 12 | 0,6349 | 0,1211 | <0,0001 | |
| **Panel L** | | | | | |
| **Measured:** | Diameter of a-Lamp1+ lysosomes (micrometer) | | | | |
| **Statistical test:** | Kruskal-Wallis test with Dunn's multiple comparisons test | | | | |
| **Simplified genotype** | **N** | **Mean** | **Standard deviation** | **P-value** | |
| | | | | **Compared to Luc Ri** | **Compared to Snap29 Ri** |
| Luc Ri | 100 | 0,9992 | 0,3486 | - | <0,0001 |
| Syx7 Ri | 100 | 0,602 | 0,732 | <0,0001 | <0,0001 |
| Snap29 Ri | 100 | 0,2452 | 0,07815 | <0,0001 | - |
| Ykt6 Ri | 100 | 0,4468 | 0,1408 | <0,0001 | <0,0001 |
| **Panel M** | | | | | |
| **Measured:** | Threshold Overlap Score (TOS) between a-Lamp1 and a-Rab7 per cell | | | | |
| **Statistical test:** | Ordinary one-way ANOVA with Tukey's multiple comparisons test | | | | |
| **Simplified genotype** | **N** | **Mean** | **Standard deviation** | **P-value** | |
| | | | | **Compared to Luc Ri** | |
| Luc Ri | 12 | 0,05277 | 0,0595 | - | |
| Syx7 Ri | 12 | 0,6269 | 0,1961 | <0,0001 | |

| | | | | |
|---|---|---|---|---|
| Snap29 Ri | 12 | 0,0227 | 0,0977 | 0,9434 |
| Ykt6 Ri | 12 | 0,4691 | 0,1334 | <0,0001 |

### Figure 2

#### Panel A

| | |
|---|---|
| **Measured:** | FITC-Avidin tracer coverage of the cell (%) |
| **Statistical test:** | Ordinary one-way ANOVA with Dunnett's multiple comparisons test |

| **Simplified genotype** | **N** | **Mean** | **Standard deviation** | **P-value** |
|---|---|---|---|---|
| | | | | **Compared to Luc Ri** |
| Luc Ri | 15 | 5,101 | 1,061 | - |
| Syx1 Ri | 15 | 3,773 | 0,9496 | 0,003 |
| Syx4 RI | 15 | 5,314 | 0,9921 | 0,9991 |
| Syx5 Ri | 15 | 3,753 | 0,9149 | 0,0024 |
| Syx7 Ri | 15 | 0 | 0 | <0,0001 |
| Syx13 Ri | 15 | 5,017 | 0,9194 | 0,9997 |
| Syx16 Ri | 15 | 4,985 | 1,035 | 0,9996 |
| Syx18 Ri | 15 | 5,168 | 0,9509 | 0,9997 |
| Gos28 Ri | 15 | 5,192 | 1,195 | 0,9997 |
| Membrin Ri | 15 | 5,065 | 1,213 | 0,9999 |
| Sec20 Ri | 15 | 5,176 | 1,007 | 0,9997 |
| Vti1a Ri | 15 | 4,956 | 0,9622 | 0,9994 |
| Bet1 Ri | 15 | 4,893 | 0,9527 | 0,9991 |
| Syx6 Ri | 15 | 5,153 | 1,169 | 0,9998 |
| Syx8 Ri | 15 | 5,076 | 1,139 | >0,9999 |
| Use1 Ri | 13 | 3,49 | 0,6768 | 0,0002 |
| Snap24 Ri | 15 | 5,362 | 0,9007 | 0,9988 |
| Snap25 Ri | 15 | 5,081 | 1,038 | >0,9999 |
| Snap29 Ri | 15 | 0 | 0 | <0,0001 |
| Sec22 Ri | 15 | 5,492 | 0,8778 | 0,9821 |
| Syb Ri | 15 | 5,257 | 0,9598 | 0,9994 |
| nSyb Ri | 15 | 5,315 | 1,038 | 0,9991 |
| Vamp7 Ri | 15 | 4,811 | 1,032 | 0,9952 |
| Ykt6 Ri | 15 | 0 | 0 | <0,0001 |

#### Panel C

| | |
|---|---|
| **Measured:** | FITC-Avidin tracer coverage of the cell (%) - Luc Ri |
| **Statistical test:** | Ordinary one-way ANOVA with Tukey's multiple comparisons test |

| **Pulse/chase** | **N** | **Mean** | **Standard deviation** | **P-value** | |
|---|---|---|---|---|---|
| | | | | **Compared to 5/0** | **Compared to 30/0** |
| 5/0 | 15 | 5,101 | 1,061 | - | <0,0001 |
| 30/0 | 15 | 11,15 | 2,931 | <0,0001 | - |
| 30/10 | 15 | 11,14 | 2,548 | <0,0001 | 0,9998 |

| | |
|---|---|
| **Measured:** | Dextran tracer coverage of the cell (%) - Luc Ri |
| **Statistical test:** | Ordinary one-way ANOVA with Tukey's multiple comparisons test |

| **Pulse/chase** | **N** | **Mean** | | **P-value** |
|---|---|---|---|---|

| | | | **Standard deviation** | **Compared to 5/0** | **Compared to 30/0** |
|---|---|---|---|---|---|
| 5/0 | 15 | 5,58 | 1,032 | - | <0,0001 |
| 30/0 | 15 | 11,57 | 2,906 | <0,0001 | - |
| 30/10 | 15 | 11,24 | 2,623 | <0,0001 | 0,9222 |
| **Panel E** | | | | | |
| **Measured:** | FITC-Avidin tracer coverage of the cell (%) - Syx7 Ri | | | | |
| **Statistical test:** | - | | | | |
| **Pulse/chase** | **N** | **Mean** | **Standard deviation** | **P-value** | |
| | | | | **Compared to 5/0** | **Compared to 30/0** |
| 5/0 | 15 | 0 | 0 | n.s. | |
| 30/0 | 15 | 0 | 0 | | |
| 30/10 | 15 | 0 | 0 | | |
| **Measured:** | Dextran tracer coverage of the cell (%) - Syx7 Ri | | | | |
| **Statistical test:** | Kruskal-Wallis test with Dunn's multiple comparisons test | | | | |
| **Pulse/chase** | **N** | **Mean** | **Standard deviation** | **P-value** | |
| | | | | **Compared to 5/0** | **Compared to 30/0** |
| 5/0 | 15 | 3,094 | 0,8097 | - | >0,9999 |
| 30/0 | 15 | 3,559 | 1,172 | >0,9999 | - |
| 30/10 | 15 | 0,1201 | 0,1838 | <0,0001 | <0,0001 |
| **Panel G** | | | | | |
| **Measured:** | FITC-Avidin tracer coverage of the cell (%) - Snap29 Ri | | | | |
| **Statistical test:** | - | | | | |
| **Pulse/chase** | **N** | **Mean** | **Standard deviation** | **P-value** | |
| | | | | **Compared to 5/0** | **Compared to 30/0** |
| 5/0 | 15 | 0 | 0 | n.s. | |
| 30/0 | 15 | 0 | 0 | | |
| 30/10 | 15 | 0 | 0 | | |
| **Measured:** | Dextran tracer coverage of the cell (%) - Snap29 Ri | | | | |
| **Statistical test:** | Ordinary one-way ANOVA with Tukey's multiple comparisons test | | | | |
| **Pulse/chase** | **N** | **Mean** | **Standard deviation** | **P-value** | |
| | | | | **Compared to 5/0** | **Compared to 30/0** |
| 5/0 | 15 | 3,293 | 0,9987 | - | 0,4928 |
| 30/0 | 15 | 3,674 | 1,189 | 0,4928 | - |
| 30/10 | 15 | 0,2081 | 0,2944 | <0,0001 | <0,0001 |
| **Panel I** | | | | | |
| **Measured:** | FITC-Avidin tracer coverage of the cell (%) - Ykt6 Ri | | | | |
| **Statistical test:** | Ordinary one-way ANOVA with Tukey's multiple comparisons test | | | | |
| **Pulse/chase** | **N** | **Mean** | **Standard deviation** | **P-value** | |
| | | | | **Compared to 5/0** | **Compared to 30/0** |
| 5/0 | 15 | 0 | 0 | - | <0,0001 |
| 30/0 | 15 | 3,293 | 1,313 | <0,0001 | - |

| | | | | | |
|---|---|---|---|---|---|
| 30/10 | 15 | 3,566 | 1,363 | <0,0001 | 0,7746 |
| **Measured:** | Dextran tracer coverage of the cell (%) - Ykt6 Ri | | | | |
| **Statistical test:** | Ordinary one-way ANOVA with Tukey's multiple comparisons test | | | | |
| **Pulse/chase** | **N** | **Mean** | **Standard deviation** | **P-value** | |
| | | | | **Compared to 5/0** | **Compared to 30/0** |
| 5/0 | 15 | 2,706 | 0,9545 | - | 0,0223 |
| 30/0 | 15 | 3,982 | 1,456 | 0,0223 | - |
| 30/10 | 15 | 3,875 | 1,323 | 0,039 | 0,9709 |
| **Panel K** | | | | | |
| **Measured:** | FITC-Avidin tracer coverage of the cell (%) - 5 min pulse, 0 min chase | | | | |
| **Statistical test:** | Ordinary one-way ANOVA with Tukey's multiple comparisons test | | | | |
| **Simplified genotype** | **N** | **Mean** | **Standard deviation** | **P-value** | |
| | | | | **Compared to Luc Ri** | **Compared to Ykt6 Ri** |
| Luc Ri | 15 | 5,101 | 1,061 | - | <0,0001 |
| Syx7 Ri | 15 | 0 | 0 | <0,0001 | >0,9999 |
| Snap29 Ri | 15 | 0 | 0 | <0,0001 | >0,9999 |
| Ykt6 Ri | 15 | 0 | 0 | <0,0001 | - |
| **Measured:** | Dextran tracer coverage of the cell (%) - 5 min pulse, 0 min chase | | | | |
| **Statistical test:** | Ordinary one-way ANOVA with Tukey's multiple comparisons test | | | | |
| **Simplified genotype** | **N** | **Mean** | **Standard deviation** | **P-value** | |
| | | | | **Compared to Luc Ri** | **Compared to Ykt6 Ri** |
| Luc Ri | 15 | 5,58 | 1,032 | - | <0,0001 |
| Syx7 Ri | 15 | 3,094 | 0,8097 | <0,0001 | 0,6828 |
| Snap29 Ri | 15 | 3,293 | 0,9987 | <0,0001 | 0,3408 |
| Ykt6 Ri | 15 | 2,706 | 0,9545 | <0,0001 | - |
| **Panel P** | | | | | |
| **Measured:** | FITC-Avidin tracer coverage of the cell (%) - 30 min pulse, 0 min chase | | | | |
| **Statistical test:** | Ordinary one-way ANOVA with Tukey's multiple comparisons test | | | | |
| **Simplified genotype** | **N** | **Mean** | **Standard deviation** | **P-value** | |
| | | | | **Compared to Luc Ri** | **Compared to Ykt6 Ri** |
| Luc Ri | 15 | 11,15 | 2,931 | - | <0,0001 |
| Syx7 Ri | 15 | 0 | 0 | <0,0001 | <0,0001 |
| Snap29 Ri | 15 | 0 | 0 | <0,0001 | <0,0001 |
| Ykt6 Ri | 15 | 3,293 | 1,313 | <0,0001 | - |
| **Measured:** | Dextran tracer coverage of the cell (%) - 30 min pulse, 0 min chase | | | | |
| **Statistical test:** | Ordinary one-way ANOVA with Tukey's multiple comparisons test | | | | |
| **Simplified genotype** | **N** | **Mean** | **Standard deviation** | **P-value** | |
| | | | | **Compared to Luc Ri** | **Compared to Ykt6 Ri** |
| Luc Ri | 15 | 11,57 | 2,906 | - | <0,0001 |

| | | | | | |
|---|---|---|---|---|---|
| Syx7 Ri | 15 | 3,559 | 1,172 | <0,0001 | 0,9205 |
| Snap29 Ri | 15 | 3,674 | 1,189 | <0,0001 | 0,9669 |
| Ykt6 Ri | 15 | 3,982 | 1,456 | <0,0001 | - |
| **Panel U** | | | | | |
| **Measured:** | FITC-Avidin tracer coverage of the cell (%) - 30 min pulse, 10 min chase | | | | |
| **Statistical test:** | Ordinary one-way ANOVA with Holm-Šídák's multiple comparisons test | | | | |
| **Simplified genotype** | **N** | **Mean** | **Standard deviation** | **P-value** | |
| | | | | **Compared to Luc Ri** | **Compared to Ykt6 Ri** |
| Luc Ri | 15 | 11,14 | 2,548 | - | <0,0001 |
| Syx7 Ri | 15 | 0 | 0 | <0,0001 | <0,0001 |
| Snap29 Ri | 15 | 0 | 0 | <0,0001 | <0,0001 |
| Ykt6 Ri | 15 | 3,566 | 1,363 | <0,0001 | - |
| **Measured:** | Dextran tracer coverage of the cell (%) - 30 min pulse, 10 min chase | | | | |
| **Statistical test:** | Ordinary one-way ANOVA with Tukey's multiple comparisons test | | | | |
| **Simplified genotype** | **N** | **Mean** | **Standard deviation** | **P-value** | |
| | | | | **Compared to Luc Ri** | **Compared to Ykt6 Ri** |
| Luc Ri | 15 | 11,24 | 2,623 | - | <0,0001 |
| Syx7 Ri | 15 | 0,1201 | 0,1838 | <0,0001 | <0,0001 |
| Snap29 Ri | 15 | 0,2081 | 0,2944 | <0,0001 | <0,0001 |
| Ykt6 Ri | 15 | 3,875 | 1,323 | <0,0001 | - |
| **Figure 3** | | | | | |
| **Panel G** | | | | | |
| **Measured:** | Threshold Overlap Score (TOS) per cell | | | | |
| **Statistical test:** | Unpaired T-test | | | | |
| **Simplified genotype** | **N** | **Mean** | **Standard deviation** | **P-value** | |
| Rab5-GFP:a-Syx7 | 12 | 0,6019 | 0,06971 | <0,0001 | |
| a-Rab7:a-Syx7 | 12 | 0,0434 | 0,5778 | | |
| **Measured:** | Threshold Overlap Score (TOS) per cell | | | | |
| **Statistical test:** | Unpaired T-test | | | | |
| **Simplified genotype** | **N** | **Mean** | **Standard deviation** | **P-value** | |
| Rab5-GFP:a-Snap29 | 12 | 0,6287 | 0,05781 | <0,0001 | |
| a-Rab7:a-Snap29 | 12 | 0,00778 | 0,0728 | | |
| **Measured:** | Threshold Overlap Score (TOS) per cell | | | | |
| **Statistical test:** | Unpaired T-test | | | | |
| **Simplified genotype** | **N** | **Mean** | **Standard deviation** | **P-value** | |

| | | | | | |
|---|---|---|---|---|---|
| Rab5-GFP:Ykt6-HA | 12 | 0,5532 | 0,04995 | <0,0001 | |
| a-Rab7:Ykt6-HA | 12 | 0,03036 | 0,05485 | | |
| **Measured:** | Threshold Overlap Score (TOS) per cell | | | | |
| **Simplified genotype** | **N** | **Mean** | **Standard deviation** | - | |
| a-Syx7:a-Snap29 | 12 | 0,7195 | 0,1196 | | |
| Ykt6-HA:a-Snap29 | 12 | 0,5002 | 0,06497 | | |
| **Figure 4** | | | | | |
| **Panel N** | | | | | |
| **Measured:** | Diameter of a-Rab7+ endosomes (micrometer) | | | | |
| **Statistical test:** | Kruskal-Wallis test with Dunn's multiple comparisons test | | | | |
| **Simplified genotype** | **N** | **Mean** | **Standard deviation** | **P-value** | |
| | | | | **Compared to Luc Ri** | **Compared to Syx7 Ri** |
| Luc Ri | 100 | 1,375 | 0,3193 | - | <0,0001 |
| hSTX7-HA | 100 | 1,406 | 0,3387 | >0,9999 | <0,0001 |
| hSTX12-HA | 100 | 1,454 | 0,364 | >0,9999 | <0,0001 |
| Syx7 Ri | 100 | 0,5926 | 0,719 | <0,0001 | - |
| hSTX7-HA, Syx7 Ri | 100 | 0,7663 | 0,6624 | <0,0001 | 0,4201 |
| hSTX12-HA, Syx7 Ri | 100 | 1,318 | 0,3813 | >0,9999 | <0,0001 |
| **Measured:** | Diameter of a-Rab7+ endosomes (micrometer) | | | | |
| **Statistical test:** | Kruskal-Wallis test with Dunn's multiple comparisons test | | | | |
| **Simplified genotype** | **N** | **Mean** | **Standard deviation** | **P-value** | |
| | | | | **Compared to Luc Ri** | **Compared to Syx13 Ri** |
| Luc Ri | 100 | 1,375 | 0,3193 | - | <0,0001 |
| hSTX7-HA | 100 | 1,406 | 0,3387 | >0,9999 | <0,0001 |
| hSTX12-HA | 100 | 1,454 | 0,364 | >0,9999 | <0,0001 |
| Syx13 Ri | 100 | 2,275 | 0,6173 | <0,0001 | - |
| hSTX7-HA, Syx13 Ri | 100 | 1,37 | 0,3904 | >0,9999 | <0,0001 |
| hSTX12-HA, Syx13 Ri | 100 | 2,19 | 0,7786 | <0,0001 | >0,9999 |
| **Panel U** | | | | | |
| **Measured:** | FITC-Avidin tracer coverage of the cell (%) | | | | |
| **Statistical test:** | Ordinary one-way ANOVA with Tukey's multiple comparisons test | | | | |
| **Simplified genotype** | **N** | **Mean** | **Standard deviation** | **P-value** | |
| | | | | **Compared to Luc Ri** | **Compared to Syx7 Ri** |
| Luc Ri | 15 | 5,058 | 1,013 | - | <0,0001 |
| hSTX7-HA | 15 | 5,055 | 0,9432 | >0,9999 | <0,0001 |
| hSTX12-HA | 15 | 5,021 | 1,012 | >0,9999 | <0,0001 |
| Syx7 Ri | 15 | 0 | 0 | <0,0001 | - |
| hSTX7-HA, Syx7 Ri | 15 | 0,3497 | 0,3663 | <0,0001 | 0,8644 |
| hSTX12-HA, Syx7 Ri | 15 | 2,367 | 1,086 | <0,0001 | <0,0001 |
| **Figure 5** | | | | | |

| Panel L | | | | | |
|---|---|---|---|---|---|
| **Measured:** | Average lacunar channel depth (micrometer) per cell | | | | |
| **Statistical test:** | Ordinary one-way ANOVA with Tukey's multiple comparisons test | | | | |
| **Simplified genotype** | **N** | **Mean** | **Standard deviation** | **P-value** | |
| | | | | **Compared to Luc Ri** | **Compared to Snap29 Ri** |
| Luc Ri | 15 | 1,803 | 0,2983 | - | <0,0001 |
| Syx12L Ri | 15 | 4,142 | 0,6242 | <0,0001 | 0,9243 |
| Snap29 Ri | 15 | 3,943 | 0,6403 | <0,0001 | - |
| Ykt6 Ri | 15 | 0,5313 | 0,1767 | <0,0001 | <0,0001 |
| Vps8 Ri | 15 | 2,32 | 0,4295 | 0,005 | <0,0001 |
| Rbsn5 Ri | 15 | 3,249 | 0,4676 | <0,0001 | <0,0001 |
| Snap29 Ri, Ykt6 Ri | 15 | 0,5587 | 0,2068 | <0,0001 | <0,0001 |
| Sec5 Ri | 15 | 0,5027 | 0,1078 | <0,0001 | <0,0001 |
| Snap29 Ri, Sec5 Ri | 15 | 0,586 | 0,223 | <0,0001 | <0,0001 |
| Vps11 Ri | 15 | 0,8833 | 0,1205 | <0,0001 | <0,0001 |
| Vps41 Ri | 15 | 0,844 | 0,0678 | <0,0001 | <0,0001 |
| Syx7L Ri | 15 | 1,893 | 0,2111 | >0,9999 | <0,0001 |

#### Figure 7

| Panel G | | | | |
|---|---|---|---|---|
| **Measured:** | Diameter of a-Rab7+ endosomes (micrometer) | | | |
| **Statistical test:** | Kruskal-Wallis test with Dunn's multiple comparisons test | | | |
| **Simplified genotype** | **N** | **Mean** | **Standard deviation** | **P-value** |
| | | | | **Compared to Syx12L Ri** |
| Syx12L Ri (Fig8.) | 100 | 0,7201 | 0,8547 | - |
| Syx12L Ri, Snap29 Ri | 100 | 0,3517 | 0,1081 | <0,0001 |
| Syx12L Ri, Ykt6 Ri | 100 | 0,3558 | 0,111 | 0,0001 |
| Syx12L Ri, Syx7L Ri | 100 | 0,3661 | 0,09536 | 0,0049 |
| Syx12L Ri, Vamp7 Ri | 100 | 0,6296 | 0,4692 | >0,9999 |

| Panel H | | | | |
|---|---|---|---|---|
| **Measured:** | Diameter of a-Rab7+ endosomes (micrometer) | | | |
| **Statistical test:** | Kruskal-Wallis test with Dunn's multiple comparisons test | | | |
| **Simplified genotype** | **N** | **Mean** | **Standard deviation** | **P-value** |
| | | | | **Compared to Ykt6 Ri** |
| Ykt6 Ri (Fig8.) | 100 | 0,6074 | 0,1822 | - |
| Ykt6 Ri, Snap29 Ri | 100 | 0,4029 | 0,1456 | <0,0001 |
| Ykt6 Ri, Syx12L Ri | 100 | 0,3558 | 0,111 | <0,0001 |
| Ykt6 Ri, Vamp7 Ri | 100 | 0,3976 | 0,1265 | <0,0001 |
| Ykt6 Ri, Syx7L Ri | 100 | 0,7416 | 0,2349 | 0,039 |

#### Figure 8

| Panel E | | | | |
|---|---|---|---|---|
| **Measured:** | Diameter of Rab5-YFP+ endosomes (micrometer) | | | |
| **Statistical test:** | Kruskal-Wallis test with Dunn's multiple comparisons test | | | |
| **Simplified genotype** | **N** | **Mean** | | **P-value** |

| | | | Standard deviation | Compared to Luc Ri | Compared to Snap29 Ri |
|---|---|---|---|---|---|
| Luc Ri | 100 | 1,018 | 0,3645 | - | <0,0001 |
| Snap29 Ri | 100 | 0,2564 | 0,07054 | <0,0001 | - |
| Syx12L Ri (Fig7.) | 100 | 0,3481 | 0,1025 | <0,0001 | 0,0009 |
| Ykt6 Ri (Fig7.) | 100 | 0,5726 | 0,1802 | <0,0001 | <0,0001 |
| **Measured:** | Diameter of a-Rab7+ endosomes (micrometer) | | | | |
| **Statistical test:** | Kruskal-Wallis test with Dunn's multiple comparisons test | | | | |
| **Simplified genotype** | **N** | **Mean** | **Standard deviation** | **P-value** | |
| | | | | **Compared to Luc Ri** | **Compared to Snap29 Ri** |
| Luc Ri | 100 | 1,437 | 0,45 | - | <0,0001 |
| Snap29 Ri | 100 | 0,274 | 0,09596 | <0,0001 | - |
| Syx12L Ri (Fig7.) | 100 | 0,7201 | 0,8547 | <0,0001 | <0,0001 |
| Ykt6 Ri (Fig7.) | 100 | 0,6074 | 0,1822 | <0,0001 | <0,0001 |
| **Panel J** | | | | | |
| **Measured:** | Diameter of Rab5CA-YFP+ endosomes (micrometer) | | | | |
| **Statistical test:** | Kruskal-Wallis test with Dunn's multiple comparisons test | | | | |
| **Simplified genotype** | **N** | **Mean** | **Standard deviation** | **P-value** | |
| | | | | **Compared to Luc Ri** | **Compared to Snap29 Ri** |
| Luc Ri | 100 | 3,085 | 2,029 | - | <0,0001 |
| Snap29 Ri | 100 | 0,2645 | 0,08694 | <0,0001 | - |
| Syx12L Ri | 100 | 0,7091 | 0,775 | <0,0001 | <0,0001 |
| Ykt6 Ri | 100 | 1,308 | 0,5974 | <0,0001 | <0,0001 |
| **Measured:** | Diameter of a-Rab7+ endosomes (micrometer) | | | | |
| **Statistical test:** | Kruskal-Wallis test with Dunn's multiple comparisons test | | | | |
| **Simplified genotype** | **N** | **Mean** | **Standard deviation** | **P-value** | |
| | | | | **Compared to Luc Ri** | **Compared to Snap29 Ri** |
| Luc Ri, Rab5CA | 100 | 3,278 | 2,258 | - | <0,0001 |
| Snap29 Ri, Rab5CA | 100 | 0,2774 | 0,06551 | <0,0001 | - |
| Syx12L Ri, Rab5CA | 100 | 0,7444 | 0,8215 | <0,0001 | <0,0001 |
| Ykt6 Ri, Rab5CA | 100 | 1,32 | 0,5459 | <0,0001 | <0,0001 |
| **Panel O** | | | | | |
| **Measured:** | Diameter of Rab5-YFP+ endosomes (micrometer) | | | | |
| **Statistical test:** | Kruskal-Wallis test with Dunn's multiple comparisons test | | | | |
| **Simplified genotype** | **N** | **Mean** | **Standard deviation** | **P-value** | |
| | | | | **Compared to Luc Ri** | **Compared to Snap29 Ri** |
| Rab7 Ri, Luc Ri | 100 | 1,176 | 0,5025 | - | <0,0001 |
| Rab7 Ri, Snap29 Ri | 100 | 0,2485 | 0,08114 | <0,0001 | - |
| Rab7 Ri, Syx12L Ri | 100 | 0,3195 | 0,08927 | <0,0001 | 0,0046 |
| Rab7 Ri, Ykt6 Ri | 100 | 0,4672 | 0,121 | <0,0001 | <0,0001 |
| **Panel T** | | | | | |
| **Measured:** | Diameter of Rab5CA-YFP+ endosomes (micrometer) | | | | |

| | | | | | |
|---|---|---|---|---|---|
| **Statistical test:** | Kruskal-Wallis test with Dunn's multiple comparisons test | | | | |
| **Simplified genotype** | **N** | **Mean** | **Standard deviation** | **P-value** | |
| | | | | **Compared to Luc Ri** | **Compared to Snap29 Ri** |
| Rab7 Ri, Luc Ri | 100 | 2,39 | 1,24 | - | <0,0001 |
| Rab7 Ri, Snap29 Ri | 100 | 0,231 | 0,07911 | <0,0001 | - |
| Rab7 Ri, Syx12L Ri | 100 | 0,3719 | 0,1471 | <0,0001 | 0,0002 |
| Rab7 Ri, Ykt6 Ri | 100 | 0,9054 | 0,4107 | <0,0001 | <0,0001 |
| **Panel U** | | | | | |
| **Measured:** | Diameter of Rab5-YFP/Rab5CA-YFP+ endosomes (micrometer) - Luc Ri | | | | |
| **Statistical test:** | Kruskal-Wallis test with Dunn's multiple comparisons test | | | | |
| **Simplified genotype** | **N** | **Mean** | **Standard deviation** | **P-value** | |
| | | | | **Compared to Rab5-YFP** | **Compared to Rab7 Ri, Rab5-YFP** |
| Rab5-YFP | 100 | 1,018 | 0,3645 | - | 0,4992 |
| Rab5CA-YFP | 100 | 3,085 | 2,029 | <0,0001 | <0,0001 |
| Rab7 Ri, Rab5-YFP | 100 | 1,176 | 0,5025 | 0,4992 | - |
| Rab7 Ri, Rab5CA-YFP | 100 | 2,39 | 1,24 | <0,0001 | <0,0001 |
| **Measured:** | Diameter of Rab7+ endosomes (micrometer) - Luc Ri | | | | |
| **Statistical test:** | Mann-Whitney test | | | | |
| **Simplified genotype** | **N** | **Mean** | **Standard deviation** | **P-value** | |
| Rab5-YFP | 100 | 1,437 | 0,45 | <0,0001 | |
| Rab5CA-YFP | 100 | 3,278 | 2,258 | | |
| **Panel V** | | | | | |
| **Measured:** | Diameter of Rab5-YFP/Rab5CA-YFP+ endosomes (micrometer) - Snap29 Ri | | | | |
| **Statistical test:** | Kruskal-Wallis test with Dunn's multiple comparisons test | | | | |
| **Simplified genotype** | **N** | **Mean** | **Standard deviation** | **P-value** | |
| | | | | **Compared to Rab5-YFP** | **Compared to Rab7 Ri, Rab5-YFP** |
| Rab5-YFP | 100 | 0,2564 | 0,07054 | - | >0,9999 |
| Rab5CA-YFP | 100 | 0,2645 | 0,08694 | >0,9999 | >0,9999 |
| Rab7 Ri, Rab5-YFP | 100 | 0,2485 | 0,08114 | >0,9999 | - |
| Rab7 Ri, Rab5CA-YFP | 100 | 0,231 | 0,07911 | 0,1409 | >0,9999 |
| **Measured:** | Diameter of Rab7+ endosomes (micrometer) - Snap29 Ri | | | | |
| **Statistical test:** | Mann-Whitney test | | | | |
| **Simplified genotype** | **N** | **Mean** | **Standard deviation** | **P-value** | |
| Rab5-YFP | 100 | 0,274 | 0,09596 | 0,5098 | |

<table>
<tr><td>Rab5CA-YFP</td><td>100</td><td>0,2774</td><td>0,06551</td><td colspan="2"></td></tr>
<tr><td colspan="6">Panel W</td></tr>
<tr><td>Measured:</td><td colspan="5">Diameter of Rab5-YFP/Rab5CA-YFP+ endosomes (micrometer) - Syx12L Ri</td></tr>
<tr><td>Statistical test:</td><td colspan="5">Kruskal-Wallis test with Dunn's multiple comparisons test</td></tr>
<tr><td rowspan="2">Simplified genotype</td><td rowspan="2">N</td><td rowspan="2">Mean</td><td rowspan="2">Standard deviation</td><td colspan="2">P-value</td></tr>
<tr><td>Compared to Rab5-YFP</td><td>Compared to Rab7 Ri, Rab5-YFP</td></tr>
<tr><td>Rab5-YFP (Fig7.)</td><td>100</td><td>0,3481</td><td>0,1025</td><td>-</td><td>0,4074</td></tr>
<tr><td>Rab5CA-YFP</td><td>100</td><td>0,7091</td><td>0,775</td><td>0,0004</td><td><0,0001</td></tr>
<tr><td>Rab7 Ri, Rab5-YFP</td><td>100</td><td>0,3195</td><td>0,08927</td><td>0,4074</td><td>-</td></tr>
<tr><td>Rab7 Ri, Rab5CA-YFP</td><td>100</td><td>0,3719</td><td>0,1471</td><td>>0,9999</td><td>0,0868</td></tr>
<tr><td>Measured:</td><td colspan="5">Diameter of Rab7+ endosomes (micrometer) - Syx12L Ri</td></tr>
<tr><td>Statistical test:</td><td colspan="5">Mann-Whitney test</td></tr>
<tr><td>Simplified genotype</td><td>N</td><td>Mean</td><td>Standard deviation</td><td colspan="2">P-value</td></tr>
<tr><td>Rab5-YFP (Fig7.)</td><td>100</td><td>0,7201</td><td>0,8547</td><td colspan="2" rowspan="2">0,843</td></tr>
<tr><td>Rab5CA-YFP</td><td>100</td><td>0,7444</td><td>0,8215</td></tr>
<tr><td colspan="6">Panel X</td></tr>
<tr><td>Measured:</td><td colspan="5">Diameter of Rab5-YFP/Rab5CA-YFP+ endosomes (micrometer) - Ykt6 Ri</td></tr>
<tr><td>Statistical test:</td><td colspan="5">Kruskal-Wallis test with Dunn's multiple comparisons test</td></tr>
<tr><td rowspan="2">Simplified genotype</td><td rowspan="2">N</td><td rowspan="2">Mean</td><td rowspan="2">Standard deviation</td><td colspan="2">P-value</td></tr>
<tr><td>Compared to Rab5-YFP</td><td>Compared to Rab7 Ri, Rab5-YFP</td></tr>
<tr><td>Rab5-YFP (Fig7.)</td><td>100</td><td>0,5726</td><td>0,1802</td><td>-</td><td>0,0166</td></tr>
<tr><td>Rab5CA-YFP</td><td>100</td><td>1,308</td><td>0,5974</td><td><0,0001</td><td><0,0001</td></tr>
<tr><td>Rab7 Ri, Rab5-YFP</td><td>100</td><td>0,4672</td><td>0,121</td><td>0,0166</td><td>-</td></tr>
<tr><td>Rab7 Ri, Rab5CA-YFP</td><td>100</td><td>0,9054</td><td>0,4107</td><td><0,0001</td><td><0,0001</td></tr>
<tr><td>Measured:</td><td colspan="5">Diameter of Rab7+ endosomes (micrometer) - Ykt6 Ri</td></tr>
<tr><td>Statistical test:</td><td colspan="5">Mann-Whitney test</td></tr>
<tr><td>Simplified genotype</td><td>N</td><td>Mean</td><td>Standard deviation</td><td colspan="2">P-value</td></tr>
<tr><td>Rab5-YFP (Fig7.)</td><td>100</td><td>0,6074</td><td>0,1822</td><td colspan="2" rowspan="2"><0,0001</td></tr>
<tr><td>Rab5CA-YFP</td><td>100</td><td>1,32</td><td>0,5459</td></tr>
<tr><td colspan="6">Supplementary Figure 2.</td></tr>
<tr><td colspan="6">Panel I.</td></tr>
<tr><td>Measured:</td><td colspan="5">Diameter of a-Rab7+ endosomes (micrometer)</td></tr>
<tr><td>Statistical test:</td><td colspan="5">Kruskal-Wallis test with Dunn's multiple comparisons test</td></tr>
<tr><td>Simplified genotype</td><td>N</td><td>Mean</td><td></td><td colspan="2">P-value</td></tr>
</table>

| | | | Standard deviation | Compared to Luc Ri |
|---|---|---|---|---|
| Luc Ri | 100 | 1,338 | 0,3757 | - |
| Syx1 R1 | 100 | 1,973 | 0,5894 | <0,0001 |
| Syx1 R2 | 100 | 1,775 | 0,5086 | <0,0001 |
| Syx4 R1 | 100 | 1,356 | 0,3937 | >0,9999 |
| Syx4 R2 | 100 | 1,592 | 0,4703 | 0,1415 |
| Syx4 R3 | 100 | 1,892 | 0,5892 | <0,0001 |
| Syx4 R4 | 100 | 1,357 | 0,3408 | >0,9999 |
| Syx5 R1 | 100 | 2,205 | 0,7461 | <0,0001 |
| Syx5 R2 | 100 | 1,48 | 0,9879 | >0,9999 |
| Syx7 R1 | 100 | 0,7765 | 0,8261 | <0,0001 |
| Syx7 R2 | 100 | 0,669 | 0,7206 | <0,0001 |
| Syx7 R3 | 100 | 0,609 | 0,6186 | <0,0001 |
| Syx7 R4 | 100 | 0,5989 | 0,5868 | <0,0001 |
| Syx13 R1 | 100 | 2,265 | 0,8622 | <0,0001 |
| Syx13 R2 | 100 | 1,37 | 0,3791 | >0,9999 |
| Syx13 R3 | 100 | 2,03 | 0,8136 | <0,0001 |
| Syx13 R4 | 100 | 2,016 | 0,7496 | <0,0001 |
| Syx16 R1 | 100 | 1,601 | 0,4649 | 0,064 |
| Syx16 R2 | 100 | 1,427 | 0,4098 | >0,9999 |
| Syx16 R3 | 100 | 1,825 | 0,4351 | <0,0001 |
| Syx16 R4 | 100 | 1,643 | 0,3989 | 0,0037 |
| Syx18 R1 | 100 | 1,752 | 0,5761 | 0,0003 |
| Syx18 R2 | 100 | 1,982 | 0,5886 | <0,0001 |
| Syx18 R3 | 100 | 1,613 | 0,5605 | 0,1152 |
| Syx18 R4 | 100 | 1,252 | 0,3604 | >0,9999 |
| Gos28 R1 | 100 | 1,292 | 0,371 | >0,9999 |
| Gos28 R2 | 100 | 1,304 | 0,3482 | >0,9999 |
| Gos28 R3 | 100 | 1,427 | 0,338 | >0,9999 |
| Membrin R1 | 100 | 1,932 | 0,465 | <0,0001 |
| Membrin R2 | 100 | 1,791 | 0,4593 | <0,0001 |
| Membrin R3 | 100 | 1,329 | 0,4229 | >0,9999 |
| Membrin R4 | 100 | 1,424 | 0,3623 | >0,9999 |
| Sec20 R1 | 100 | 2,123 | 0,7819 | <0,0001 |
| Sec20 R2 | 100 | 2,118 | 0,6514 | <0,0001 |
| Vti1a R1 | 100 | 1,474 | 0,4459 | >0,9999 |
| Vti1a R2 | 100 | 1,438 | 0,3905 | >0,9999 |
| Bet1 R1 | 100 | 1,456 | 0,4011 | >0,9999 |
| Bet1 R2 | 100 | 1,36 | 0,3461 | >0,9999 |
| Syx6 R1 | 100 | 1,285 | 0,3784 | >0,9999 |
| Syx6 R2 | 100 | 1,438 | 0,3966 | >0,9999 |
| Syx6 R3 | 100 | 1,356 | 0,3552 | >0,9999 |
| Syx8 R1 | 100 | 1,734 | 0,6424 | 0,0044 |
| Syx8 R2 | 100 | 1,316 | 0,362 | >0,9999 |
| Syx8 R3 | 100 | 1,285 | 0,5466 | >0,9999 |
| Use1 R1 | 100 | 1,061 | 0,2813 | 0,0138 |

| | | | | |
|---|---|---|---|---|
| Use1 R2 | 100 | 1,347 | 0,382 | >0,9999 |
| Use1 R3 | 100 | 1,318 | 0,5204 | >0,9999 |
| Snap24 R1 | 100 | 1,958 | 0,6426 | <0,0001 |
| Snap24 R2 | 100 | 1,437 | 0,3481 | >0,9999 |
| Snap24 R3 | 100 | 1,51 | 0,3307 | >0,9999 |
| Snap25 R1 | 100 | 1,421 | 0,4199 | >0,9999 |
| Snap25 R2 | 100 | 1,373 | 0,3462 | >0,9999 |
| Snap29 R1 | 100 | 0,2977 | 0,08781 | <0,0001 |
| Snap29 R2 | 100 | 0,3212 | 0,09467 | <0,0001 |
| Snap29 R3 | 100 | 1,375 | 0,4512 | >0,9999 |
| Snap29 R4 | 100 | 0,3645 | 0,1231 | <0,0001 |
| Sec22 R1 | 100 | 1,583 | 0,4625 | 0,1388 |
| Sec22 R2 | 100 | 1,86 | 0,5901 | <0,0001 |
| Sec22 R3 | 100 | 1,564 | 0,4788 | 0,435 |
| Syb R1 | 100 | 2,208 | 0,7313 | <0,0001 |
| Syb R2 | 100 | 1,267 | 0,3795 | >0,9999 |
| Syb R3 | 100 | 1,265 | 0,3747 | >0,9999 |
| Syb R4 | 100 | 2,21 | 0,91 | <0,0001 |
| Syb R5 | 100 | 1,216 | 0,3489 | >0,9999 |
| nSyb R1 | 100 | 1,397 | 0,4094 | >0,9999 |
| nSyb R2 | 100 | 1,562 | 0,4649 | 0,3435 |
| nSyb R3 | 100 | 1,396 | 0,4639 | >0,9999 |
| nSyb R4 | 100 | 1,433 | 0,4397 | >0,9999 |
| Vamp7 R1 | 100 | 1,853 | 0,5466 | <0,0001 |
| Vamp7 R2 | 100 | 1,682 | 0,4909 | 0,0033 |
| Vamp7 R3 | 100 | 1,945 | 0,5616 | <0,0001 |
| Vamp7 R4 | 100 | 1,615 | 0,4691 | 0,0526 |
| Vamp7 R5 | 100 | 1,969 | 0,5797 | <0,0001 |
| Ykt6 R1 | 100 | 0,5926 | 0,1339 | <0,0001 |
| Ykt6 R2 | 100 | 0,6512 | 0,1456 | <0,0001 |
| Ykt6 R3 | 100 | 1,445 | 0,438 | >0,9999 |
| Ykt6 R4 | 100 | 0,784 | 0,159 | <0,0001 |
| Ykt6 R5 | 100 | 0,7143 | 0,1585 | <0,0001 |

#### Supplementary Figure 4.

##### Panel J.

| | | | | |
|---|---|---|---|---|
| **Measured:** | Surface density of a-Pyd (Number/10 micrometers) per cell | | | |
| **Statistical test:** | Ordinary one-way ANOVA with Dunnett's multiple comparisons test | | | |
| **Simplified genotype** | **N** | **Mean** | **Standard deviation** | **P-value** |
| | | | | **Compared to Luc Ri** |
| Luc Ri | 12 | 15,6 | 1,527 | - |
| Syx12L Ri | 12 | 9,202 | 0,7386 | <0,0001 |
| Snap29 Ri | 12 | 15,51 | 1,445 | 0,9954 |
| Ykt6 Ri | 12 | 11,21 | 1,036 | <0,0001 |
| **Measured:** | Surface density of a-Sns (Number/10 micrometers) per cell | | | |

| Statistical test: | Ordinary one-way ANOVA with Dunnett's multiple comparisons test | | | |
|---|---|---|---|---|
| **Simplified genotype** | **N** | **Mean** | **Standard deviation** | **P-value Compared to Luc Ri** |
| Luc Ri | 12 | 15,99 | 1,591 | - |
| Syx12L Ri | 12 | 9,042 | 1,646 | <0,0001 |
| Snap29 Ri | 12 | 15,51 | 1,327 | 0,7863 |
| Ykt6 Ri | 12 | 11,8 | 1,613 | <0,0001 |
| **Panel K.** | | | | |
| **Measured:** | Threshold Overlap Score (TOS) between a-Pyd and a-Sns per cell (Cortical) | | | |
| **Statistical test:** | Kruskal-Wallis test with Dunn's multiple comparisons test | | | |
| **Simplified genotype** | **N** | **Mean** | **Standard deviation** | **P-value Compared to Luc Ri** |
| Luc Ri | 12 | 0,5975 | 0,04444 | - |
| Syx12L Ri | 12 | 0,4261 | 0,05846 | 0,0438 |
| Snap29 Ri | 12 | 0,6239 | 0,0493 | >0,9999 |
| Ykt6 Ri | 12 | 0,04421 | 0,03758 | <0,0001 |
| **Panel T.** | | | | |
| **Measured:** | Maximum signal depth of a-Pyd (micrometer) per cell (Medial) | | | |
| **Statistical test:** | Ordinary one-way ANOVA with Dunnett's multiple comparisons test | | | |
| **Simplified genotype** | **N** | **Mean** | **Standard deviation** | **P-value Compared to Luc Ri** |
| Luc Ri | 12 | 0,5928 | 0,1492 | - |
| Syx12L Ri | 12 | 4,937 | 1,665 | <0,0001 |
| Snap29 Ri | 12 | 4,402 | 0,8022 | <0,0001 |
| Ykt6 Ri | 12 | 0,5471 | 0,1209 | 0,9987 |
| **Measured:** | Maximum signal depth of a-Sns (micrometer) per cell (Medial) | | | |
| **Statistical test:** | Ordinary one-way ANOVA with Dunnett's multiple comparisons test | | | |
| **Simplified genotype** | **N** | **Mean** | **Standard deviation** | **P-value Compared to Luc Ri** |
| Luc Ri | 12 | 0,5928 | 0,1492 | - |
| Syx12L Ri | 12 | 4,804 | 1,644 | <0,0001 |
| Snap29 Ri | 12 | 4,018 | 1,189 | <0,0001 |
| Ykt6 Ri | 12 | 0,5471 | 0,1209 | 0,999 |
| **Supplementary Figure 5.** | | | | |
| **Panel H.** | | | | |
| **Measured:** | Diameter of Rab5-GFP+ endosomes (micrometer) | | | |
| **Statistical test:** | Kruskal-Wallis test with Dunn's multiple comparisons test | | | |
| **Simplified genotype** | **N** | **Mean** | **Standard deviation** | **P-value Compared to Syx12L Ri** |
| Syx12L Ri (Fig8.) | 100 | 0,3481 | 0,1025 | - |
| Syx12L Ri, Snap29 Ri | 100 | 0,3235 | 0,08772 | 0,2132 |
| Syx12L Ri, Ykt6 Ri | 100 | 0,3559 | 0,1034 | >0,9999 |

| | | | | |
|---|---|---|---|---|
| Syx12L Ri, Syx7L Ri | 100 | 0,3882 | 0,1032 | 0,0534 |
| Syx12L Ri, Vamp7 Ri | 100 | 0,5914 | 0,4413 | <0,0001 |
| **Panel I.** | | | | |
| **Measured:** | Diameter of Rab5-GFP+ endosomes (micrometer) | | | |
| **Statistical test:** | Kruskal-Wallis test with Dunn's multiple comparisons test | | | |
| **Simplified genotype** | **N** | **Mean** | **Standard deviation** | **P-value** |
| | | | | **Compared to Ykt6 Ri** |
| Ykt6 Ri (Fig8.) | 100 | 0,5726 | 0,1802 | - |
| Ykt6 Ri, Snap29 Ri | 100 | 0,4272 | 0,1388 | <0,0001 |
| Ykt6 Ri, Syx12L Ri | 100 | 0,3559 | 0,1034 | <0,0001 |
| Ykt6 Ri, Vamp7 Ri | 100 | 0,3781 | 0,1457 | <0,0001 |
| Ykt6 Ri, Syx7L Ri | 100 | 0,7096 | 0,2097 | 0,002 |
| **Panel J.** | | | | |
| **Measured:** | Diameter of Rab5-GFP+ endosomes (micrometer) | | | |
| **Statistical test:** | Kruskal-Wallis test with Dunn's multiple comparisons test | | | |
| **Simplified genotype** | **N** | **Mean** | **Standard deviation** | **P-value** |
| | | | | **Compared to Vamp7 Ri, Syx7L Ri** |
| Vamp7 Ri | 100 | 0,9451 | 0,3387 | 0,0002 |
| Syx7L Ri | 100 | 0,8991 | 0,509 | <0,0001 |
| Vamp7 Ri, Syx7L Ri | 100 | 1,24 | 0,5243 | - |
| **Panel K.** | | | | |
| **Measured:** | Diameter of a-Rab7+ endosomes (micrometer) | | | |
| **Statistical test:** | Kruskal-Wallis test with Dunn's multiple comparisons test | | | |
| **Simplified genotype** | **N** | **Mean** | **Standard deviation** | **P-value** |
| | | | | **Compared to Vamp7 Ri, Syx7L Ri** |
| Vamp7 Ri | 100 | 1,89 | 0,5317 | 0,1653 |
| Syx7L Ri | 100 | 1,925 | 0,6299 | 0,228 |
| Vamp7 Ri, Syx7L Ri | 100 | 2,205 | 0,9526 | - |

#### Supplemental references


1. S. Jean, S. Cox, S. Nassari, A. A. Kiger, Starvation-induced MTMR13 and RAB21 activity regulates VAMP8 to promote autophagosome–lysosome fusion. *EMBO reports* **16**, 297-311 (2015).
2. V. K. Lund, K. L. Madsen, O. Kjaerulff, Drosophila Rab2 controls endosome-lysosome fusion and LAMP delivery to late endosomes. *Autophagy* **14**, 1520-1542 (2018).
3. Y. Lu, Z. Zhang, D. Sun, S. T. Sweeney, F. B. Gao, Syntaxin 13, a genetic modifier of mutant CHMP2B in frontotemporal dementia, is required for autophagosome maturation. *Molecular cell* **52**, 264-271 (2013).
4. S. Takats, G. Glatz, G. Szenci, A. Boda, G. V. Horvath, K. Hegedus, A. L. Kovacs, G. Juhasz, Non-canonical role of the SNARE protein Ykt6 in autophagosome-lysosome fusion. *PLoS genetics* **14**, e1007359 (2018).
5. P. Lorincz, Z. Lakatos, A. Varga, T. Maruzs, Z. Simon-Vecsei, Z. Darula, P. Benko, G. Csordas, M. Lippai, I. Ando, K. Hegedus, K. F. Medzihradszky, S. Takats, G. Juhasz, MiniCORVET is a Vps8-containing early endosomal tether in Drosophila. *eLife* **5**, e14226 (2016).
6. A. Atienza-Manuel, V. Castillo-Mancho, S. De Renzis, J. Culi, M. Ruiz-Gómez, Endocytosis mediated by an atypical CUBAM complex modulates slit diaphragm dynamics in nephrocytes. *Development (Cambridge, England)* **148**,  (2021).
7. S. Takats, P. Nagy, A. Varga, K. Pircs, M. Karpati, K. Varga, A. L. Kovacs, K. Hegedus, G. Juhasz, Autophagosomal Syntaxin17-dependent lysosomal degradation maintains neuronal function in Drosophila. *The Journal of cell biology* **201**, 531-539 (2013).
8. N. Chaudhry, M. Sica, S. Surabhi, D. S. Hernandez, A. Mesquita, A. Selimovic, A. Riaz, L. Lescat, H. Bai, G. C. MacIntosh, A. Jenny, Lamp1 mediates lipid transport, but is dispensable for autophagy in Drosophila. *Autophagy* **18**, 2443-2458 (2022).
9. F. Hochapfel, L. Denk, G. Mendl, U. Schulze, C. Maaßen, Y. Zaytseva, H. Pavenstädt, T. Weide, R. Rachel, R. Witzgall, Distinct functions of Crumbs regulating slit diaphragms and endocytosis in Drosophila nephrocytes. *Cellular and Molecular Life Sciences* **74**, 4573-4586 (2017).
10. H. Lu, D. Bilder, Endocytic control of epithelial polarity and proliferation in Drosophila. *Nature cell biology* **7**, 1232-1239 (2005).
11. G. Glatz, G. Gogl, A. Alexa, A. Remenyi, Structural mechanism for the specific assembly and activation of the extracellular signal regulated kinase 5 (ERK5) module. *The Journal of biological chemistry* **288**, 8596-8609 (2013).
